# A Medicinal Chemistry-Centered Evaluation of AlphaFold 3 and Boltz-2 Across Diverse Binding Modalities

**DOI:** 10.64898/2026.08.24.746785

**Authors:** Ke Chen, Zuohuang Qi, Omar Lozano Ramos, Huayu Li, Mengjiao Ma, Malla Reddy Gannarapu, Fangchao Bi, Ao Li, Hongmin Li, Rui Xiong

## Abstract

AlphaFold 3 (AF3) and Boltz-2 are state-of-the-art AI-based tools for biomolecular structure prediction, but whether their predictions provide useful guidance for lead optimization, SAR interpretation, and virtual screening remains insufficiently characterized. We benchmarked their performance using newly determined soluble epoxide hydrolase co-crystal structures and matched activity data together with a curated post-training-cutoff dataset spanning kinases, allosteric modulators, covalent systems, PROTACs, molecular glues, fragments, membrane proteins, RNA binders, and activity-cliff pairs. Both models recovered canonical orthosteric enzyme and kinase complexes, including key DFG/αC conformational states, whereas allosteric, membrane-protein, and induced-proximity complexes remained challenging. Pharmacophore RMSD was often lower than overall ligand RMSD, indicating preservation of key recognition features despite imperfect whole-ligand alignment. AF3 minPAE correlated with pose accuracy, and very low minPAE values (<0.85 Å) were strongly enriched for accurate poses. Model confidence scores were not associated with experimental activity, whereas Boltz-2 predicted affinity captured relative activity trends and distinguished the activity-cliff pair, although its performance varied across ligand series.

## Introduction

AlphaFold 3^1^ (AF3) and Boltz-2^2^ represent a new generation of AI-based tools for biomolecular structure prediction. Traditional structure-based drug design often relies on resource-intensive experimental structural biology paired with physics-based docking, molecular dynamics, or free-energy perturbation (FEP),^3–6^ workflows that can be bottlenecked by the need for high-resolution starting structures, extensive system preparation, adequate conformational sampling, and force-field parameterization.^4,7,8^ AF3 and Boltz-2 offer a potentially transformative alternative by performing joint complex prediction, or co-folding, in which proteins, ligands, nucleic acids, ions, and other molecular components are modeled together within a unified diffusion-based framework. Their architectures use learned representations and attention-based interaction modeling to capture residue–residue, residue–ligand, and interfacial interaction patterns during complex generation. In addition to predicted structures, these models provide confidence metrics such as predicted TM-score for interfaces (ipTM) and predicted aligned error (PAE), which may help estimate interface-level and pairwise uncertainty.^1,2^ Boltz-2 further incorporates an affinity-prediction module, enabling simultaneous evaluation of predicted binding poses and ligand-binding strength.^2^ Accordingly, these capabilities have generated substantial interest in using AF3 and Boltz-2 for drug discovery.^1,2^

For medicinal chemistry, however, the more relevant question is not simply whether a model reproduces the experimental ligand pose, but whether the predicted structure preserves the interactions needed to interpret SAR and guide compound design. Existing evaluations of protein–ligand prediction models often emphasize global ligand RMSD or broad pose recovery,^9,10^ but medicinal chemistry applications require a more nuanced assessment of local pharmacophore preservation^11^, ligand-induced conformational recovery,^12^ affinity ranking,^13^ and whether AI-generated confidence metrics such as ipTM or minPAE can be used to triage predictions. This gap is particularly important for challenging noncanonical recognition modes involving induced fit, shallow or cryptic pockets, induced proximity, protein–protein interaction sites, and fragment-sized ligands.^12,14,15^

Here, we provide a medicinal chemistry-centered evaluation of AF3 and Boltz-2 across diverse ligand-binding modalities. Our benchmark combines newly determined in-house soluble epoxide hydrolase co-crystal structures and matched activity data with a curated post-training-cutoff dataset covering kinases, allosteric modulators, covalent systems, PROTACs, molecular glues, fragments, membrane proteins, RNA binders, and activity-cliff pairs. We asked practical questions relevant to structure-guided design: which modalities yield ligand poses within 2 Å RMSD; whether local pharmacophore features are preserved more accurately than global ligand poses; whether confidence metrics such as ipTM or minPAE correlate with pose accuracy and can identify accurate ligand poses; and whether Boltz-2 affinity estimates can support activity ranking, activity-cliff differentiation, and virtual-screening prioritization. We also assessed limitations that can affect medicinal chemistry interpretation, including chirality errors and bond-order or bond-length deviations. Together, this work defines modality-dependent strengths and limitations of AF3 and Boltz-2 and provides a practical framework for their use in pose prediction, virtual screening, and SAR interpretation.

## Results

To assess the practical utility of AF3 and Boltz-2 in medicinal chemistry, we evaluated their performance across representative drugging modalities (**Figure 1A**) and examined how model performance varies with conserved pharmacophores and modality-specific recognition challenges, including induced fit, shallow-surface binding, cryptic-pocket formation, ternary-complex cooperativity, covalent-ligand positioning, and activity cliffs. Because a single geometric metric cannot fully capture whether a predicted pose is useful for medicinal chemistry interpretation, we applied three complementary measures (**Figure 1B**) of pose similarity, with detailed calculations provided in the Methods section.

**Figure 1.**
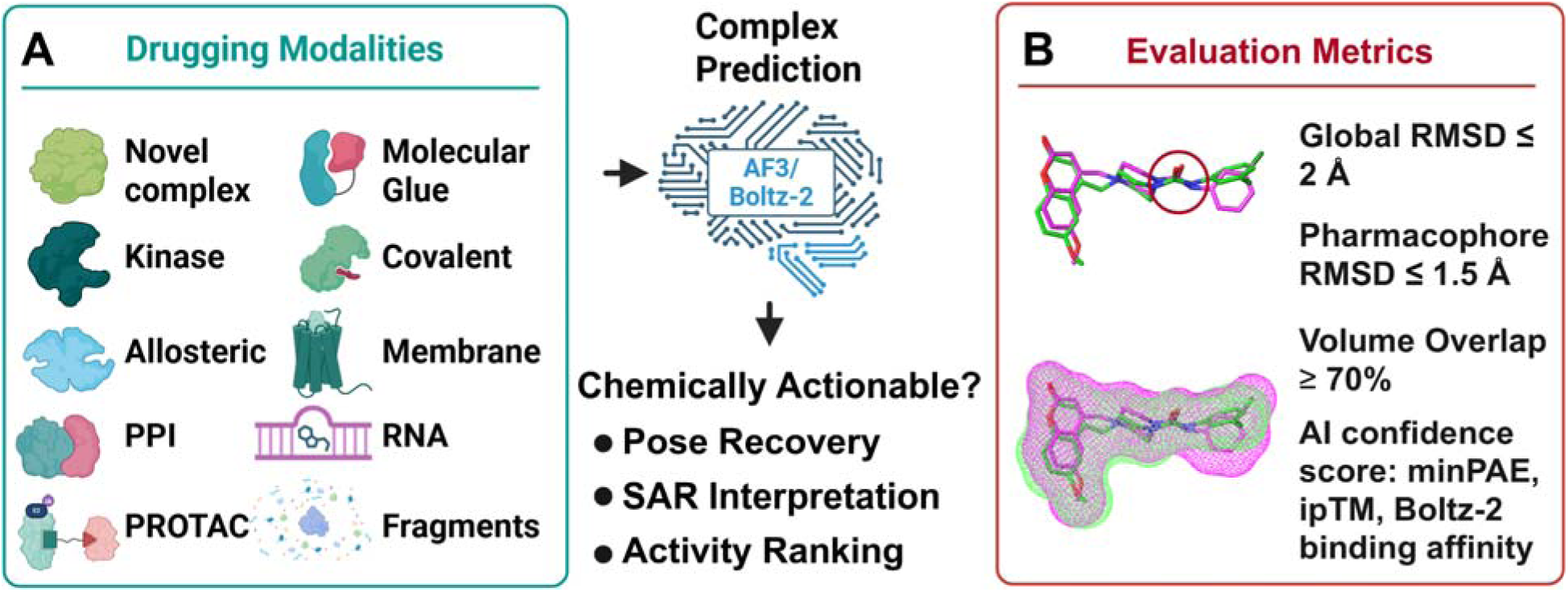
Benchmarking AI-based complex prediction for medicinal chemistry. (**A**) Diverse drug discovery modalities evaluated in this study, including orthosteric enzyme inhibitors (sEH and kinases), allosteric modulators, protein–protein interaction (PPI) inhibitors, PROTACs, molecular glues, covalent inhibitors, membrane protein ligands, RNA-targeting compounds, and fragment binders. Predicted protein–ligand, protein–protein, and protein–RNA complexes were generated using AF3 and Boltz-2. Complex predictions were assessed for their utility in pose recovery, structure–activity relationship (SAR) interpretation, and confidence-based filtering of actionable models. (**B**) Evaluation metrics used to determine prediction quality, including global ligand RMSD (≤ 2 Å), pharmacophore RMSD (≤ 1.5 Å), volume overlap (≥ 70%), and AI confidence metrics such as predicted aligned error (PAE) and interface predicted TM-score (ipTM). Together, these metrics provide a framework for assessing whether AI-generated structures are suitable for downstream medicinal chemistry and structure-guided drug design applications.

First, overall RMSD provides a global measure of positional deviation from the reference pose. It was calculated over all ligand heavy atoms after protein alignment. Second, key-pharmacophore RMSD focuses on interaction-defining ligand features, such as hydrogen-bonding and electrostatic interaction motifs, that were either reported in the literature or observed in reference structure. Third, volume overlap evaluates whether the predicted ligand occupies the same three-dimensional binding space as the experimental pose. This was assessed using RDKit shape-based Tanimoto analysis and is particularly useful in cases where atom-by-atom correspondence may be less informative. We defined chemically useful poses as those with an overall RMSD ≤ 2 Å, a pharmacophore RMSD ≤ 1.5 Å, and a volume overlap ≥ 70%. For symmetric or pseudosymmetric ligands, only a pharmacophore RMSD ≤ 1.5 Å and a volume overlap ≥ 70% were required (**Figure 1B**).

Additionally, AF3 reports several confidence metrics that help interpret the reliability of the predicted binding poses. Two particularly useful scores for evaluating protein-ligand predictions are predicted aligned error (PAE) and the interface predicted template modeling score (ipTM). Predicted aligned error (PAE) is an AF3 confidence metric that quantifies ligand–protein prediction error as local, ordered token-pair-level positional uncertainty rather than a single global RMSD-like deviation. PAE estimates the positional uncertainty between pairs of tokens in the predicted structure, such as protein residues and ligand atoms, and is reported as an error distance in angstroms. Conceptually, minPAE is analogous to RMSD: lower PAE values indicate higher confidence in the relative placement of one token with respect to another. For protein–ligand complexes, the protein-to-ligand minimum PAE (minPAE) can provide a local confidence measure for whether at least part of the ligand is confidently positioned relative to the protein binding site. In contrast, ipTM converts ligand–protein token-pair confidence into an overall interface-level score rather than reporting a local positional error or global RMSD-like deviation. It estimates the confidence of predicted interactions between different chains or molecular components in the complex, with higher ipTM values indicating greater confidence in the predicted interface. Together, PAE and ipTM provide complementary information: PAE offers a local, distance-based estimate of positional uncertainty, whereas ipTM provides a broader interface-level confidence score. In this study, we used the AF3-reported protein-to-ligand minimum PAE and overall ipTM to assess whether this confidence metrics correlate with ligand pose accuracy, pharmacophore recovery, and ligand volume overlap.

The structures were selected from entries released after September 30, 2021, to ensure they were outside the AF3 training set. For each ligand–target complex, 10 structural predictions were generated with each model. Detailed sampling procedures and input specifications are provided in the Experimental Section. Together, these metrics distinguish global pose recovery, shape-level agreement, and preservation of chemically important recognition features across diverse drugging modalities. The PDB entries, their corresponding index numbers, and annotations are provided in **Table S1**.

### 1. Pose Recovery in New sEH Co-crystals with Flexible Lipid-Binding Pockets

Soluble epoxide hydrolase (sEH, EPHX2) is a lipid-metabolizing enzyme that hydrolyzes endogenous epoxy fatty acids, such as epoxyeicosatrienoic acids, to the corresponding vicinal diols (**Figure 2A**).^16–18^ In human sEH, catalysis is mediated by a highly conserved active-site network comprising D335, Y383, Y466, D496, and H524. D335 serves as the nucleophile that attacks the epoxide, Y383 and Y466 activate and orient the epoxide through hydrogen bonding, and the H524/D496 pair supports hydrolysis of the covalent enzyme intermediate (**Figure 2B**).^19^ Structurally, the sEH active site is embedded within a largely hydrophobic cavity that accommodates lipid-like substrates and inhibitors. This cavity is commonly divided into two flexible hydrophobic regions, the long-branch pocket (LB) and short-branch pocket (SB), which adapt to the long and short aliphatic chains of endogenous epoxy lipids and synthetic inhibitors.^20^ Inhibitor-bound structures demonstrate that urea-based pharmacophores mimic the transition state by forming a conserved hydrogen-bonding network with these catalytic residues, while bulky hydrophobic substituents extend into the long- and short-branch pockets (**Figure 2C**).^21^ This combination of a well-defined catalytic network, two large and flexible lipid-accommodating pockets, and a diverse set of chemical scaffolds makes sEH a useful challenge case for AF3 and Boltz-2, because accurate prediction requires both correct placement of the ligand polar pharmacophore near the catalytic residues and appropriate accommodation of variable hydrophobic substituents within adaptable binding pockets. This dual requirement underlies the sensitivity of sEH to ligand orientation and contributes to the difficulty of accurately predicting binding poses for diverse ligands.

**Figure 2.**
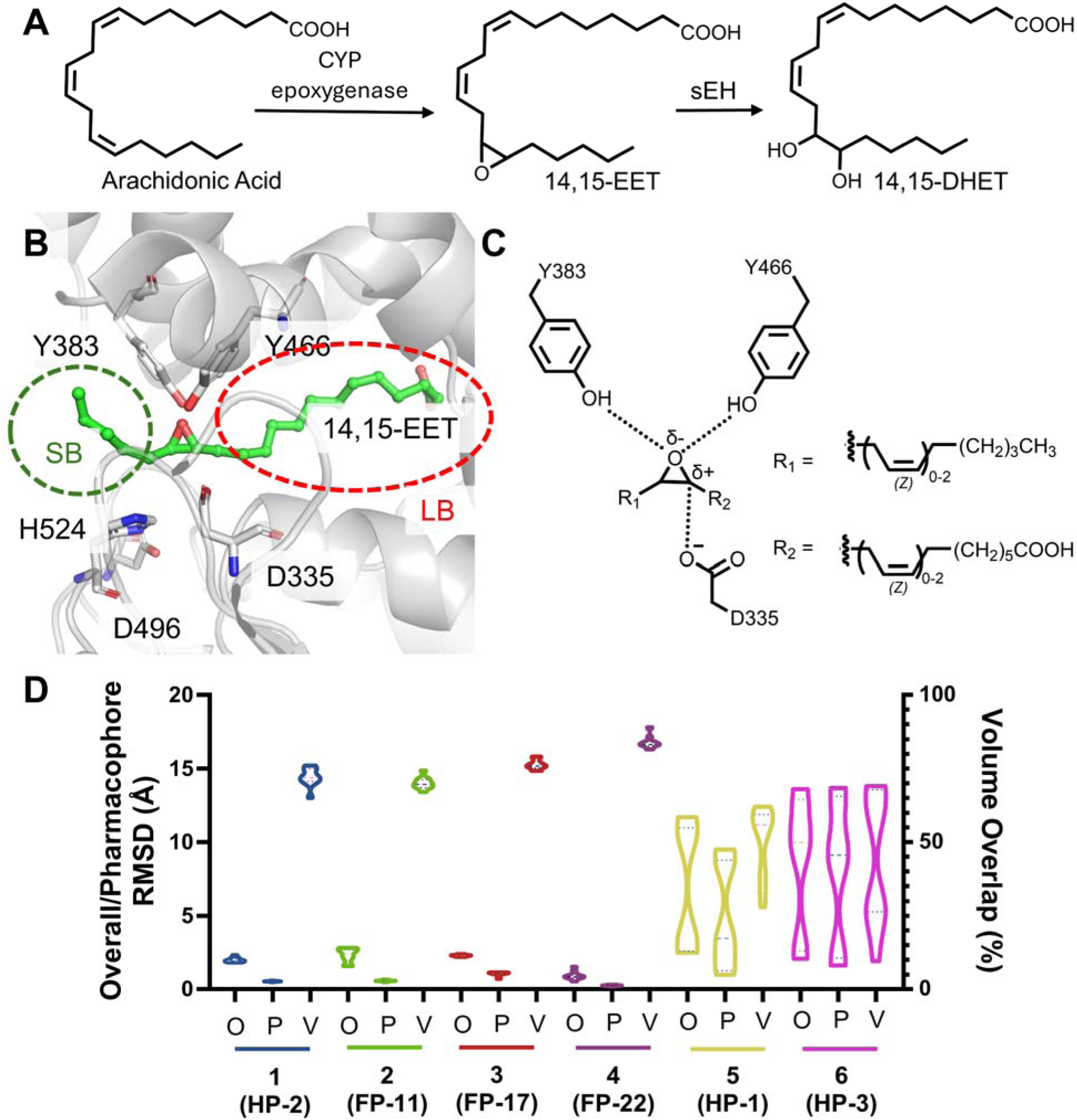
Substrate-based model for evaluating AF3 prediction of sEH ligand recognition. (**A**) Conversion of arachidonic acid to 14,15-EET by CYP epoxygenase, followed by hydrolysis of 14,15-EET to 14,15-DHET catalyzed by soluble epoxide hydrolase (sEH). (**B**) AF3-predicted placement of 14,15-EET within the sEH catalytic pocket, highlighting the catalytic residues Y383, Y466, and D335. The short-branch (SB) pocket and long-branch (LB) pocket are highlighted by green and red dashed circles, respectively. (**C**) Two-dimensional interaction model illustrating the substrate-recognition geometry used as a mechanistic reference for sEH inhibitor binding, including the polar catalytic interaction network and hydrophobic substituent orientation. (**D**) Quantitative evaluation of six sEH inhibitor predictions using ligand overall RMSD (O), pharmacophore RMSD (P), and ligand volume overlap (V).

AF3 and Boltz-2 showed generally good pose-recovery performance across the six newly solved sEH co-crystal structures (To be published),^22,23^ with AF3 accurately reproducing four of the six ligand poses (HP-2, FP-11, FP-17, and FP-22), whereas Boltz-2 showed comparable overall performance (**Figure 2D**, **Figure S1**). The molecular structures and associated testing data are provided in **Table S2**. In the successfully predicted cases, both models placed the central polar pharmacophore near the catalytic network and correctly oriented the hydrophobic substituents into the LB and SB pockets (**Figure 3A-F**). Notably, the failed case, HP-1, was not complete a failure of molecular recognition. Instead, the models largely preserved the key hydrogen-bonding interactions with the sEH catalytic residues, including D335, Y383, and Y466, but generated flipped binding modes in which the ligand arms were exchanged between the LB and SB pockets. These poses therefore retained the correct active-site pharmacophore while misassigning the topology of pocket occupancy.

**Figure 3.**
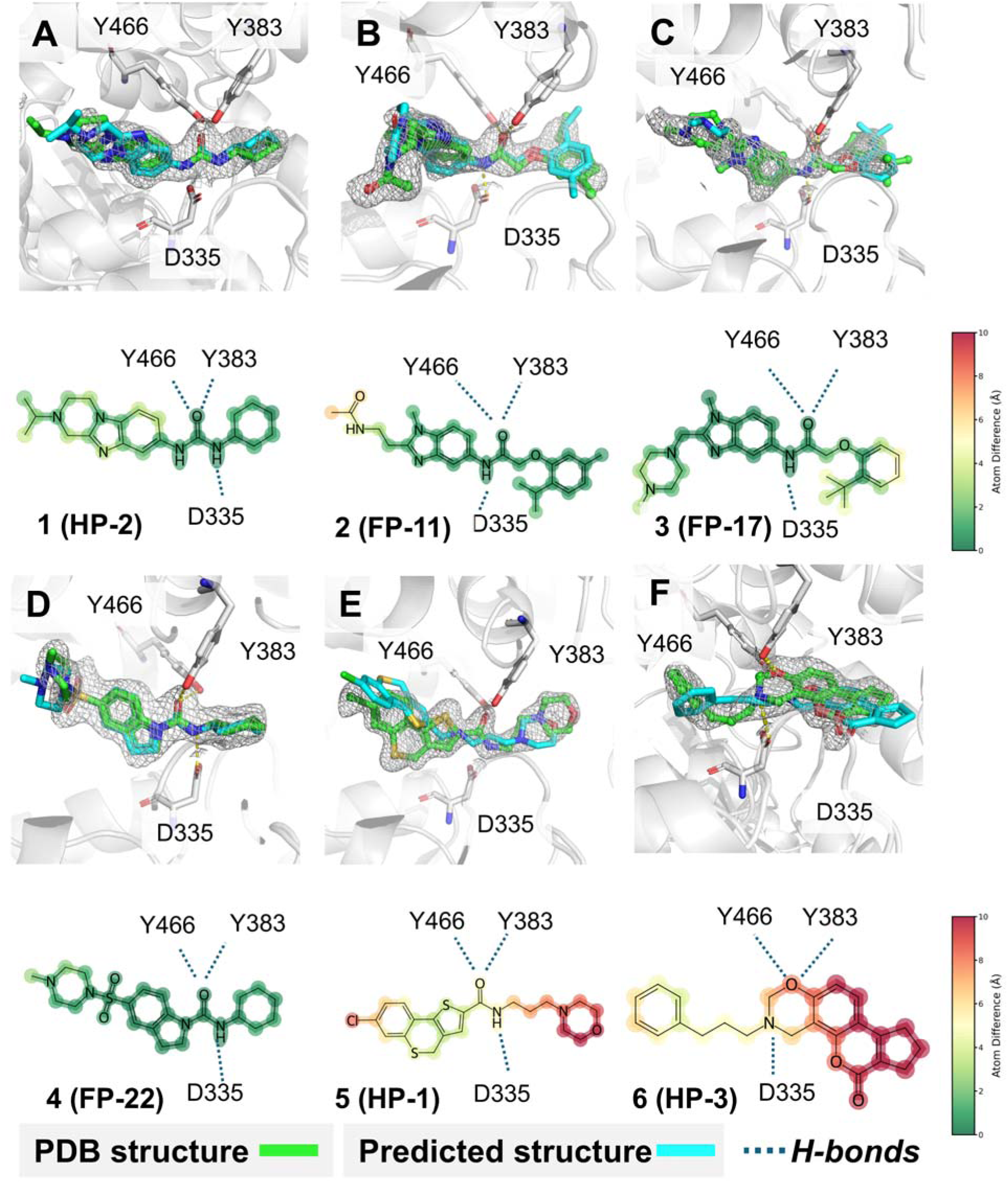
AlphaFold 3 pose prediction for six sEH inhibitor co-crystal structures. (**A**–**F**) Comparison of AF3-predicted and experimentally determined binding poses for six sEH inhibitors in the sEH catalytic pocket. In the structural overlays, experimental ligands are shown in green, AF3-predicted ligands are shown in cyan, and the catalytic residues Y383, Y466, and D335 are highlighted. The corresponding two-dimensional interaction maps show per-atom prediction deviations, colored from green for low deviation to red for high deviation, illustrating preservation or disruption of the catalytic pharmacophore and hydrophobic-arm placement across the sEH inhibitor series.

### 2. Performance in Canonical Orthosteric Ligand-Binding Complexes, with Emphasis on Kinases

We next evaluated AF3 and Boltz-2 using a diverse set of canonical orthosteric ligand-bound complexes, with particular emphasis on kinase-family ligands. The dataset includes JAK2 (**7**–**12**),^24^ TYK2 JH2 (**13**),^25^ BRAF (**14**),^26^ EGFR (**15-16**),^27^ ERK2 (**17**),^28^ PDE4D (**18**) ^29^ Hck (**19**),^30^ p38α (**20**-**21**),^31^ PPAR (**22**-**23**),^32^ and PLpro (**24**-**25**) complexes,^33^ spanning viral protease inhibitors, canonical kinase ATP-site inhibitors, conformationally selective kinase inhibitors, pseudokinase-domain ligands, and bivalent EGFR inhibitors that engage both orthosteric and allosteric regions. This collection provides a useful test of pose prediction in enzyme-centered binding sites because productive ligand recognition is often guided by conserved pharmacophores, such as kinase hinge interactions or protease substrate-groove engagement, while still requiring accurate recovery of local protein conformation. For kinase-family targets, pose accuracy may depend not only on ATP-site occupancy, but also on DFG-loop organization, αC-helix positioning, activation-state differences, and, in the case of TYK2 JH2 or bivalent EGFR inhibitors, recognition of regulatory or allosteric binding features. Thus, this target set allows us to examine how AF3 and Boltz-2 perform across enzyme-centered recognition problems ranging from conventional active-site binding to more complex conformational and mixed orthosteric/allosteric engagement.

#### 2.1 High Pose-Recovery Accuracy in Kinase-Centered Binding Sites

AF3 performed particularly well across this dataset, accurately recovering ligand poses in all 15 kinase complexes and both PPAR nuclear receptor complexes (**Figure 4A-C**, complexes **7**–**23**). The two PLpro complexes (**24** and **25**) were the only exceptions and are discussed below. Across the successful cases, the median RMSD across 10 generated poses for each complex was generally within 2 Å and often below 1.5 Å, indicating substantial progress in generating accurate poses for canonical orthosteric enzyme binding sites. In many cases, pharmacophore RMSD remained below ∼1 Å even when overall RMSD exceeded 2 Å, indicating that the core functional interactions of the ligand were accurately preserved while deviations primarily arose from flexible peripheral substituents or alternative conformations of solvent-exposed regions. Boltz-2 demonstrated a similar trend, although with slightly greater variability in several complexes. The corresponding Boltz-2 pose-distribution analysis is shown in **Figure S2**, while the molecular structures and associated testing data are provided in **Table S3**.

**Figure 4.**
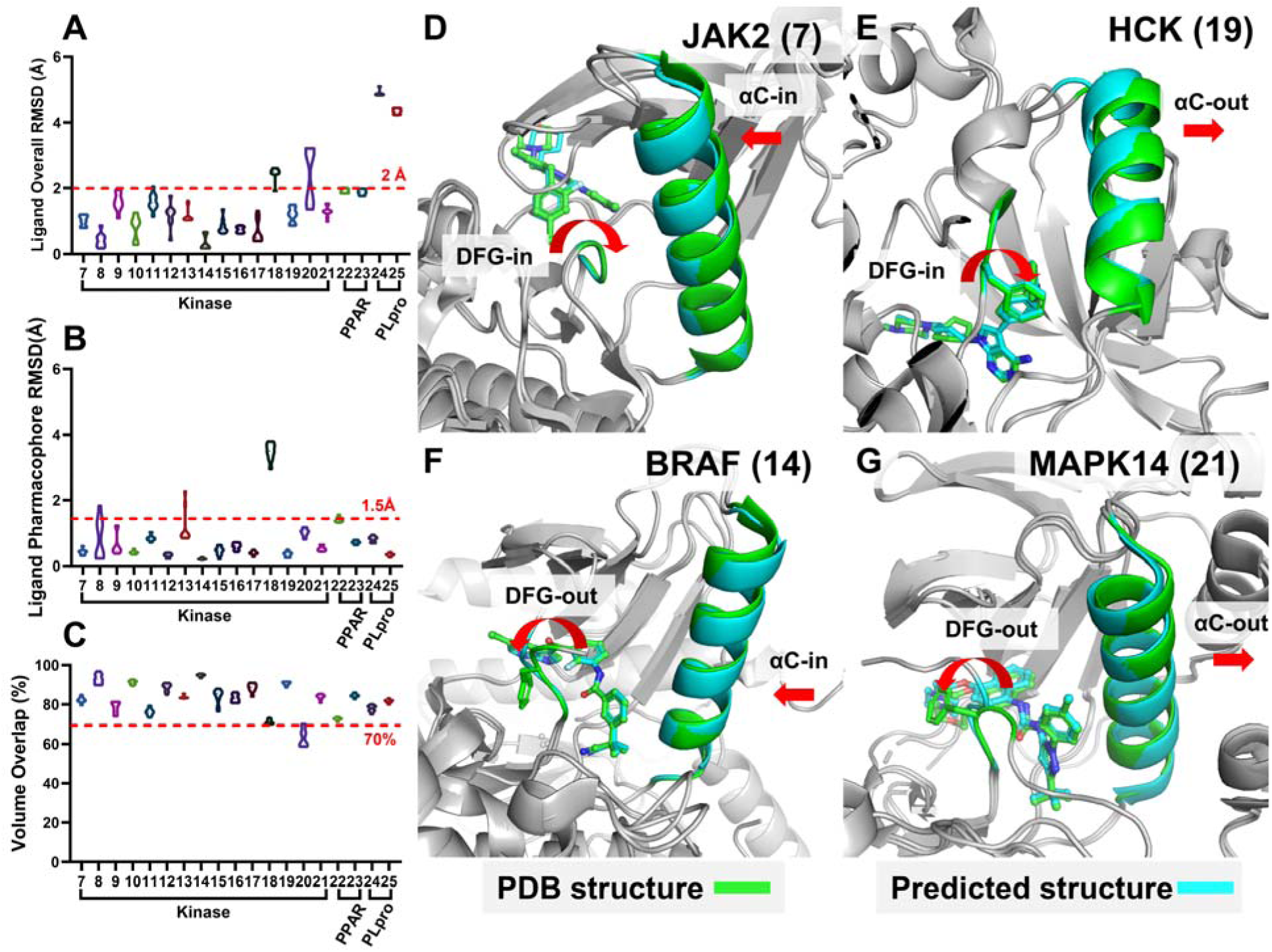
Pose prediction performance for canonical orthosteric inhibitor complexes, with emphasis on kinases. (**A**–**C**) Quantitative evaluation of ligand pose recovery across entries **7**–**25** using ligand overall RMSD, ligand pharmacophore RMSD, and ligand volume overlap. Dashed red lines indicate the predefined thresholds for chemically useful pose recovery: 2 Å for overall RMSD, 1.5 Å for pharmacophore RMSD, and 70% for volume overlap. (**D**–**G**) Representative kinase complexes illustrating recovery of ligand-associated regulatory conformations, including JAK2 (**7**: 8BM2), HCK (**19**: 9BYJ), BRAF (**14**: 8C7Y), and MAPK14/p38α (**21**: 9CJ4). Experimental structures are shown in green, predicted structures in cyan. These examples show that AF3 can recover both ligand pose and key kinase conformational features, including DFG-in/out and αC-in/out states. For entries **7**–**25**, the corresponding PDB structures are 8BM2, 8BPW, 8BX6, 8BX9, 8BXC, 8BXH, 8S9A, 8C7Y, 8FV3, 8FV4, 8U8J, 8ZQW, 9BYJ, 9CJ1, 9CJ4, 8HUK, 8HUQ, 8UOB, and 9CSY.

These observations suggest that both AF3 and Boltz-2 preferentially preserve conserved interaction motifs that dominate molecular recognition in canonical enzyme binding sites. In kinase-family targets and related enzyme-centered pockets, ligand binding is typically anchored by highly conserved pharmacophore interactions, such as hinge-region hydrogen bonding networks or catalytic residue contacts, whereas distal hydrophobic substituents often occupy larger and more conformationally permissive regions of the binding pocket. As a result, substantial differences in overall ligand orientation may occur without disrupting the key interaction geometry required for functional recognition.

The discrepancy between overall RMSD and pharmacophore RMSD further indicates that conventional whole-ligand RMSD may overestimate prediction errors for ligands containing flexible tails, symmetric substituents, or solvent-exposed peripheral groups. In contrast, pharmacophore-restricted RMSD more directly captures the accuracy of functionally relevant binding interactions and therefore provides a more mechanistically meaningful metric for evaluating ligand pose prediction in enzyme-centered binding sites.

Consistent with this interpretation, volume overlap measurements remained relatively high across most complexes even when overall RMSD increased, supporting the conclusion that many predicted poses retained broadly correct occupancy within the binding pocket despite local conformational differences.

These results suggest that kinase-family active sites and related enzyme-centered pockets provide a favorable setting for AF3 and Boltz-2 pose prediction, likely because ligand recognition is strongly constrained by conserved pharmacophores, defined pocket geometry, and recurring interaction motifs such as kinase hinge binding.

#### 2.2 Recovery of Ligand-Stabilized DFG/**α**C Kinase States

Kinase catalytic domains can adopt multiple conformational states that are broadly associated with active and inactive functional states. Transitions among these states involve two conserved structural elements: the Asp-Phe-Gly (DFG) motif and the αC helix. In the active state, the DFG motif typically adopts a DFG-in conformation, in which the aspartate side chain is oriented toward the ATP-binding site to coordinate metal ions required for nucleotide binding and catalysis. In the αC-in conformation, a conserved glutamate in the αC helix forms a regulatory salt bridge with a lysine in the β3 strand. Among the major kinase conformational states, the αC-in/DFG-in state corresponds to the canonical catalytically active conformation.^34^

To further assess whether AF3 and Boltz-2 can recover ligand-associated kinase conformational states, we selected four kinase complexes in which pose prediction is coupled to regulatory structural features: JAK2 bound to gandotinib (**7**; **Figure 4D**), BRAF V600E bound to TXV (**14**; **Figure 4E**), Hck bound to A-419259 (**19**; **Figure 4F**), and dual-phosphorylated p38α bound to BIRB796 (**21**; **Figure 4G**). In 7, Gandotinib was reported to bind the ATP site of JAK2 and stabilize a canonical DFG-in/αC helix-in active state with an intact regulatory salt bridge, providing a test of whether the models recover both standard Type I ligand placement and the associated fully active kinase conformation. In **14**, a paradox-breaking hybrid inhibitor binds BRAF(V600E) and stabilizes a rare DFG-out/αC-helix-in conformation in which the ligand occupies the deep allosteric pocket while maintaining the inward orientation of the αC-helix. In **19**, A-419259 was reported to bind the ATP site of Hck and stabilize a DFG-in/αC-helix-out state with an extended activation loop, providing a test of whether the models recover both ATP-site ligand placement and the associated Src-family kinase conformation. In **21**, nilotinib and BIRB796 bind dual-phosphorylated p38α and stabilize inhibitor-associated activation-loop conformations in which the phospho-threonine is exposed.

Surprisingly, AF3 and Boltz-2 were particularly effective in these cases, accurately recovering not only the ligand poses but also the associated kinase conformational features, including DFG-loop organization, αC-helix positioning (pink parts), and activation-loop arrangement. This ability to generate ligand-bound protein conformations highlights a key strength of diffusion-based structure prediction relative to traditional rigid-receptor docking, where the protein conformation is usually fixed or only locally relaxed. Together, these examples suggest that AF3 and Boltz-2 may be especially useful for kinase systems in which medicinal chemistry interpretation depends on both ligand pose and ligand-associated protein-state selection.

#### 2.3 Recovery of Ligand-Induced Coregulator Recruitment to Transcription Factors

We next evaluated AF3 and Boltz-2 on transcription factor systems in which small-molecule binding induces or stabilizes recruitment of peptide coregulators. These complexes provide a distinct challenge from canonical enzyme active sites because successful prediction requires simultaneous recovery of three coupled recognition events: small-molecule binding, peptide coregulator positioning, and the protein conformational state that accommodates both interactions. Unlike rigid orthosteric enzyme pockets, transcription factor regulatory surfaces are frequently shallow and partially solvent exposed, and peptide recognition often depends on ligand-induced stabilization of transient protein conformations.

To examine this class of complexes, we analyzed the transcription factor PPARα, **22** and **23**, each containing both a small-molecule ligand and a bound peptide coregulator. Remarkably for **22**, both AF3 and Boltz-2 demonstrated strong performance across these systems, accurately recovering not only the small-molecule binding pose but also the associated peptide binding geometry (**Figure 5A**). Across both complexes, ligand pharmacophore RMSD values remained low and volume overlap values remained high, indicating substantial preservation of the experimentally observed binding mode. Peptide RMSD values were similarly favorable, suggesting that both models successfully captured the coupled recognition between ligand binding and coregulator recruitment (**Figure 5B**).

**Figure 5.**
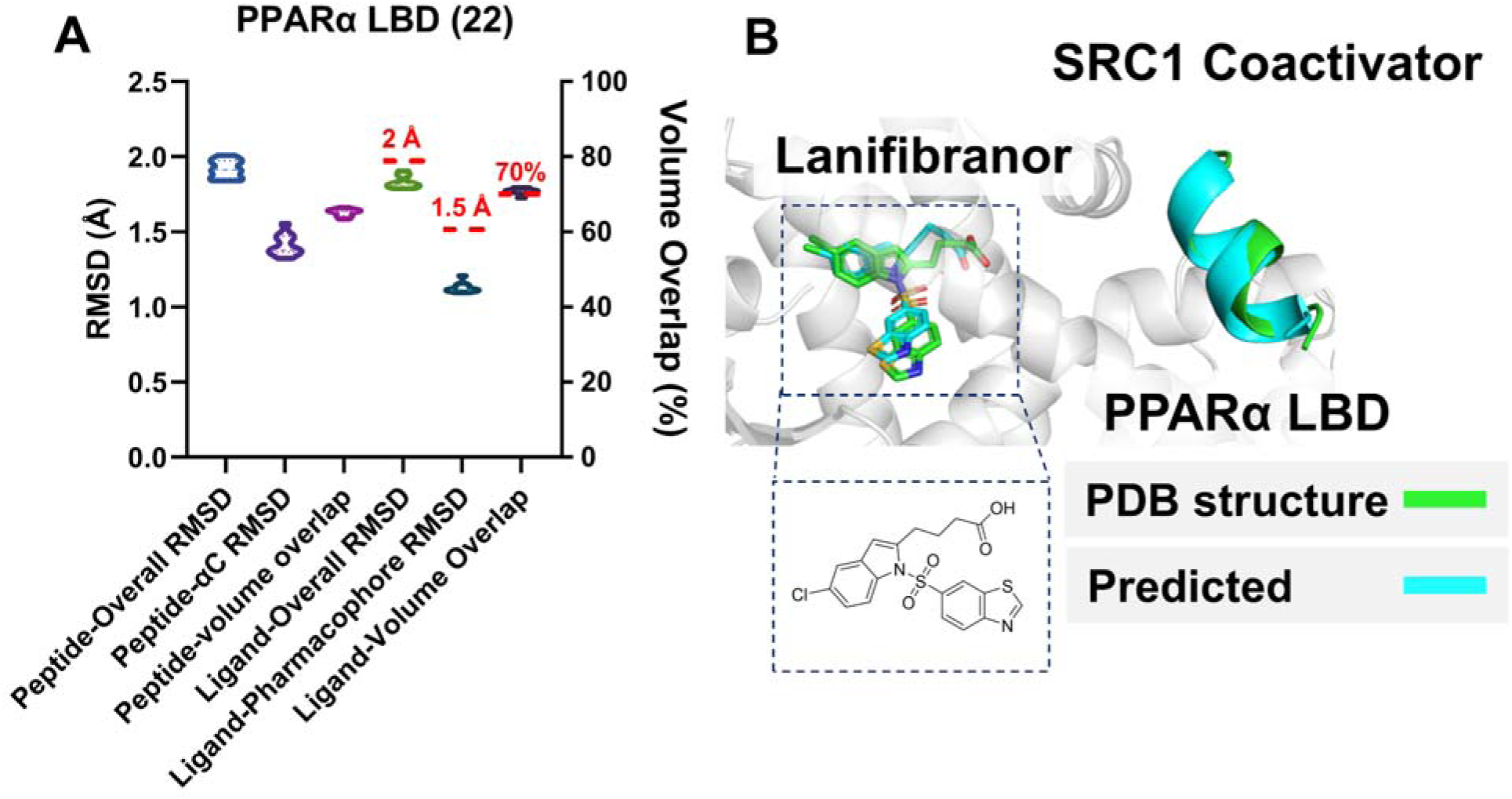
AF3 prediction of ligand-induced coregulator recruitment in the PPARα transcription factor complex. (**A**) Pose-evaluation metrics for the PPARα–lanifibranor–SRC1 coactivator complex (**22**: 8HUK), including peptide overall RMSD, peptide Cα RMSD, peptide volume overlap, ligand overall RMSD, ligand pharmacophore RMSD, and ligand volume overlap. (**B**) Representative AF3-predicted ternary assembly overlaid with the experimental structure, showing lanifibranor bound within the PPARα ligand-binding domain (LBD) and recruitment of the SRC1 coactivator peptide. The ligand structure is shown in the inset.

AF3 consistently showed slightly improved performance relative to Boltz-2 across both systems, exhibiting lower pose variability and more accurate recovery of peptide and ligand geometry. AF3 generated highly consistent ligand pharmacophore placement together with accurate positioning of the peptide coregulator within the induced binding interface. The relatively narrow violin plot distributions further suggest that AF3 sampled a more stable conformational solution across independently generated models.

The successful prediction of **22** and **23** contrasts sharply with the poorer performance observed by many highly dynamic allosteric systems (**Sections 3-6**). One likely explanation is that ligand-induced coregulator interfaces in these transcription factor complexes form a comparatively well-defined composite binding surface once the ligand is bound. In this context, the small molecule acts as a stabilizing scaffold that constrains the peptide-binding interface, thereby reducing conformational ambiguity and facilitating simultaneous recovery of both ligand and peptide recognition. These findings suggest that AF3 and Boltz-2 are capable of accurately modeling coupled small-molecule and peptide recognition when ligand binding induces a sufficiently stable and geometrically constrained regulatory interface.

#### 2.4 Pseudo-Symmetry and the Need for Pharmacophore- and Volume-Aware Evaluation

We next examined two SARS-CoV-2 PLpro predictions as instructive cases in which conventional pose-evaluation metrics diverged (**Figure 6**). PLpro is a multifunctional viral cysteine protease and deubiquitinase whose substrate-binding cleft mimics ubiquitin recognition, with the flexible BL2 loop gating access to the active-site region. In both PLpro examples, AF3 and Boltz-2 showed a large discrepancy among ligand RMSD, pharmacophore RMSD, and volume overlap. Although ligand RMSD alone suggested poor pose recovery, the pharmacophore RMSD and volume-overlap metrics indicated substantial preservation of the binding mode. This discrepancy indicates that the models captured the overall pharmacophore placement and BL2-loop-associated pocket organization but did not always reproduce the atom-mapped ligand orientation within the ubiquitin Val70-recognition region and the adjacent BL2-groove binding site.

**Figure 6.**
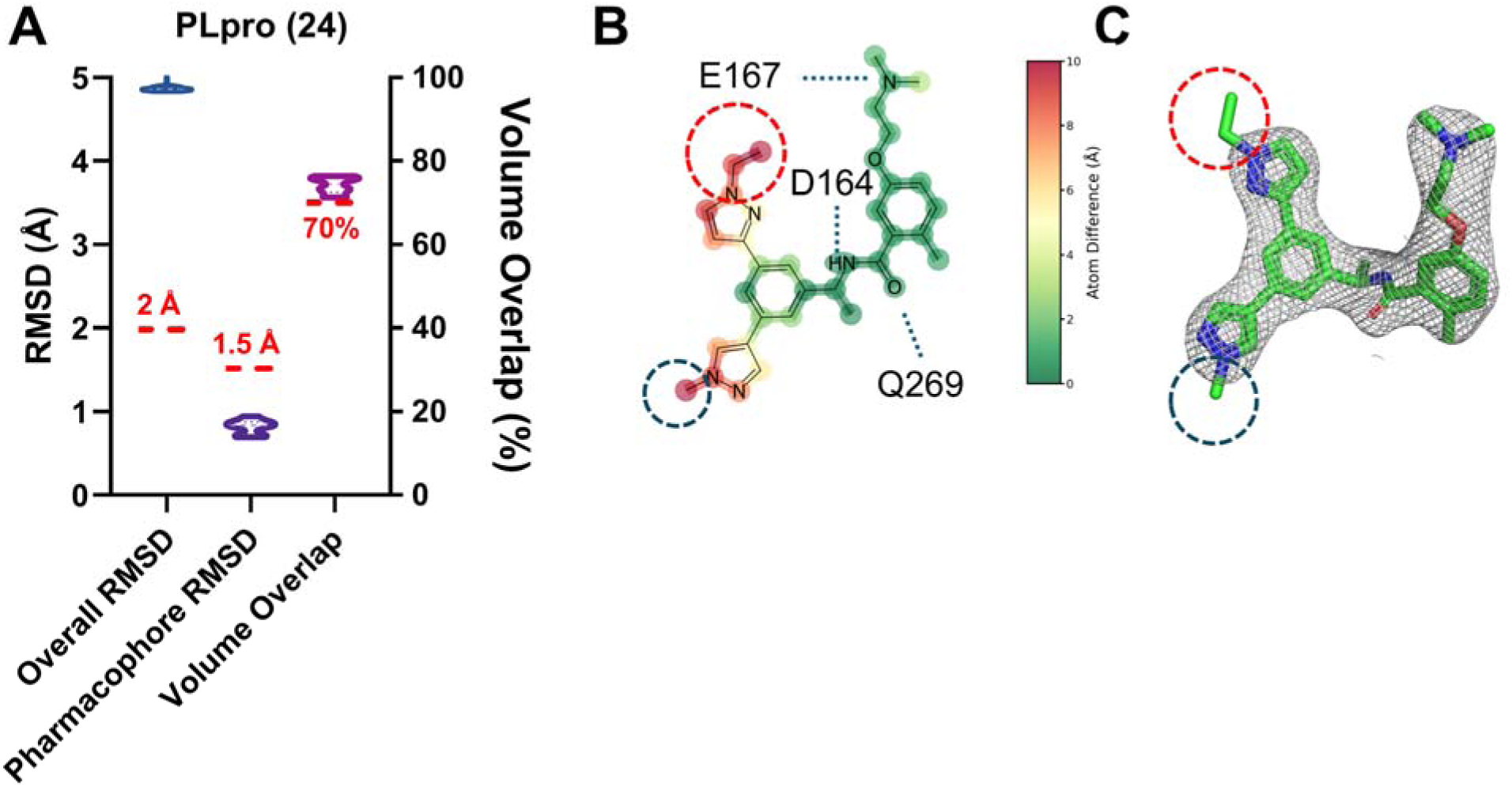
AF3 prediction of a pseudo-symmetric SARS-CoV-2 PLpro inhibitor complex. (**A**) Pose-evaluation metrics for the PLpro–Jun12682 complex (**24**: 8UOB), including overall RMSD, pharmacophore RMSD, and volume overlap. (**B**) Two-dimensional interaction map of the AF3-predicted pose, with dashed circles marking regions of orientation ambiguity associated with the pseudo-symmetric scaffold. (**C**) Experimental 2Fo-Fc electron-density map contoured at 1.0σ for Jun12682, showing limited discrimination between the two alternative ligand orientations within the circled regions, consistent with the observed pseudo-symmetry.

The **24** complex illustrates why this distinction is important (**Figure 6A**). The ligand contains a pseudo-symmetric scaffold with similarly substituted pyrazole groups, allowing alternative orientations that preserve the same general interaction pattern and occupy a similar pocket volume while producing a high atom-mapped ligand RMSD. Inspection of the corresponding electron density further supports this interpretation: the density does not clearly define a unique ligand orientation, suggesting that the AF3- or Boltz-2-predicted pose may not be strictly incorrect but may instead represent an alternative geometrically plausible solution compatible with the experimental map. In this case, ligand RMSD alone over penalizes a symmetry-related pose, whereas pharmacophore RMSD and volume overlap better capture the preserved chemical recognition (**Figure 6B, 6C**). Together, these examples demonstrate how symmetry-aware, pharmacophore- and volume-based evaluation provides a more chemically interpretable assessment of AI-generated ligand poses than ligand RMSD alone.

### 3. Performance in Allosteric Modulators

Following evaluation of canonical orthosteric enzyme inhibitors, we next examined AF3 and Boltz-2 performance on allosteric modulators (**Figure 7A-C**, **Figure S3**). This class provides a stringent test for pose prediction because allosteric binding often depends on induced-fit, cryptic-pocket formation, or stabilization of a transient protein conformation rather than recognition of a preorganized active site. Unlike kinase ATP sites or protease substrate grooves, allosteric sites are frequently shallow, dynamic, and scaffold dependent, requiring accurate prediction of both ligand placement and ligand-induced pocket formation.

**Figure 7.**
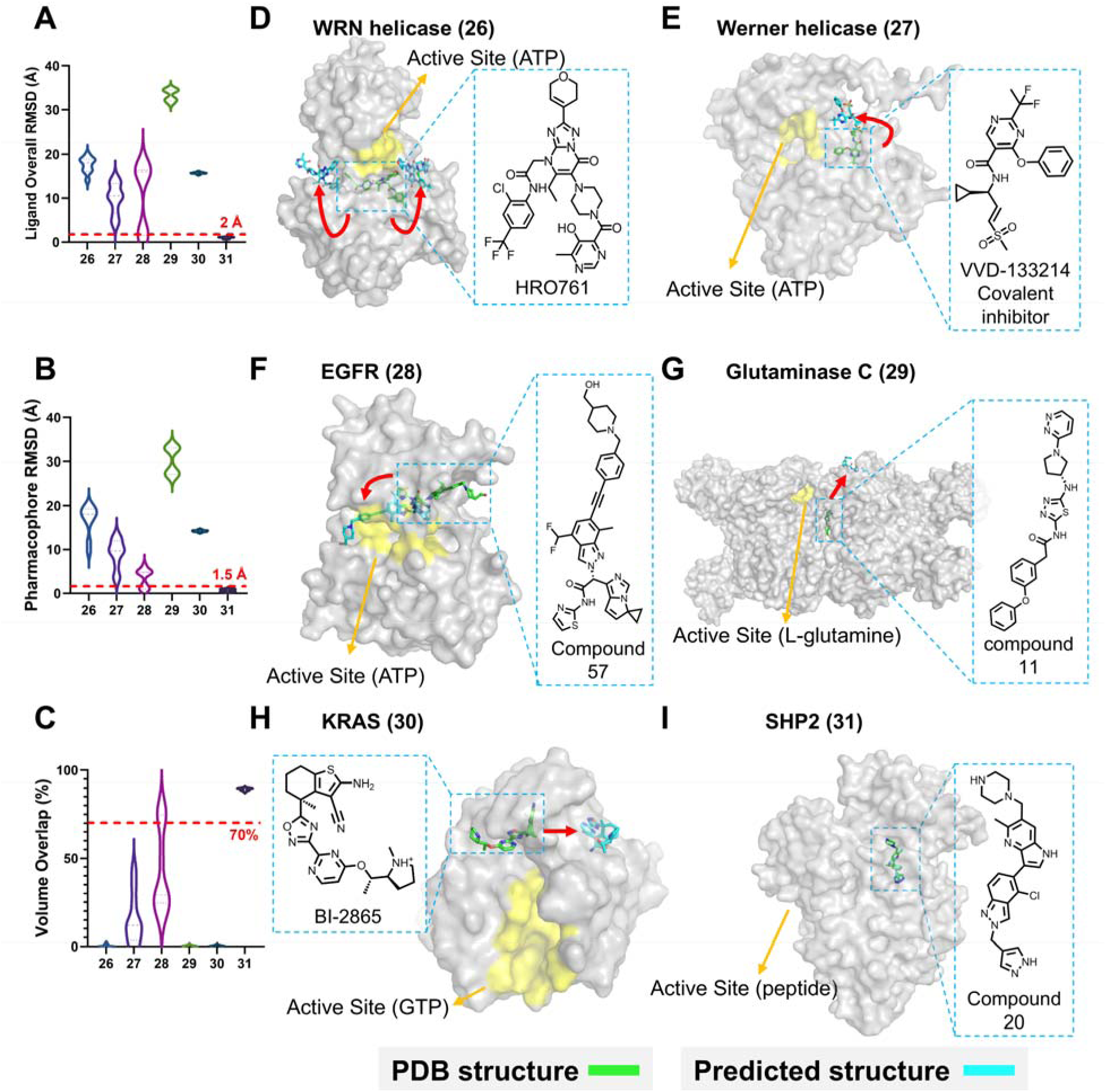
AF3 analysis of allosteric inhibitor recognition across diverse regulatory binding sites. (**A–C**) Dataset-wide evaluation using ligand overall RMSD, ligand pharmacophore RMSD, and ligand volume overlap. (**D–I**) Representative examples spanning helicase, kinase, metabolic enzyme, phosphatase, and KRAS allosteric systems. Experimental active sites are indicated to emphasize separation between orthosteric and allosteric ligand-binding regions. Examples include WRN helicase inhibitors HRO761 (**26**: 8PFO) and covalent inhibitor VVD-133214 (**27**: 7GQU), allosteric EGFR inhibitor complex (**28**: 8A2D), Glutaminase C inhibitor complex (**29**: 8JUE), KRAS Switch-II pocket ligand BI-2865 (**30**: 8AZV), and SHP2 allosteric inhibitor complex (**31**: 8S0O). These systems highlight the challenges posed by induced-pocket formation, conformational plasticity, and ligand-dependent regulatory-site recognition in allosteric pose prediction.

We curated a diverse allosteric modulator set, including WRN helicase inhibitors HRO761 (**26**; **Figure 7D**) and VVD-133214 (**27**; **Figure 7E**),^35,36^ the latter targeting an allosteric cysteine site; the first-in-class allosteric EGFR inhibitor complex (**28**; **Figure 7F**);^37^ the glutaminase C allosteric inhibitor complex (**29**; **Figure 7G**);^38^ the KRAS Switch II pocket binder BI-2865 (**30**; **Figure 7H**);^39^ and a fragment-based inhibitor of SHP2 (**31**; **Figure 7I**).^40^ These examples span reversible and covalent allosteric ligands, cryptic or induced pockets, regulatory sites, and shallow protein-surface binding sites. In contrast to the strong pose recovery observed by many canonical enzyme active-site inhibitors, both AF3 and Boltz-2 generally failed to reproduce the correct allosteric binding modes. The molecular structures and associated testing data are provided in **Table S4**.

An important exception within the allosteric dataset was the SHP2 fragment inhibitor complex (**31**), for which both AF3 and Boltz-2 achieved highly accurate pose recovery across ligand RMSD, pharmacophore RMSD, and volume overlap metrics. Unlike several other allosteric systems that required substantial induced-fit rearrangements or stabilization of transient conformational states, the **31** binding site appears comparatively preorganized and structurally constrained. The ligand occupies a compact pocket with limited conformational ambiguity, reducing the requirement for large-scale protein rearrangement during molecular recognition. In addition, the fragment-like scaffold contains fewer flexible substituents and a more localized interaction network than larger allosteric inhibitors, thereby reducing the number of alternatives geometrically plausible poses. These features likely decrease the conformational search complexity encountered by AF3 and Boltz-2 and enable more accurate recovery of the experimentally observed binding mode.

The main limitation was failure to recover the induced pocket geometry and local protein rearrangements required for productive ligand recognition. In several cases, the models generated plausible ligand placements near the correct region, but did not reproduce the experimentally observed molecular recognition pattern.

These findings indicate that allosteric modulators remain one of the most challenging categories for AF3 and Boltz-2. For medicinal chemistry applications, predicted allosteric poses should therefore be treated primarily as hypotheses and interpreted alongside experimental structures, SAR, mutagenesis, biophysical mapping, or physics-based simulations before being used to guide analog design.

### 4. Performance in Protein–Protein Interaction Inhibitors

We next evaluated AF3 and Boltz-2 on small-molecule inhibitors that disrupt protein–protein interactions (PPIs), a target class traditionally considered challenging in drug discovery.^14^ Unlike canonical enzymes and receptors, which often contain deep binding pockets for substrates, cofactors, or ligands, PPI interfaces are typically large, relatively flat, dynamic, and solvent-exposed.^14^ In some cases, small-molecule PPI inhibitors engage induced or cryptic pockets that form within these interfaces. These features make PPI inhibitors a stringent test for pose prediction because accurate modeling requires not only correct ligand placement, but also recognition of surface topology, local pocket formation, and interaction hot spots in regions that were not evolved to bind small molecules.^14,41^ The molecular structures and associated testing data are provided in **Table S5**.

To evaluate this modality, we selected a diverse set of PPI inhibitor complexes (**Figure 8A**-**C**, **Figure S4**), including Aurora A–TPX2 interaction inhibitors (**32**; **Figure 8D**),^42^ and the RhoGDI2–Rac1 PPI inhibitor complex (**33**; **Figure 8E**),^43^ SOS1/SOS2–KRAS interaction inhibitors (**34**; **Figure 8F**) and KEAP1–NRF2 inhibitors (**35**; **Figure 8G**).^44,45^ These systems represent distinct PPI-recognition mechanisms. Aurora A–TPX2 inhibitors interfere with a regulatory kinase–activator interaction, whereas RhoGDI2–Rac1 represents a more challenging interface involving a small GTPase and its regulatory binding partner. SOS1/SOS2–KRAS inhibitors bind at or near the KRAS–GEF interaction surface and disrupt nucleotide-exchange signaling. KEAP1–NRF2 inhibitors target a well-defined hotspot within the KEAP1 Kelch domain that recognizes the NRF2 degron motif. Together, these cases span hotspot-driven peptide-mimetic recognition, shallow-surface binding, and regulatory PPI disruption.

**Figure 8.**
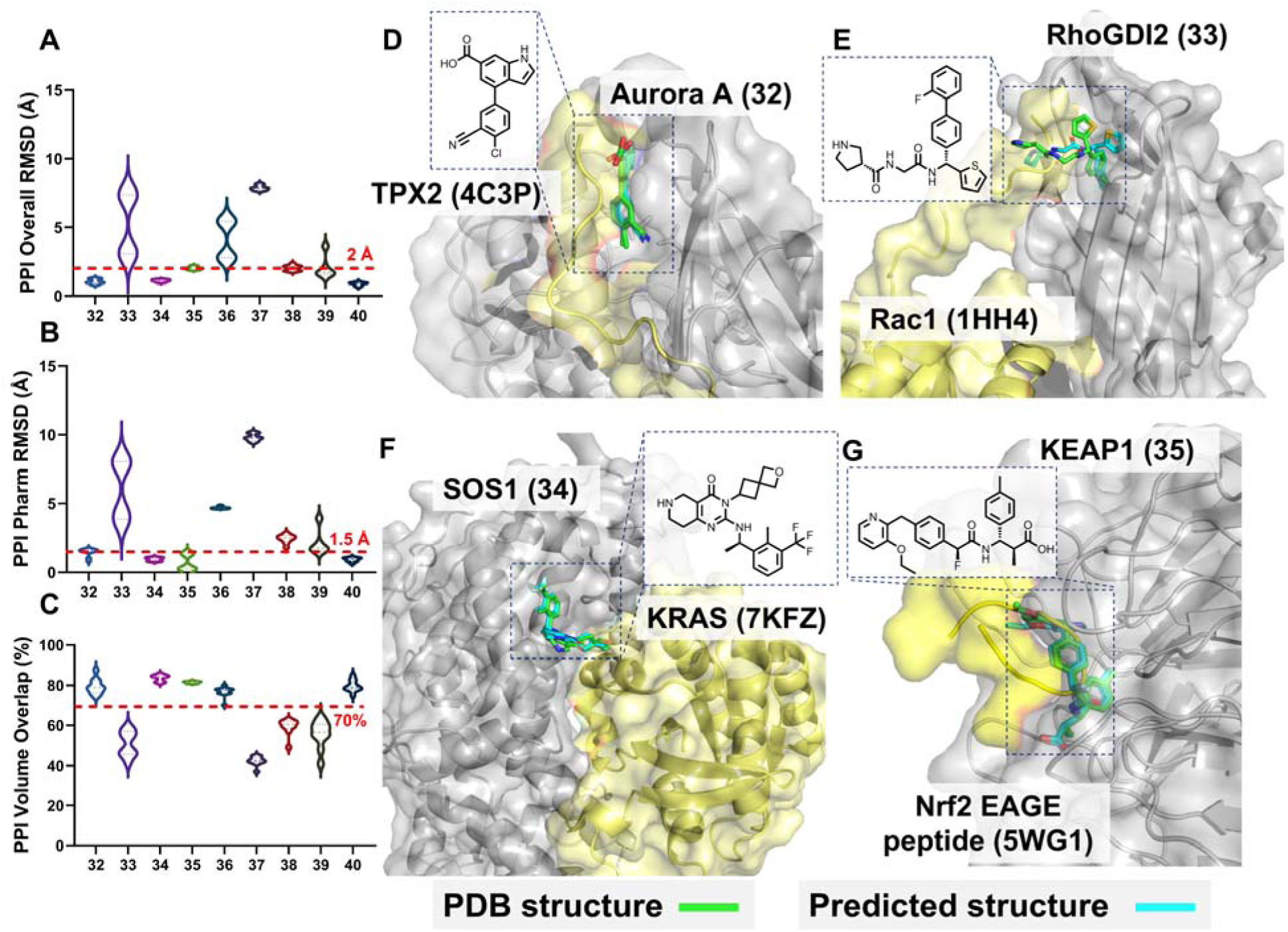
AF3 prediction performance for protein–protein interaction (PPI) inhibito complexes. (**A**) Ligand overall RMSD. (**B**) Ligand pharmacophore RMSD. (**C**) Ligand volume overlaps across the PPI inhibitor dataset. (**D–G**) Representative AF3 predictions illustrating ligand recognition within structurally defined protein–protein interaction interfaces. Structures are shown for Aurora A–TPX2 inhibitor complex (**32**: 8C14), and RhoGDI2–Rac1 inhibitor complex (**33**: 9X53), SOS1–KRAS inhibitor complex (**34**: 9MJM) and KEAP1–Nrf2 inhibitor complex (**35**: 8XGV). The corresponding endogenous protein–protein interfaces are indicated using KRAS (7KFZ), Nrf2 EAGE peptide (5WG1), TPX2 peptide (4C3P), and Rac1 (1HH4), respectively. AF3 generally performed well for hotspot-defined PPI pockets. For entries **36**-**40**, the corresponding structures are 8T5G, 8T5M, 8T5R, 8UC9 and 8UH0.

In contrast to the poor performance observed for many allosteric modulators, AF3 performed surprisingly well across this PPI-focused dataset, accurately recovering 6 of 9 tested complexes, with the notable exception of the RhoGDI2–Rac1 case. Although the predicted poses exhibited greater variability than canonical enzyme-targeted complexes, a substantial subset of generated models recovered ligand conformations with low RMSD and preserved pharmacophore geometry, indicating that AF3 could identify the correct binding mode despite incomplete convergence across all predictions. This result was unexpected because PPI inhibitors, like allosteric modulators, often lack an endogenous small-molecule ligand template comparable to ATP in kinases or substrates in enzymes or receptors. However, the favorable performance suggests that some PPI inhibitor binding sites provide clearer structural cues for pose prediction than typical cryptic allosteric pockets, particularly when the ligand engages a preorganized hot spot or peptide-recognition groove. These findings suggest that AF3 may have practical value for modeling PPI inhibitors, especially in systems where the small molecule occupies a structurally defined interface pocket rather than relying on extensive cryptic-pocket formation. More broadly, this result highlights a potentially important application of AF3 in addressing historically difficult PPI targets, while also emphasizing the need to distinguish hotspot-defined PPI pockets from highly dynamic or weakly preorganized protein surfaces.

### 5. Performance in Heterobifunctional PROTAC Complexes

Proteolysis-targeting chimeras (PROTACs) represent a distinct induced-proximity modality in which a heterobifunctional molecule simultaneously engages a protein of interest and an E3 ubiquitin ligase to promote target ubiquitination and proteasomal degradation.^46^ This mechanism can produce functional outcomes that are not achievable through occupancy-driven inhibition alone, particularly when target removal, rather than active-site blockade, is required for biological activity.^47^ From a structural-prediction perspective, PROTACs are challenging because productive degradation depends not only on binary binding to the target and E3 ligase, but also on the geometry, cooperativity, linker conformation, and protein–protein interface of the induced ternary complex.^48^

We focused on two widely used E3 ligase systems, cereblon (CRBN) and von Hippel–Lindau protein (VHL). CRBN functions as the substrate receptor of the CRL4^CRBN E3 ubiquitin ligase complex, in association with DDB1, CUL4, and RBX1, and has been extensively exploited by thalidomide-derived immunomodulatory drugs and CRBN-recruiting degraders. VHL is the substrate-recognition component of the CRL2^VHL ligase complex, in association with elongin B, elongin C, CUL2, and RBX1, and is commonly recruited by hydroxyproline-based ligands in VHL-directed PROTACs (**Figure 9A-C**). To evaluate this modality, we selected two recently reported cryo-EM ternary-complex structures: CDK2/Cyclin E1 in complex with CRBN/DDB1 and a CDK-targeting degrader (**41**; **Figure 9D**, **E**),^49^ and KRAS G12D C118S–GDP in complex with the pVHL:elongin C:elongin B complex and a KRAS-directed VHL-recruiting degrader (**42**; **Figure 9F**, **G**).^50^ These two ligase systems provide complementary test cases because they differ in ligand chemistry, substrate-recognition architecture, ternary-complex topology, and degrader-induced protein–protein interfaces. The molecular structures and associated testing data are provided in **Table S6**.

**Figure 9.**
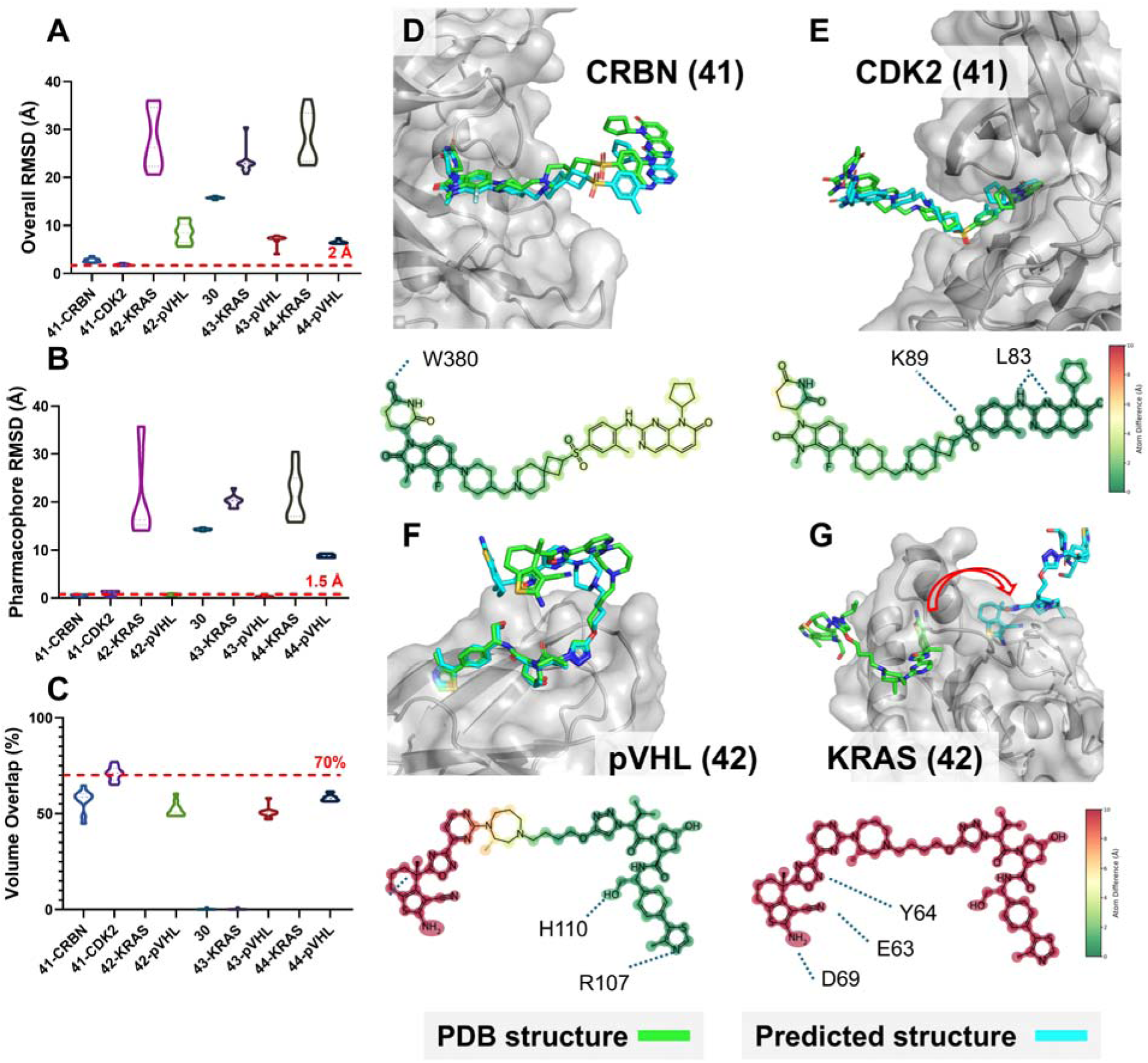
AF3 prediction analysis of PROTAC heterobifunctional degraders and ternary complexes. (**A**–**C**) Quantitative evaluation of ligand overall RMSD, pharmacophore RMSD, and volume overlap for the degrader-associated dataset. Dashed red lines indicate the predefined thresholds for chemically useful pose recovery. (**D&E**) AF3 prediction of the CDK2/Cyclin E1–CRBN/DDB1 degrader complex (**41**: 9D0X), shown after alignment to CRBN (D) or CDK2 (E). (**F&G**) AF3 prediction of the KRAS–pVHL degrader complex (**42**: 8QVU), shown after alignment to pVHL (F) or KRAS (G). Predicted ligand poses are overlaid with the corresponding experimental structures in the E3 ligase and target-protein binding pockets. The associated two-dimensional interaction maps show per-atom prediction deviations, colored from green for low deviation to red for high deviation. For entries **30**, **43**, and **44**, the corresponding structures are 8AZV, 9RK8, and 9L6F, respectively.

In the CDK2–CRBN system, AF3 consistently recovered the overall degrader trajectory and generated relatively accurate placement of both the CRBN-binding and CDK2-binding moieties, as reflected by the pharmacophore RMSD and volume-overlap metrics. The CRBN-binding and CDK2-binding moieties remained comparatively well aligned with the experimental structure, whereas the central rigid linker adopted several alternative trajectories across models. This finding is consistent with the earlier individual-ligand analysis, as both CDK2 and CRBN contain canonical orthosteric binding sites with abundant complex structures represented in the PDB dataset. Boltz-2 generated a similar global ternary architecture but exhibited greater conformational diversity, particularly within the linker region and other solvent-exposed degrader segments.

In contrast, the KRAS-directed degrader system (**42**) proved substantially more challenging. AF3 was unable to predict the allosteric cryptic Switch II binding pocket (**43**), similar to the KRAS ligand alone (**30**), despite correctly recovering the VHL ligand (**44**). Boltz-2 (**Figure S5**) was able to recover the individual KRAS- and VHL-binding ligands, likely because Switch II pocket-binding KRAS ligand structures are represented in the training data; however, it failed to consistently reproduce the correct ternary organization between KRAS and the VHL complex. This may reflect the greater conformational search complexity of this system, in which the flexible linker, compared with the CDK2 PROTAC, spans a more extended distance across a partially solvent-exposed interface. Together, these data suggest that AF3 and Boltz-2 are relatively effective at recovering individual ligand geometry for canonical orthosteric targets and E3 ligase ligands, but remain limited in predicting the correct spatial organization of ternary degrader complexes, particularly when flexible linkers and cryptic or solvent-exposed binding interfaces are involved.

### 6. Performance in Molecular Glue Complexes

Molecular glues are small molecules that induce or stabilize ternary protein complexes by promoting ligand-dependent protein–protein interactions.^51,52^ Unlike PROTACs, which use two bespoke binding ligands connected by a linker to recruit an E3 ligase to a target protein, molecular glues typically act by creating or stabilizing complementary interaction surfaces at the protein–protein interface.^51,53^ Classic FDA-approved examples, such as sirolimus/rapamycin, illustrate this mechanism by stabilizing the FKBP12–mTOR interaction.^54^ More recently, molecular glues have attracted renewed interest as an induced-proximity strategy that may overcome some limitations of PROTACs, including large molecular size, linker-dependent optimization, poor permeability, and challenging drug-like properties.^51^

We curated a small but mechanistically diverse set of recently reported molecular-glue ternary complexes (**Figure 10A-C**, **Figure S6**), including the pan-RAS glue daraxonrasib/RMC-6236 bound to NRAS WT and cyclophilin A (CYPA) (**45**),^55^ the covalent tri-complex formed by RMC-4998, KRAS G12C, and CYPA (**46**),^56^ the intramolecular bivalent glue degrader IBG1 bound to BRD4 and DCAF16:DDB1ΔBPB (**47**),^57^ the molecular glue dHTC1 bound to the ENL YEATS domain and the E3 ligase cereblon (**48**) and the CRBN–DDB1–WIZ zinc finger 7 complex stabilized by dWIZ-1 (**49**).^58,59^ Together, these systems span CypA-recruiting RAS glues, covalent glue mechanisms, intramolecular bivalent glue degraders, and CRBN-mediated neosubstrate recognition. The molecular structures and associated testing data are provided in **Table S7**.

**Figure 10.**
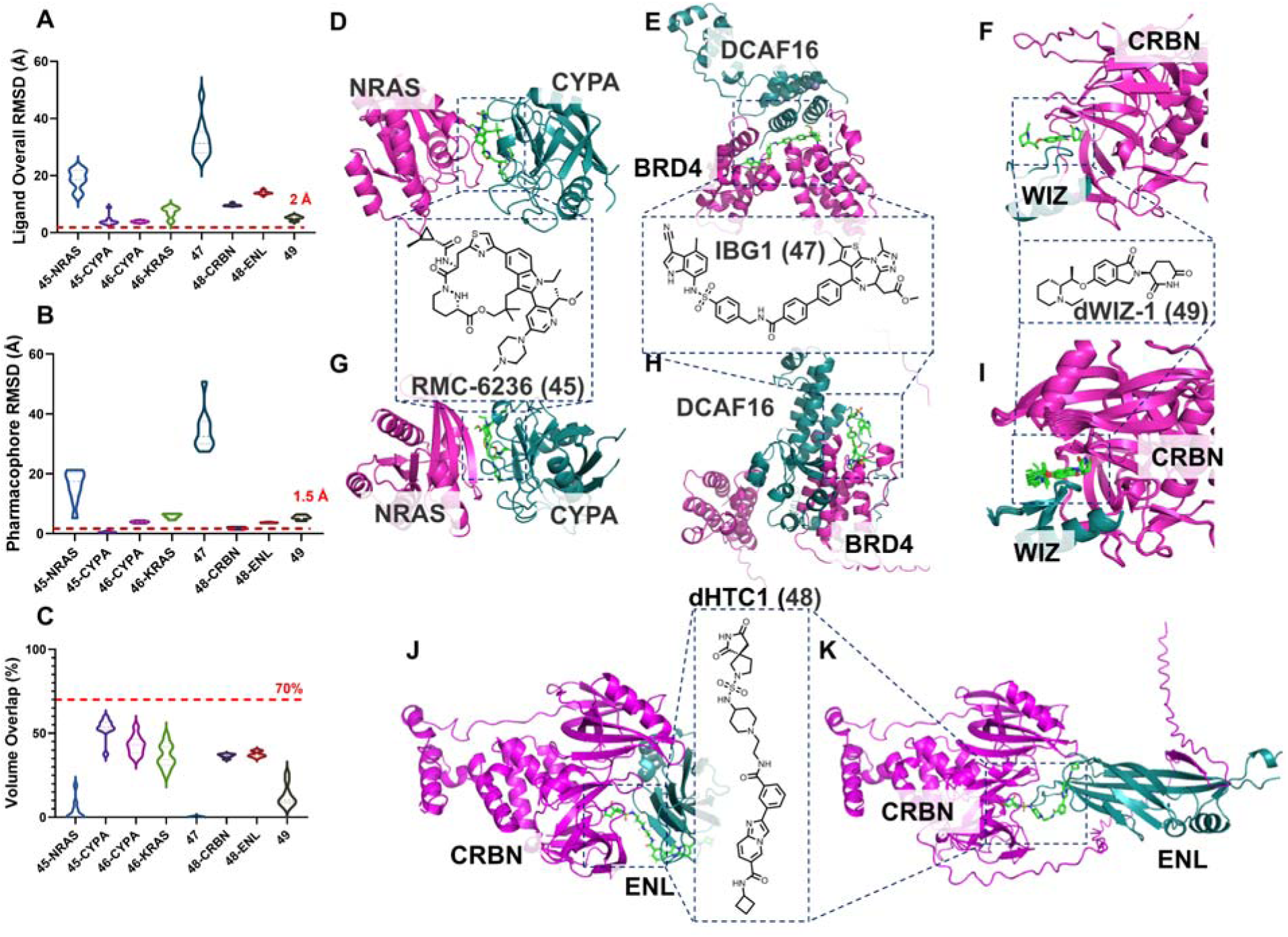
AF3 prediction performance for molecular glue and induced-proximity complexes. (**A**) Ligand overall RMSD. (**B**) Ligand pharmacophore RMSD. (**C**) Ligand volume overlap across the molecular glue dataset (2D chemical structures of representative molecular glues shown in inset). (**D**-**K**) Representative examples of experimentally observed and AF3-predicted molecular glue ternary complexes. (**D**-**F**) Experimental structures of RMC-6236–NRAS–CypA (**45**: 9BG0) complexes, IBG1–BRD4–DCAF16 (**47**: 8OV6) and dWIZ-1–CRBN–WIZ (**49**: 8TZX). (**G**–**I**) Corresponding AF3 predictions for **45**, **47**, and **49**. (**J**-**K**) Experimental and AF3-predicted structures of the CRBN–ENL molecular glue complex (**48**: 9DUR). AF3 showed variable performance across molecular glue systems, ranging from successful recovery of selected CRBN-mediated and hotspot-defined complexes to failures involving induced protein–protein interface organization, ternary assembly geometry, and ligand-mediated interprotein recognition. For entries **46**, the corresponding structure is 8G9P.

Across this dataset, AF3 and Boltz-2 showed a recurring pattern: individual ligand-recognition elements were often partially recovered, whereas the full induced ternary assembly was less reliably predicted. In the RAS–CypA glue systems (**45** and **46**), the models partially captured ligand recognition by CypA and approximate placement of the macrocyclic scaffold within the composite binding region. However, this local recognition did not consistently translate into accurate recovery of the RAS–CypA interface, leading to variability in ligand RMSD, pharmacophore RMSD, volume overlap, and overall ternary geometry (**Figure 10D, G, Figure S6, Figure S28**). Similarly, in the CRBN–ENL complex, the models were able to partially recover ENL-side ligand recognition and imide engagement with CRBN, but failed to accurately reproduce the induced protein–protein interaction between ENL and CRBN (**Figure 10J, K)**. These results suggest that correct recognition of one or both ligand-binding elements is not sufficient when productive molecular-glue activity depends on precise organization of a newly induced interprotein interface. The IBG1–BRD4–DCAF16:DDB1ΔBPB complex was particularly challenging because it requires recovery of a larger higher-order assembly rather than a compact binary interface (**Figure 10E, H**). Both AF3 and Boltz-2 failed to accurately reconstruct the overall protein arrangement, resulting in substantial deviations in ternary-complex geometry.

In contrast, the CRBN–WIZ–dWIZ-1 complex represented a notable success case. dWIZ-1 recruits WIZ zinc finger 7 to CRBN–DDB1, leading to WIZ degradation and fetal hemoglobin induction, a therapeutically relevant strategy for sickle cell disease. AF3 and Boltz-2 successfully captured the induced neosubstrate-recognition geometry between CRBN and WIZ, with ligand RMSD, pharmacophore RMSD, and volume-overlap metrics indicating strong agreement with the experimental structure (**Figure 10F, I**). Structural overlays further showed substantial preservation of the CRBN-mediated recruitment mode. Compared with the other systems, the WIZ component was represented experimentally and computationally as a zinc-finger fragment (30 amino acids), which may reduce the conformational and interaction search space. Nevertheless, the result is significant because neosubstrate recognition is central to molecular-glue pharmacology and suggests that AF3 and Boltz-2 may be useful for selected induced-proximity systems when the recruited interface is compact, well constrained, and supported by a defined ligand-recognition mode.

### 7. Performance in Targeted Covalent Inhibitors

Targeted covalent inhibitors have reemerged as an important drugging modality because they combine reversible molecular recognition with selective covalent engagement of a strategically positioned nucleophilic residue.^60^ This mechanism can improve potency, extend target residence time, and sustain pharmacodynamic activity beyond the plasma clearance of the parent compound.^60^ However, covalent inhibitor modeling requires more than recovery of the overall ligand pose: accurate prediction must also place the electrophilic warhead in the correct orientation relative to the target nucleophile.^61^

We curated a set of recently reported targeted covalent inhibitor complexes that span viral proteases and oncogenic KRAS mutants (**Figure 11A-C**). The set includes SARS-CoV-2 PLpro complex **50** (**Figure 11D**),^33^which represents a PLpro inhibitor binding near the ubiquitin-binding/BL2-groove region; KRAS G12C complexes **51** (**Figure 11E**) and **52** (**Figure 11F**) with BBO-8520, a direct covalent dual inhibitor designed to engage both GDP-bound and active nucleotide-bound KRAS G12C states through the Switch-II/Helix-3 pocket;^62^ SARS-CoV-2 main protease complex **53** (**Figure 11G**) with PF-07817883, a second-generation orally bioavailable Mpro inhibitor developed to improve metabolic stability relative to nirmatrelvir;^63^ and KRAS G12D complex **54** (**Figure 11H**) with YK-8S, an oxirane-based covalent ligand reported to engage both KRAS G12D and KRAS G12C mutants.^64^ We also considered covalent examples discussed in other sections, including the allosteric WRN inhibitor VVD-214 **27** (**Figure 11I**) and the covalent RAS molecular-glue complex RMC-4998–KRAS G12C–CypA **46**. Together, these structures cover conventional active-site covalent inhibitors, covalent KRAS switch-pocket ligands, covalent allosteric inhibitors, and covalent induced-proximity complexes.

**Figure 11.**
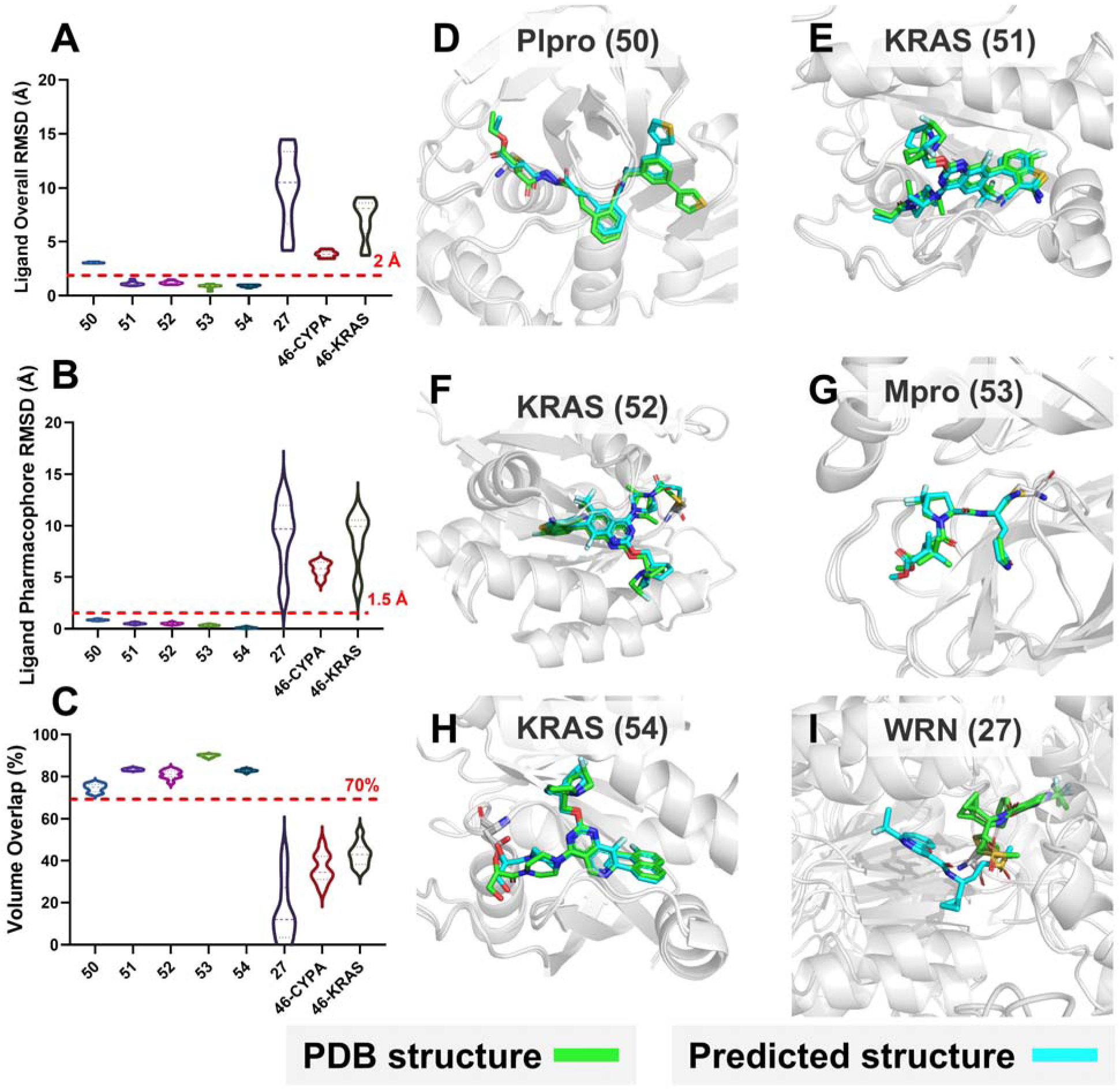
AF3 prediction performance for covalent inhibitor complexes. (**A**–**C**) Ligand overall RMSD, pharmacophore RMSD, and volume overlap across the covalent inhibitor dataset. Dashed red lines mark chemically useful pose-recovery thresholds. (**D-H**) The dataset includes SARS-CoV-2 PLpro (**50**: 8UVM), KRAS G12C–BBO-8520 complexes (**51**: 8V3A; **52**: 8V39), SARS-CoV-2 Mpro–PF-07817883 (**53**: 8V4U), KRAS G12D–YK-8S (**54**: 8JHL), and the (**I**) WRN–VVD-214 (**27**: 7GQU).

Overall, AF3 and Boltz-2 (**Figure S7**) showed strong pose recovery across this category, with most complexes showing low RMSD values, including several near or below 1–1.5 Å. These results suggest that covalent ligands can be accurately modeled when binding is anchored by a well-defined pocket and a strong recognition pharmacophore. The failures were more informative than random. The more difficult cases involved either incorrect ligand orientation, allosteric pocket formation, macrocyclic conformational sampling, or ternary-complex organization. Thus, covalent chemistry itself does not appear to be the dominant limitation. Instead, AF3 and Boltz-2 performance depends on whether the covalent ligand is presented in a structurally constrained binding site. The molecular structures and associated testing data are provided in **Table S8**.

The SARS-CoV-2 PLpro complex **50** provided a particularly informative example because overall ligand placement was recovered with high accuracy by both AF3 and Boltz-2, while the geometry of the covalent bond remained substantially distorted relative to the experimental structure. In covalent cysteine adducts, the thioether geometry surrounding the sulfur atom typically adopts an approximately tetrahedral arrangement, with C–S–C bond angles commonly near ∼101–109°, depending on local chemical environment and hybridization state. In the experimental 8UVM structure, the observed covalent geometry exhibited a bond angle of approximately 130.8°. In contrast, AF3 predictions produced substantially expanded geometries, with an average angle of approximately 162.2° across ten models, indicating a nearly linearized sulfur linkage inconsistent with the experimentally observed covalent arrangement. Boltz-2 generated comparatively improved geometries, with an average angle of approximately 125.8°, although these values still remained systematically larger than the experimental structure.

These observations suggest that current AI-based structure prediction methods can successfully recover the global binding mode of covalent inhibitors while still failing to reproduce chemically realistic local covalent geometry. This limitation is mechanistically plausible because AF3 and Boltz-2 primarily optimize structural consistency and learned spatial relationships rather than explicitly modeling electronic structure, orbital geometry, or transition-state chemistry associated with covalent bond formation. Similar limitations have been noted in docking and AI-based covalent modeling workflows, where accurate global pose recovery does not necessarily guarantee chemically optimal warhead orientation or reaction geometry. Consequently, evaluation of covalent inhibitor predictions should include explicit inspection of local bond geometry in addition to conventional RMSD-based pose metrics.

More broadly, these results indicate that covalent inhibitor prediction by AF3 and Boltz-2 is feasible when the surrounding recognition environment is structurally constrained, but chemically accurate modeling of the reactive center remains an unresolved challenge for current AI-based structural prediction approaches.

### 8. Performance in Membrane Proteins

Membrane proteins represent a challenging class for structure-based modeling because ligand recognition often depends on the native bilayer environment, transporter or receptor conformational state, and lipid–protein interactions.^65^ They are also experimentally difficult targets, requiring specialized purification, detergent or nanodisc reconstitution, and cryo-EM or crystallographic workflows.^66^ To evaluate this modality, we selected three recent membrane-protein ligand complexes: MCT8 bound to silychristin (**55**),^67^ OATP1B1 bound to atorvastatin (**56**),^68^ and GAT3 bound to GABA (**57**).^69^

Across these examples, AF3 (**Figure 12A-C**) and Boltz-2 (**Figure S8**) showed target-dependent performance rather than uniformly reliable ligand-pose prediction. The most severe failure was observed for **55**, where the predicted protein architecture was substantially incorrect. The molecular structures and associated testing data are provided in **Table S9**. In this case, ligand-pose evaluation becomes difficult to interpret because the binding-site environment itself was not recovered. Thus, the high ligand RMSD and poor overlap likely reflect failure of membrane-protein fold or conformational-state prediction rather than ligand placement alone.

**Figure 12.**
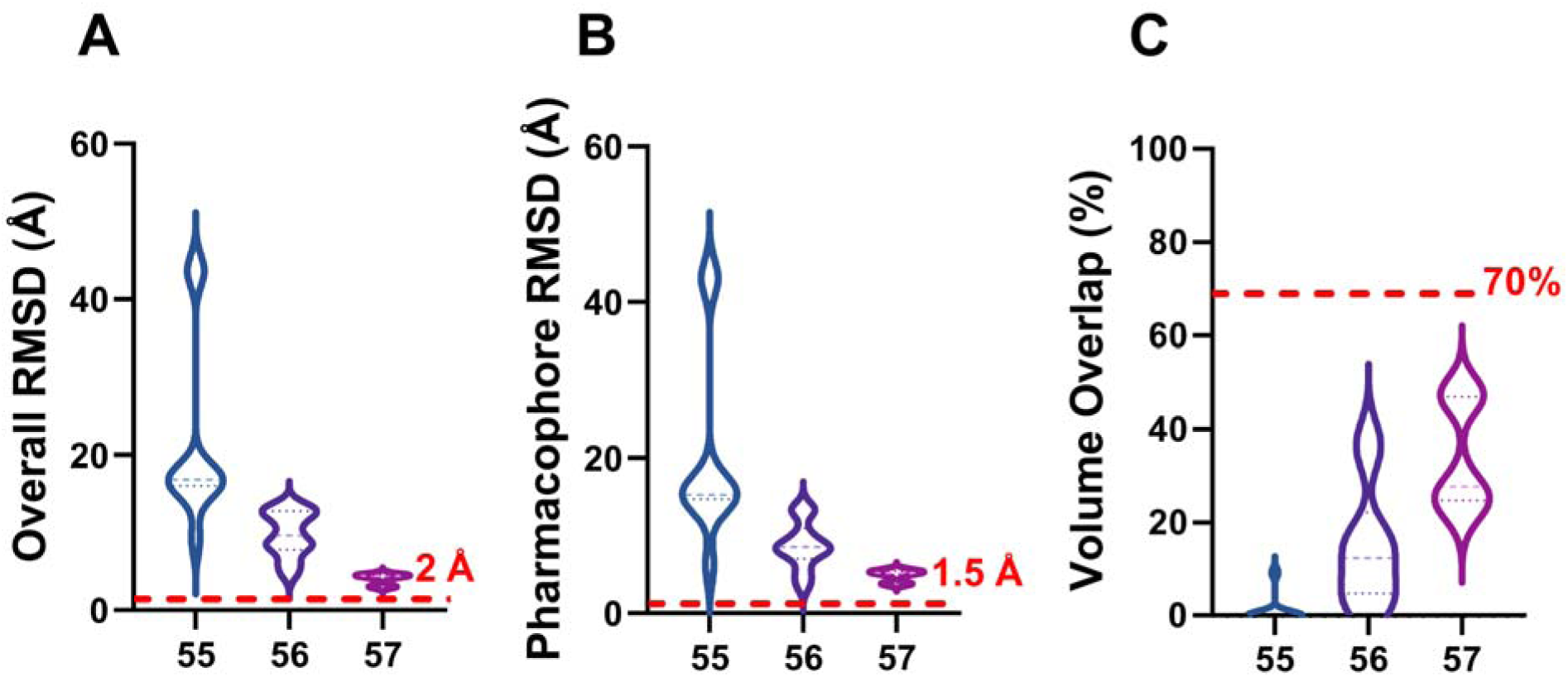
AF3 prediction analysis of membrane protein ligand complexes. (**A–C**) Ligand overall RMSD, pharmacophore RMSD, and volume overlap for membrane-protein systems. Prediction performance varied substantially across the dataset, with poor recovery for MCT8–silychristin (**55**: 8ZKN), intermediate performance for OATP1B1–atorvastatin (**56**: 9CY3), and relatively better recovery of the GAT3–GABA complex (**57**: 9LK8), highlighting the challenges associated with membrane-dependent pocket geometry and conformational state prediction.

In contrast, **56** showed partial recovery of the transporter architecture. Although the overall ligand metrics were not optimal, both models captured the central translocation channel and placed the ligand near the experimentally observed binding region. This suggests that AF3 and Boltz-2 can sometimes identify the correct membrane-protein binding cavity even when precise ligand orientation and local interaction geometry are imperfect.

The **57** GAT3–GABA complex represented the most favorable case. Both AF3 and Boltz-2 recovered the transmembrane protein architecture and identified the small-molecule binding pocket. Boltz-2 showed more accurate local ligand placement, whereas AF3 also localized GABA to the correct region but with greater conformational dispersion. The better performance for 9LK8 may reflect the small size and limited flexibility of GABA, together with a compact and well-defined substrate-binding cavity.

Together, these results indicate that membrane-protein ligand prediction remains highly sensitive to recovery of the correct protein fold and conformational state. When the transmembrane architecture or channel geometry is correctly modeled, AF3 and Boltz-2 may localize the ligand near the correct binding pocket, but accurate pose recovery remains challenging without explicit membrane context.

### 9. Performance in RNA Binders

RNA binders present a distinct challenge because ligand recognition depends on RNA secondary and tertiary features, including bulges, internal loops, mismatches, repeat motifs, base stacking, groove shape, and ion-dependent folding.^70^ The approval of the splicing modifier risdiplam for spinal muscular atrophy established RNA as a clinically validated small-molecule target,^71^ while emerging RIBOTAC strategies further extend RNA binding from occupancy-driven recognition to induced RNA degradation.^72^

We evaluated AF3 (**Figure 13A-C**) and Boltz-2 (**Figure S9**) using recent RNA–ligand complexes, including risdiplam bound to the SMN2/U1 snRNA splice-site RNA helix (**58**),^73^ ANP77 bound to C9orf72-related G2C4 repeat RNA (**59**),^74^ SMN-CX bound to a splice-site RNA helix (**60**),^73^ naphthyridine-azaquinolone bound to an RNA ACG/AUA motif (**61**),^75^ disease-associated r(CUG) repeat RNA binders (**62**),^76^ and an additional 2025 RNA–ligand complex (**63**).^77^ Pose recovery was variable, with RMSD values ranging from accurate predictions such as **58** at 1.73 Å and **60** at 2.35 Å to failed cases such as **59** at 16.73 Å and 8ZNQ at 10.88 Å. These results suggest that RNA binders remain challenging, particularly when ligand recognition depends on flexible RNA motifs, repeat structures, or local RNA remodeling. The molecular structures and associated testing data are provided in **Table S10**.

**Figure 13.**
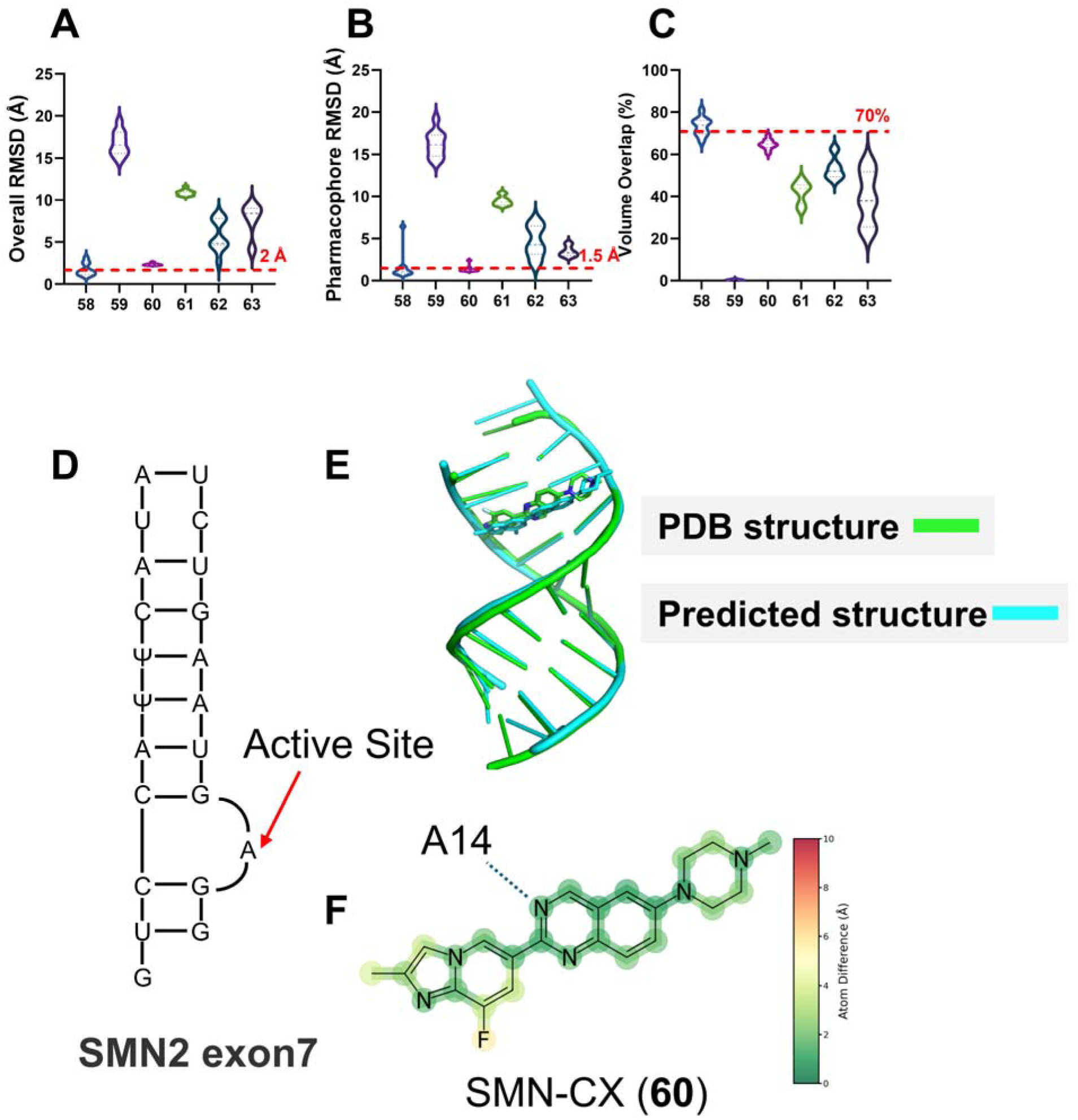
AF3 prediction performance for RNA-targeting small molecules. (**A–C**) Ligand overall RMSD, pharmacophore RMSD, and volume overlap across the RNA dataset. (**D**) Representative analysis of the SMN-CX splice-site RNA complex (**60**: 8R8P). (**E**) AF3 recovered the overall RNA fold and preserved key ligand recognition features, including the interaction centered on nucleotide A14. (F) The corresponding 2D interaction map illustrates per-atom prediction accuracy and predicted ligand–RNA interaction geometry. For entries **58**, **59**, **61**, **62** and **63**, the corresponding structures are 8R62, 8QMH, 8ZNQ, 9CPD and 9IO0.

The splice-site RNA complexes were among the better predicted examples. For SMN-CX (**60**; **Figure 13D-F**), AF3 reproduced the overall RNA architecture and placed the ligand close to the experimental pose while retaining the aromatic stacking interaction near A14. Boltz-2 showed greater variation in ligand orientation but still recovered the central recognition region. Similar performance for **58** suggests that relatively well-defined splice-site helices are more amenable to current prediction methods.

In contrast, substantially larger errors were observed for several repeat-RNA and noncanonical motif systems. In these cases, incorrect local RNA geometry was accompanied by poor ligand RMSD, pharmacophore RMSD, and volume overlap. Repeat RNAs and internal-loop structures can sample multiple local conformations, so relatively small differences in base orientation or groove geometry may lead to much larger differences in ligand placement.

The RNA examples therefore do not show a uniform failure of AF3 or Boltz-2. Rather, prediction was more successful for structurally constrained RNA binding sites and less reliable when ligand recognition depended on flexible or heterogeneous local RNA conformations.

### 10. Performance on Fragment-Sized Ligands in Early Discovery

We next evaluated fragment-sized ligands, which are central to fragment-based drug discovery but pose a distinct challenge for structure prediction (**Figure 14A-C**, **Figure S10**). Because fragments are small and often bind with weak affinity, they provide fewer interaction anchors than lead-like molecules and may occupy shallow, partially formed, or multiple possible sites. Accurate prediction therefore requires both correct binding-site identification and recognition of a minimal pharmacophore, rather than simply optimizing a well-defined ligand pose within a known pocket (**Figure 14D**). The molecular structures and associated testing data are provided in **Table S11**.

**Figure 14.**
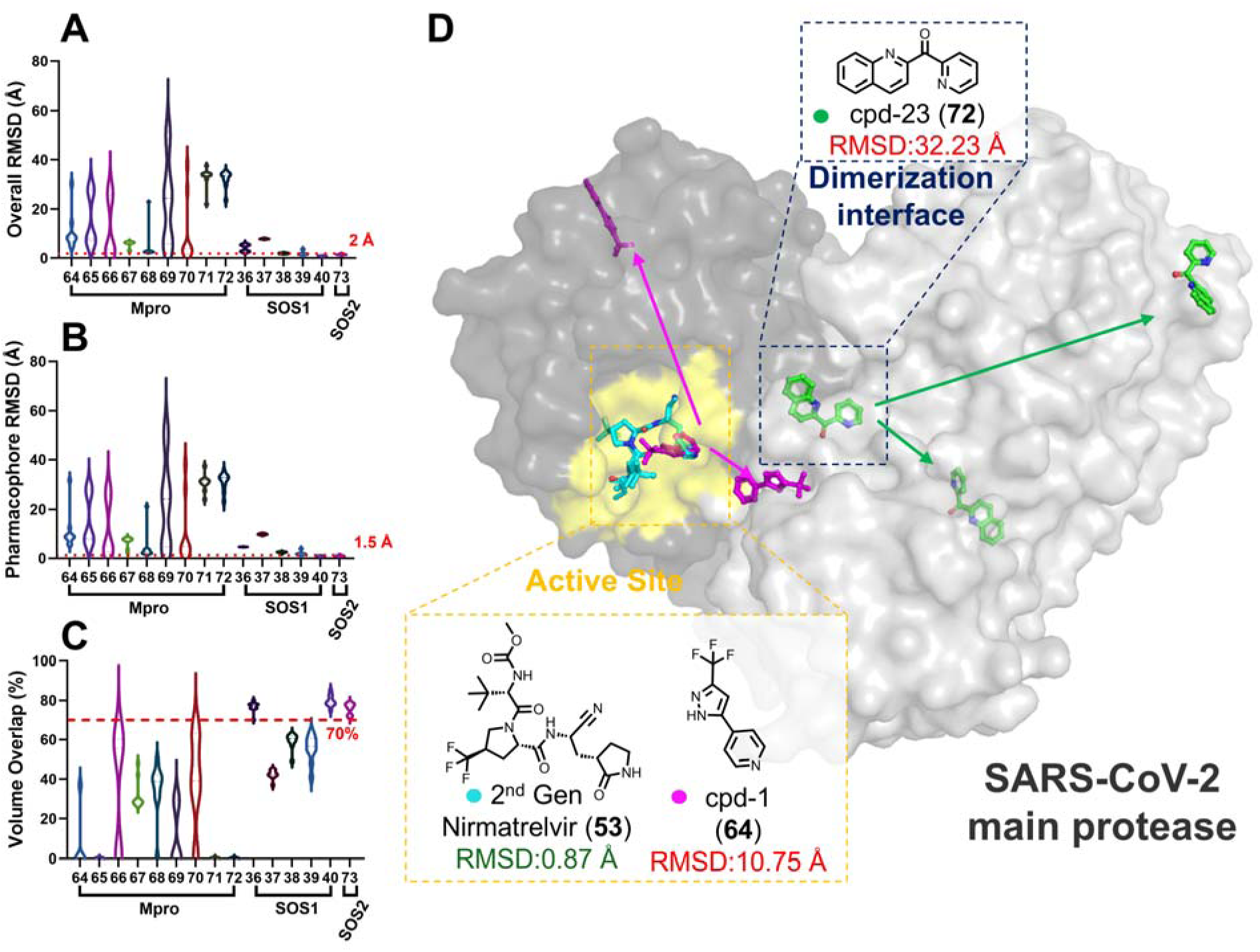
(**A-C**) Ligand overall RMSD; Ligand pharmacophore RMSD and ligand volume overlap. (**D**) Representative AF3 predictions for SARS-CoV-2 main protease (Mpro) complex with fragments. Experimental Mpro structures containing the active-site inhibitor nirmatrelvir derivative (**53**: 8V4U, cyan), the fragment ligand (**64**: 7GRE, magenta), and the dimerization-interface ligand cpd-23 (**72**: 7GS0, green) are superimposed with the corresponding AF3-predicted structure (orange).^78^ Ligands are indicated by arrows. AF3 accurately recovered ligand placement within the canonical catalytic pocket but showed reduced accuracy for ligands targeting the shallow dimerization interface, highlighting differences in prediction performance between orthosteric active-site recognition and protein–protein interface binding. For entries **36**-**40**, **65**-**71** and **73**, the corresponding structures are 8T5G, 8T5M, 8T5R, 8UC9, 8UH0, 7GRF 7GRJ, 7GRN, 7GRS, 7GRT, 7GRU, 7GRZ and 9BVE.

We focused on two fragment-rich systems with high chemical diversity: SARS-CoV-2 main protease and SOS1- and SOS2–KRAS PPI fragments (**Figure 14A-C**). The main protease was modeled as a dimer, creating multiple potential ligandable regions, including active-site and interfacial sites, which increased the difficulty of assigning the correct fragment-binding location and pose (**64**, **72** in **Figure 14D**). In contrast, SOS1- and SOS2–KRAS PPI fragments were predicted with high accuracy and preferentially localized to the SOS1/2–KRAS PPI site, consistent with the strong performance observed for this interface in earlier analyses. These results suggest that AF3 and Boltz-2 can be useful for fragment pose generation when the binding site contains a defined pharmacophore-rich pocket, but performance may decrease when small fragments have multiple plausible binding regions or insufficient interaction density to constrain the prediction. Because the SOS1- and SOS2–KRAS PPI fragments occupy a recurrent and consistently defined pocket across related experimental structures, the models may benefit from a more strongly constrained recognition environment, which could partly explain their higher prediction accuracy for this system.

### 11. Model-Derived Confidence Metrics for Ligand Pose Triage

We next examined whether AF3 confidence metrics could predict ligand-pose quality across the dataset of 83 complexes. We compared AF3 minPAE and ipTM, as well as Boltz-2 ipTM, with overall ligand RMSD. AF3 minPAE showed a moderate positive correlation with overall RMSD (Pearson r = 0.665) and a stronger Spearman correlation (r = 0.758), indicating that higher minPAE values were generally associated with less accurate poses (**Figure S11**). AF3 and Boltz-2 ipTM were negatively correlated with overall RMSD, with stronger correlations by Spearman than by Pearson analysis. We then performed the same analysis within individual ligand-binding modalities, but no subclass showed a consistently strong relationship between the confidence scores and overall RMSD (**Figures S12–S21**). Although these confidence metrics were associated with pose accuracy across the full dataset, the relationships varied across ligand-binding modalities.

Because the correlations between confidence scores and pose accuracy were not uniform across the dataset, we next grouped minPAE and ipTM values into score ranges to determine whether particular ranges were enriched for accurate poses (**Figure 15A-C, Figure S23-S25**). Low AF3 minPAE values showed a clear enrichment for pharmacophore recovery (<1.5 Å): 100% (22/22) for minPAE <0.85 Å, 72.7% (8/11) for 0.85–1.0 Å, 61.5% (8/13) for 1.0–1.5 Å, 42.9% (3/7) for 1.5–2.0 Å, and 3.3% (1/30) for >2.0 Å. A similar trend was observed for overall RMSD, with accurate poses found in 81.8% (18/22), 45.5% (5/11), 30.8% (4/13), 14.3% (1/7), and 6.7% (2/30) of predictions across the same minPAE ranges, respectively. Higher minPAE values also more often corresponded to larger ligand displacement, poorer pharmacophore alignment, and reduced volume overlap.

**Figure 15.**
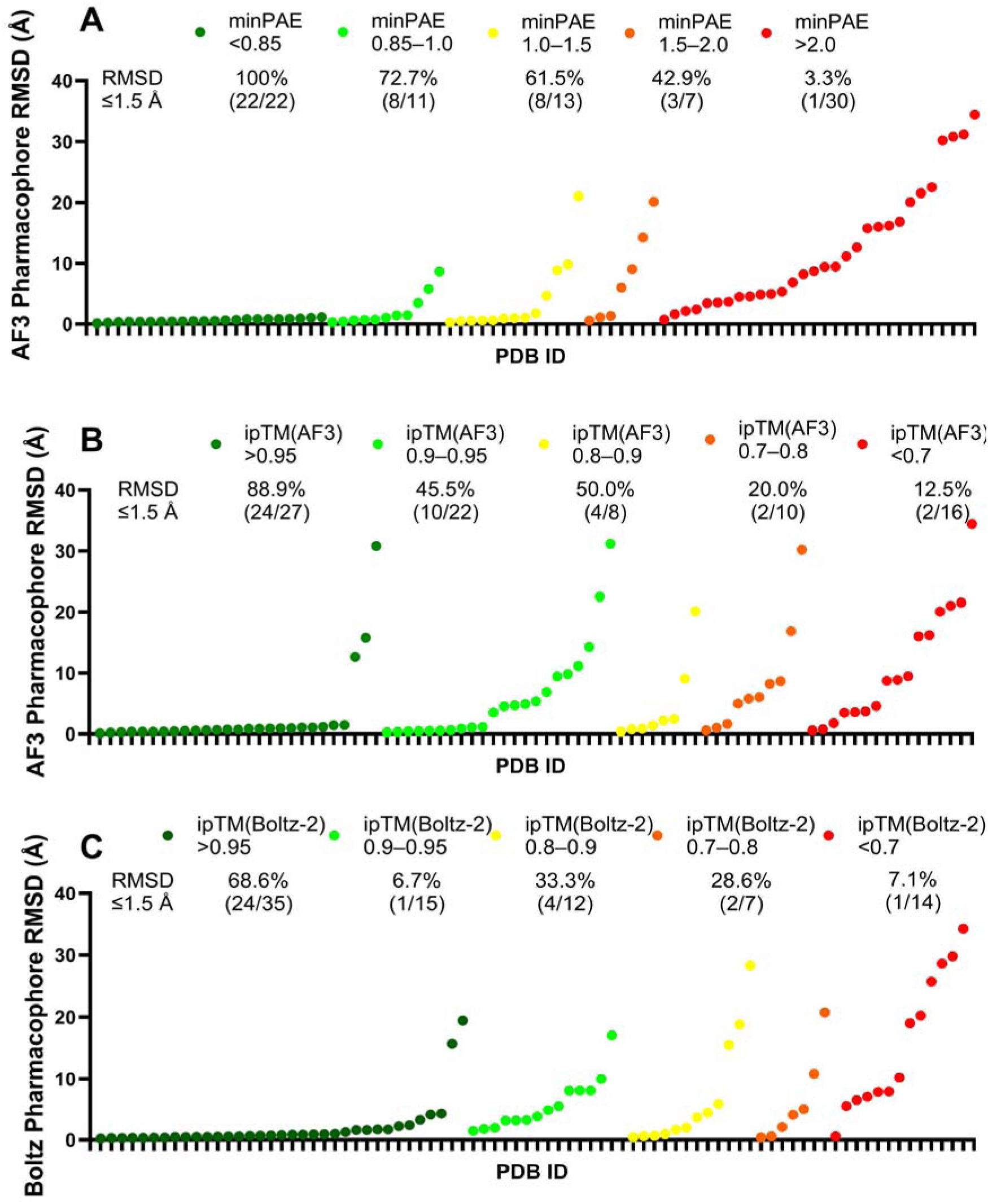
Relationship between model-reported confidence metrics and ligand pharmacophore RMSD. (**A**) AF3 protein–ligand minimum predicted aligned error (minPAE) plotted against ligand pharmacophore RMSD, with points grouped by minPAE ranges. (**B**) AF3 ipTM plotted against ligand pharmacophore RMSD, with points grouped by ipTM ranges. (**C**) Boltz-2 ipTM plotted against ligand pharmacophore RMSD, with points grouped by ipTM ranges. Lower minPAE values show stronger enrichment for low pharmacophore RMSD poses, whereas ipTM values from AF3 or Boltz-2 show weaker separation between accurate and inaccurate ligand poses.

We next examined whether low minPAE scores were preferentially associated with particular ligand-binding modalities. The fraction of predictions with minPAE <0.85 Å was 0% for sEH, 73.7% for canonical orthosteric ligands, 28.6% for allosteric ligands, 11.1% for PPIs, 10.0% for PROTACs, 0% for molecular glues, 62.5% for covalent ligands, and 0% for membrane proteins, RNA-binding ligands, fragments, and E3 ligase ligands (**Figure S22**). Canonical orthosteric ligands, including both noncovalent and covalent examples, therefore showed the highest representation in the minPAE <0.85 Å group. Interestingly, none of the in-house sEH inhibitors had a minPAE below 0.85 Å despite generally strong pose recovery. This may reflect the large and flexible lipid-binding pocket of sEH, which can accommodate multiple plausible ligand configurations. More challenging systems, including allosteric pockets, membrane proteins, flexible RNA motifs, and large induced-proximity assemblies, generally showed fewer predictions in the minPAE <0.85 Å group and a broader distribution of minPAE values.

Grouping of AF3 ipTM scores showed a similar, although somewhat weaker, enrichment. The fraction of predictions with pharmacophore RMSD <1.5 Å was 88.9% (24/27) for ipTM >0.95, 45.5% (10/22) for 0.90–0.95, 50.0% (4/8) for 0.80–0.90, 20.0% (2/10) for 0.70–0.80, and 12.5% (2/16) for <0.70. This trend was less consistent than that observed for minPAE. Boltz-2 ipTM showed still weaker separation: 68.6% (24/35) for >0.95, 6.7% (1/15) for 0.90–0.95, 33.3% (4/12) for 0.80–0.90, 28.6% (2/7) for 0.70–0.80, and 7.1% (1/14) for <0.70.

Overall, these results suggest that, within our dataset, certain confidence-score ranges can substantially enrich for accurate ligand poses, although the strength of the relationship varies across ligand-binding modalities. AF3 minPAE was the most useful metric examined, with minPAE <0.85 Å strongly enriching for accurate pose and pharmacophore recovery. However, neither minPAE nor ipTM alone reliably identifies chemically realistic ligand poses across all target classes. Visual inspection, pharmacophore analysis, and local chemical-geometry validation therefore remain important before predicted structures are used for medicinal chemistry interpretation or compound-design decisions.

### 12. Ligand Activity Ranking

Having established that minPAE can enrich for accurate poses, we next examined whether model-derived scores, predicted binding affinity, or pose-quality metrics were associated with experimental activity. We first examined six sEH inhibitors using the Boltz-2 binding-affinity module (**Figure 16A, B**). Because all six compounds were highly active and spanned a narrow experimental IC_50_ range (0.009–0.115 μM), this small series was not intended as a definitive affinity benchmark. The predicted AF3 minPAE scores and Boltz-2 affinities did not reproduce the experimental activity ranking within this series. In addition, predicted affinity showed no clear association with pose-quality metrics by either Pearson or Spearman analysis, including overall ligand RMSD, pharmacophore RMSD, or molecular volume overlap.

**Figure 16.**
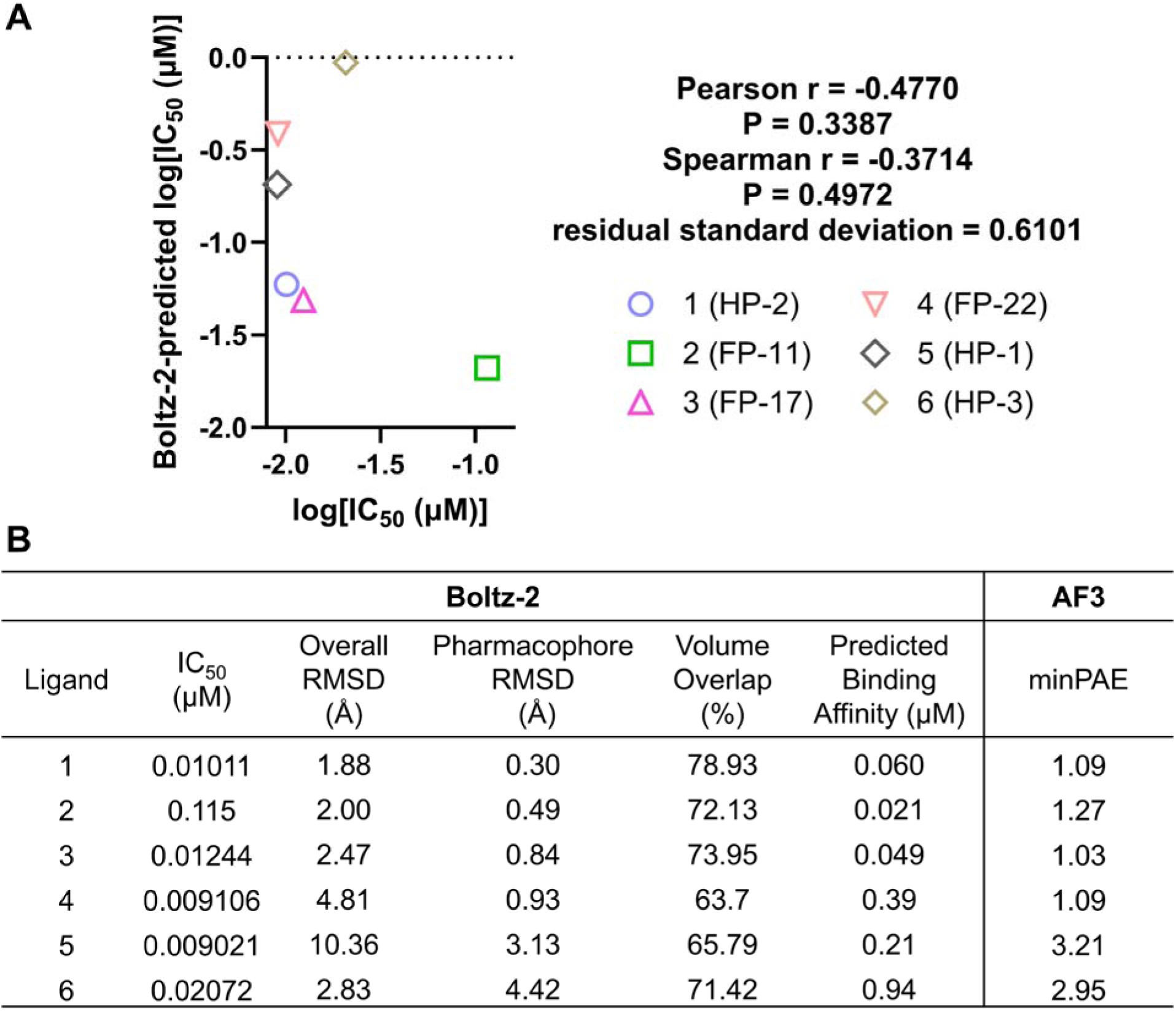
Relationship between experimental activity, AI-derived predictions, and pose-recovery metrics for the sEH inhibitor series. (**A**) Comparison of experimental potency log_10_[IC_50_ (μM)] with Boltz-2-predicted binding affinity for six sEH inhibitors. (**B**) Experimental IC_50_ values, Boltz-2-predicted binding affinities, AF3 minPAE values, ligand overall RMSD, pharmacophore RMSD, and volume overlap for the same inhibitor series. Additional scatter plots comparing experimental IC_50_ values with the other predictions and pose-recovery metrics listed in panel B are shown in **Figure S26**. Overall, Boltz-2-predicted affinity does not correlate well with measured biochemical activity, despite several compounds showing accurate pose recovery by RMSD and volume-overlap metrics.

Given the small size and narrow activity range of the sEH series, we next evaluated Boltz-2 on a larger external dataset consisting of 20 SARS-CoV-2 Mpro–ligand complexes with reported biochemical activity and experimentally determined X-ray structures from a single study to minimize assay variability (**Figure 17**). The predicted ligand poses generally agreed well with the experimental structures, showing low ligand RMSD values and high molecular overlap. In this dataset, Boltz-2 predicted affinity correlated significantly with experimental activity (Pearson r = 0.742, P = 0.0002; Spearman r = 0.695, P = 0.0007). However, the residual standard deviation of 0.63 log units indicates substantial compound-level variability around this trend. We also tested whether minPAE correlated with experimental activity, but no clear correlation was observed (**Figure S27**). Thus, while Boltz-2 affinity predictions captured relative activity trends in this dataset, their precision was limited for quantitative potency prediction. Together, the sEH and Mpro datasets suggest that Boltz-2 can capture activity trends in some ligand series, but its performance varies across chemical systems. Overall, these predictions appear more useful for compound prioritization than for estimating absolute potency.

**Figure 17.**
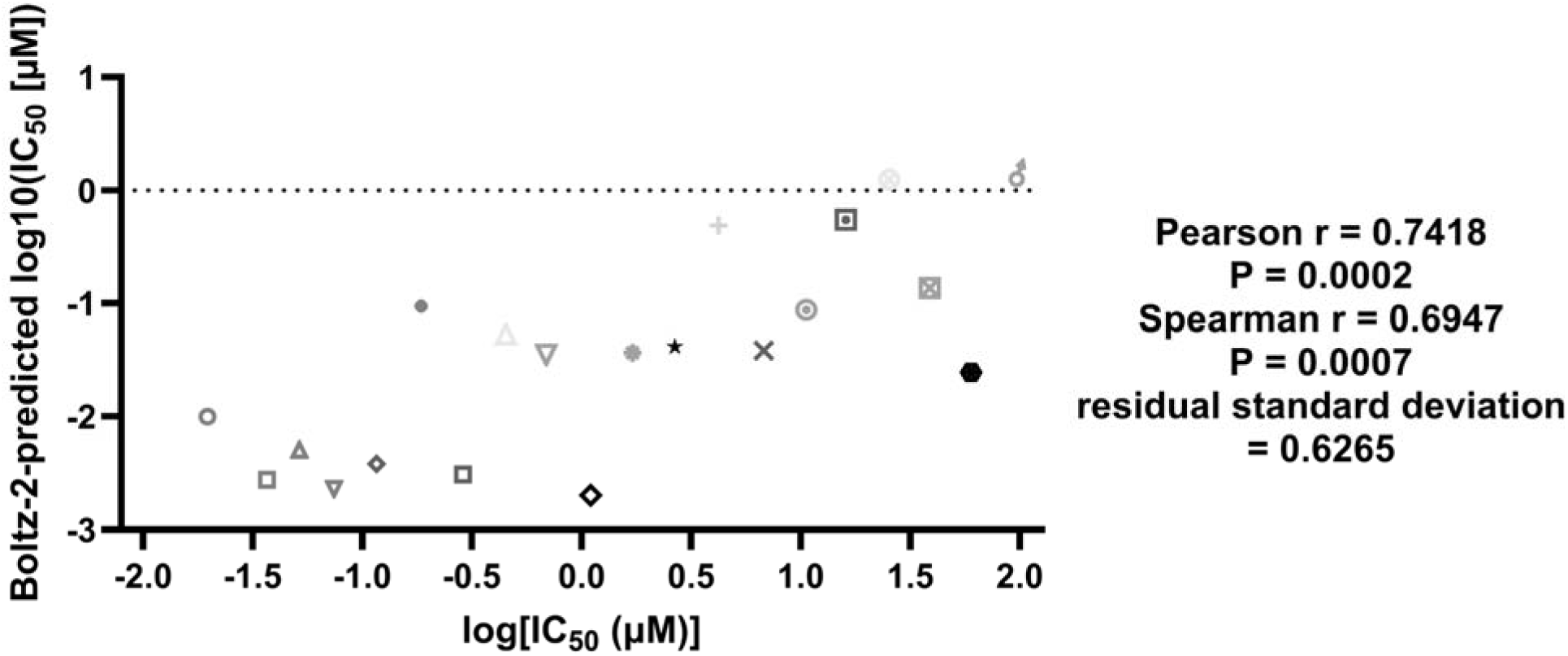
Comparison of experimental activity and Boltz-2-predicted binding affinity for 20 SARS-CoV-2 Mpro inhibitors. Experimental log10[IC50 (μM)] is plotted on the x-axis and Boltz-2-predicted log10[IC50 (μM)] on the y-axis for 20 Mpro inhibitors. For entries **76–95**, the corresponding structures are 7GLV, 7GAW, 7GKS, 7GNT, 7GLG, 7GIU, 7GM1, 7GNA, 7GIX, 7GHM, 7GG2, 7GMR, 7GE3, 7GMP, 7GE2, 7GE4, 7GGC, 7GFJ, 7GCK, and 7GBV, respectively. Entries were arranged in ascending order according to their experimental IC_50_ values, with log_10_[IC_50_ (μM)] values ranging from **−1.70 to 1.98**. Experimental and predicted activity values are listed in **Table S13**.

### 13. Performance on Activity Cliff Pairs

To further examine whether the models could separate active compounds from closely related inactive analogs, we examined an activity-cliff pair, S-thalidomide and N-methyl-S-thalidomide (**Figure 18A, 18B**), to determine whether the models could distinguish a positive ligand from a closely related inactive analog when the activity separation was large. S-Thalidomide is a well-characterized CRBN ligand and is likely represented in the structural training landscape, and both models accurately reproduced its CRBN-bound pose.^79^ In contrast, N-methyl-S-thalidomide was included as a near-analog negative control because glutarimide N-methylation is expected to disrupt productive CRBN engagement.^80,81^ The molecular structures and associated testing data are provided in **Table S12**.

**Figure 18.**
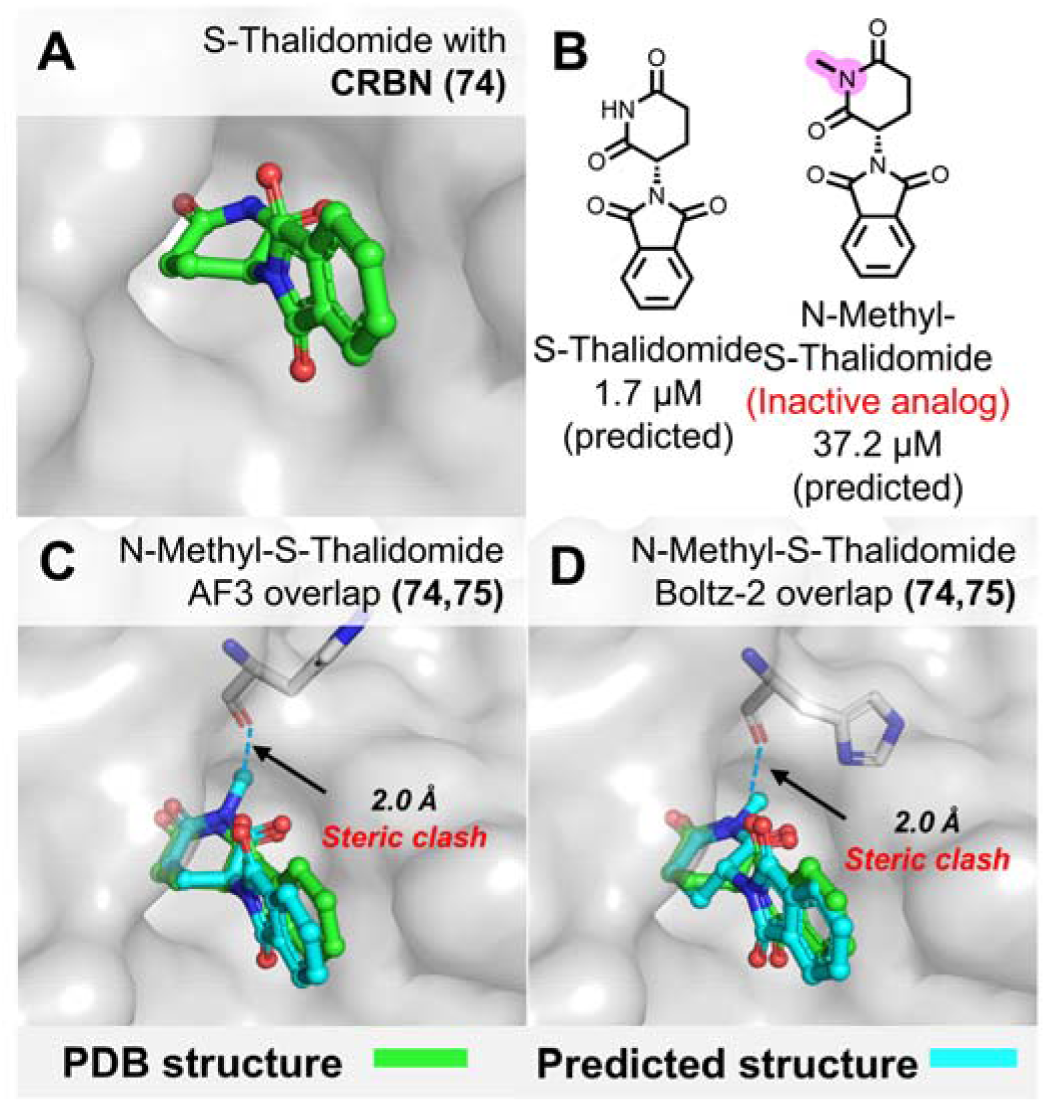
AF3 and Boltz-2 prediction of cereblon (CRBN) ligand recognition and an inactive analog. (**A**) Experimental structure of S-thalidomide bound to CRBN (PDB: 4CI1). (**B**) Chemical structures and predicted binding affinities of S-thalidomide and the inactive analog *N*-methyl-S-thalidomide. (**C, D**) AF3- (C) and Boltz-2-predicted (D) poses of *N*-methyl-S-thalidomide overlaid with the experimental S-thalidomide-bound structure. Despite the loss of activity upon *N*-methylation, both models place the inactive analog in the CRBN binding pocket and additionally generate an ∼2.0 Å interatomic contact consistent with a steric clash.

The expected outcome for N-methyl-S-thalidomide is therefore loss of productive binding. However, both AF3 and Boltz-2 still generated plausible-looking CRBN-bound poses for the inactive analog. Notably, the additional methyl group in N-methyl-S-thalidomide was placed in a sterically unfavorable position, with interatomic distances shorter than the summed van der Waals radii of nearby atoms (**Figure 18C, 18D**). Thus, although the overall pose appeared reasonable, the local binding geometry contained an obvious steric clash. This illustrates an important limitation of AF3 and Boltz-2 pose prediction: the models can generate plausible binding modes for inactive compounds, while local chemical incompatibility may not be fully captured. AF3 minPAE increased from 0.91 to 1.64 Å for the inactive analog, suggesting some separation, but not enough to serve as a reliable triage criterion. In contrast, the Boltz-2 ipTM score remained nearly unchanged (0.99 versus 0.98).

Interestingly, the Boltz-2 binding-affinity module partially captured the activity cliff, predicting stronger binding for thalidomide than for N-methyl-S-thalidomide. Specifically, Boltz-2 predicted an affinity of 1.7 μM for thalidomide compared with 37.2 μM for N-methyl-S-thalidomide, providing some ability to prioritize the active ligand over the inactive analog. Together, this example is consistent with the activity-ranking results above: Boltz-2 affinity predictions can provide useful separation in some cases, but pose prediction alone does not determine whether a ligand will bind productively. These results emphasize that predicted structures require chemical evaluation, particularly for close analogs where small modifications can disrupt binding.

### 14. Practical Chemical Issues in AF3 and Boltz-2 Predictions

Beyond pose prediction accuracy, we observed several chemical-structure and pose generation issues in AF3 and Boltz-2 predictions. Similar stereochemical and ligand-geometry limitations have recently been reported for AF3 and Boltz-family protein–ligand predictions, including errors in chirality, bond lengths, bond angles, and ligand conformations.^82^ A predicted pose can therefore appear reasonable at the protein–ligand complex level while containing chemically unrealistic ligand geometry (**Figure 19A-F**). Before predicted structures are used for SAR analysis or compound design, ligand stereochemistry, bond order and length, alkene planarity, alkyne linearity, ring conformation, and protein–ligand clashes should be inspected. When such errors are present, the ligand geometry should be corrected using appropriate physics-based structure-refinement tools.

- **sp³ center stereochemistry errors**. Both AF3 and Boltz-2 generated incorrect stereochemistry at tetrahedral centers (**Figure 19A**). In the 1W6K example,^83^ the predicted ligand was inverted from the reference R configuration to the S configuration.
- **Double-bond stereochemistry errors**. AF3 can incorrectly assign or distort alkene stereochemistry. In the 8BPV example (**Figure 19B**),^24^ the predicted ligand showed incorrect E/Z stereochemistry and bond length. In addition, both AF3 and Boltz-2 failed to maintain the planar geometry of the alkene.
- **Triple-bond geometry errors**. AF3 failed to preserve the expected linear geometry of the alkyne in the 4AV4 example (**Figure 19C**),^84^ whereas Boltz-2 performed better in this case.
- **Bond-order and ring-geometry distortion**. Incorrect treatment of bond order can also produce larger ligand-geometry distortions. In the example of 7X1T (**Figure 19D**),^85^ The predicted structure showed altered ring geometry and planarity.
- **Single-bond length errors**. Both AF3 and Boltz-2 showed incorrect single-bond lengths in the 8BPV example (**Figure 19B**).
- **Bond angle errors for covalent inhibitors**. Both AF3 and Boltz-2 showed inaccuracies in local covalent-bond geometry despite recovering reasonable overall ligand poses. In the covalent PLpro inhibitor complex **50** (**Figure 19E**), the predicted sulfur linkage showed a bond angle of 161.7°, compared with 130.8° in the experimental structure, with AF3 showing the larger deviation.
- **Steric clash poses.** Beyond these chemical issues, we also observed physically unfavorable poses in which the predicted ligand–protein distances were shorter than the sum of their van der Waals radii, resulting in steric clashes. This was observed for AF3 in the KRAS example (8QVU; **Figure 19F)**, whereas Boltz-2 showed slightly better performance.

**Figure 19.**
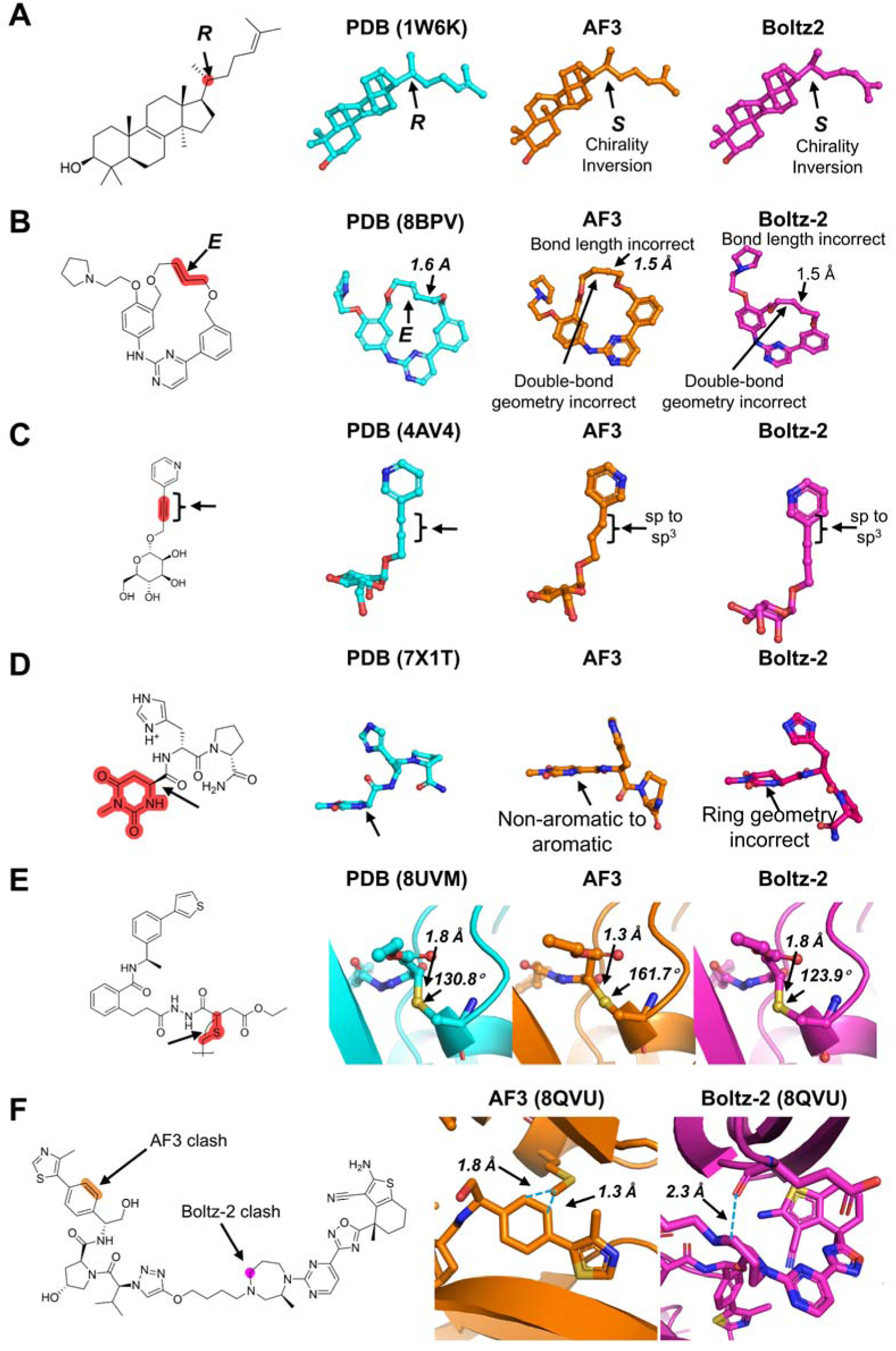
Representative local ligand chemical-structure and geometry errors in AF3 and Boltz-2 predictions. (**A**) Stereochemical inversion (PDB: 1W6K), where both AF3 and Boltz-2 predict the opposite configuration at the indicated chiral center relative to the experimental structure. (**B**) Alkene E/Z geometry and adjacent bond-length distortion (PDB: 8BPV). (**C**) Loss of linear sp-hybridized alkyne geometry (PDB: 4AV4), indicating a bond-order/hybridization-associated geometry error. (**D**) Ring-geometry distortion (PDB: 7X1T), including incorrect treatment of local aromatic/non-aromatic geometry and ring planarity. (**E**) Covalent bond-angle deviation in the SARS-CoV-2 PLpro covalent inhibitor complex (PDB: 8UVM). Together, these examples highlight local ligand chemical-structure errors that can occur despite reasonable global pose prediction, including stereochemical inversion, incorrect E/Z alkene geometry, bond-order-dependent distortion, ring-planarity errors, and covalent warhead-geometry deviations. (**F**) Representative steric clashes, where intermolecular distances are substantially shorter than typical noncovalent contact distances (PDB: 8QVU).

#### Perspective and Conclusion

Overall, our results suggest that AF3 and Boltz-2 are already useful for structure-based hypothesis generation, particularly when the binding site is well defined and ligand recognition is guided by conserved pharmacophores. The strongest performance was observed for canonical enzyme active-site ligands, many kinase inhibitors, selected covalent inhibitors, and several pharmacophore-defined PPI inhibitors. In these systems, the models frequently recovered key protein–ligand interactions and, in some kinase complexes, ligand-associated conformational states such as DFG and αC-helix organization. AF3 minPAE also provided useful pose triage: predictions with minPAE <0.85 Å were strongly enriched for accurate poses and pharmacophore recovery.

Performance was less consistent for allosteric modulators, cryptic-pocket binders, membrane-protein ligands, RNA binders, PROTACs, and molecular glues. These systems often require recovery of ligand-induced pocket formation, membrane context, RNA rearrangement, or ternary-interface organization. Although there were individual successes, predictions in these more complex systems were less reliable and generally required closer chemical inspection.

Beyond pose prediction, an important question is whether model-derived scores can support hit selection or affinity ranking. Across the evaluated datasets, pose-quality metrics and confidence scores such as minPAE did not show a consistent relationship with experimental activity. In contrast, the Boltz-2 affinity module captured relative activity trends in selected cases, including the Mpro dataset and the CRBN activity-cliff pair, although substantial compound-level variability remained and performance varied across ligand series. These results suggest that model-derived affinity estimates may be useful for compound prioritization and ranking, but are not sufficiently reliable for quantitative potency prediction.

A practical near-term workflow is therefore to use AF3 or Boltz-2 to generate ligand-bound structural hypotheses, followed by chemistry-aware evaluation and, when needed, physics-based refinement. Predictions can first be screened using confidence metrics and structural measures such as pharmacophore recovery and volume overlap, then inspected for stereochemistry, bond geometry, steric clashes, warhead orientation, subpocket occupancy, and consistency with SAR. Higher-priority poses can subsequently be evaluated using molecular dynamics, MM-GBSA, FEP, or related methods. This approach takes advantage of the ability of AF3 and Boltz-2 to generate protein conformations and ligand poses that may be difficult to obtain from rigid docking, while physics-based and chemistry-aware evaluation can help with compound prioritization.

We observed a clear trend in which prediction accuracy decreased with increasing ligand conformational flexibility, as seen in our sEH dataset. This limitation is likely multifactorial: additional rotatable bonds expand the conformational search space, input conformers generated by ETKDGv3-type methods may not represent the protein-bound state for more flexible ligands, and the models may not reliably rank multiple chemically plausible poses. In several cases, key pharmacophore interactions were preserved, yet flexible ligand arms were assigned to the wrong subpockets, suggesting that the primary challenge was global pose assignment rather than local interaction recognition. These findings do not preclude practical use in structure-guided design, but suggest that predictions for highly flexible ligands should be treated as candidate binding hypotheses and evaluated together with experimental SAR or additional computational analysis. Overall, AF3 and Boltz-2 are most useful as tools for generating and prioritizing structural hypotheses, rather than as stand-alone predictors of binding pose or activity.

## EXPERIMENTAL SECTION

### AF3 and Boltz-2 Complex Prediction

Complex predictions were generated using locally installed versions of the official AlphaFold 3 (v3.0.1) and Boltz-2 (v2.2.0) GitHub implementations and their released pretrained model weights, following the standard workflows described in the respective repositories. No modifications were made to the model architecture, pretrained weights, or inference code. Unless otherwise stated, default data-processing and inference settings were used. For AF3, random seeds 10 and 42 were used, with five structural predictions generated per seed, yielding 10 predictions per complex. For Boltz-2, 10 structural predictions were generated directly for each complex using the standard inference workflow. Predictions were performed using NVIDIA GeForce RTX 4090 GPUs with 24 GB of GPU memory, NVIDIA RTX PRO 6000 GPUs with 96 GB of GPU memory. The curated dataset with PDB details are summarized in **Table S1**. Protein, peptide, and RNA components were supplied as separate polymer chains, and small-molecule ligands were specified using SMILES strings, CCD identifiers, or user-defined CCD components, as appropriate. Where applicable, covalent linkages were explicitly defined using bonded atom pairs in AF3 and bond constraints in Boltz-2. The Boltz-2 affinity-prediction module was enabled for affinity analyses of the sEH inhibitors and Mpro inhibitors. Detailed analysis protocols and scripts are provided in the accompanying GitHub repository https://github.com/RX-Medchem/docking-evaluation.

### Structural Alignment and Small-Molecule Extraction

Predicted AF3 and Boltz-2 complexes were aligned to the corresponding experimental structures using PyMOL (Schrödinger LLC).^86^ Predicted structures were superimposed onto the corresponding experimental structures using PyMOL’s align algorithm, which employs a custom BLOSUM62-weighted, BLAST-like dynamic-programming sequence-alignment algorithm followed by structural superposition and iterative outlier rejection. The alignment was restricted to the corresponding polymer components, with ligand and solvent atoms excluded. For multichain complexes, corresponding chains were first matched using pairwise Cα-based alignments and then used for final structural superposition. The resulting transformation was applied to the entire predicted complex, so that ligand coordinates were transformed together with the protein without additional ligand fitting. Reference ligands were extracted from the experimental structures after removal of polymer atoms, solvent molecules, and common nucleotide cofactors (ATP, ADP, GTP, and GDP). The aligned predicted ligands were then compared with the experimental binding poses using overall ligand RMSD, pharmacophore RMSD, and volume overlap. Per-atom positional deviations were also calculated and mapped onto two-dimensional ligand structures to visualize local differences between predicted and experimental poses.

### Ligand Pose RMSD, Pharmacophore RMSD, and Volume Overlap Analysis

Ligand pose differences were quantified using RDKit-based heavy-atom mapping and RMSD analysis. AF3 output mmCIF files do not preserve the original PDB chain IDs or residue numbering. Atom mapping was performed using RDKit’s GetSubstructMatch() function based on atom types and connectivity after removal of hydrogen. RMSD was calculated from the distances between corresponding heavy atoms after protein alignment.

Pharmacophore-specific RMSD was calculated using ligand atoms preferentially selected based on key recognition interactions identified from the experimental complex and relevant structural literature. RMSD was restricted to these predefined pharmacophore atoms to evaluate the accuracy of functionally important binding features.

Volume overlap was defined as the fraction of the experimental reference-ligand volume occupied by the predicted ligand, *V_intersection_*/*V_ref_* . Because the RDKit shape-comparison workflow does not directly provide the intersecting volume, *V_intersection_* was derived from the shape Tanimoto similarity. For shape comparison, RDKit represents each heavy atom using its element-specific van der Waals radius (e.g., 1.70 Å for carbon; vdwScale = 1.0) and encodes the resulting molecular shape on a three-dimensional grid. Molecular volumes were calculated with hydrogen atoms excluded. Tanimoto similarity was calculated as T= *V_intersection_*/(*V_ref_* + *V_pred_* - *V_intersection_*), corresponding to the ratio of the intersecting volume to the union volume. The intersecting volume was then obtained from this relationship as *V_intersection_* = *T*(*V_ref_* + *V_pred_*)/(1+ T) . The final volume-overlap metric was calculated as *V_intersection_*/*V_ref_* = *T*(*V_ref_* + *V_pred_*)/[(1+ *T*)*V_ref_*].

### Two-Dimensional Visualization of Ligand Structural Differences

Two-dimensional per-atom deviation heatmaps were generated using RDKit to visualize local differences in ligand poses. For each ligand atom, the positional deviation from the experimental reference was mapped onto a fixed 0–10 Å color scale using the reversed RdYlGn colormap. In addition to individual predicted poses, a global-average heatmap was generated by averaging the positional deviation of each atom across all predicted models.

### Statistical analysis

Pearson correlation analysis was used to assess linear relationships between AI-derived model metrics and reference measurements, while Spearman rank correlation analysis was used to evaluate monotonic associations without assuming linearity. Specifically, correlations were evaluated between model-derived confidence or prediction metrics and reference measurements, including ligand RMSD, pharmacophore RMSD, volume overlap, and reference binding affinity values, where applicable. All correlation analyses were performed using GraphPad Prism. Two-tailed *P* values were reported, with *P* < 0.05 considered statistically significant. The residual standard deviation from simple linear regression was also reported where applicable.

### Supporting Information

Boltz-2 prediction violin plots; summary of PDB entries, names, index numbers, and annotations; detailed tables of AF3 and Boltz-2 entries and prediction statistics; coordinate files for computational models.

### PDB ID Codes

Previously deposited structures used in this study: 8BM2 (7); 8BPW (8); 8BX6 (9); 8BX9 (10); 8BXC (11); 8BXH (12); 8S9A (13); 8C7Y (14); 8FV3 (15); 8FV4 (16); 8U8J (17); 8ZQW (18); 9BYJ (19); 9CJ1 (20); 9CJ4 (21); 8HUK (22); 8HUQ (23); 8UOB (24); 9CSY (25); 8PFO (26); 7GQU (27); 8A2D (28); 8JUE (29); 8AZV (30); 8S0O (31); 8C14 (32); 9X53 (33); 9MJM (34); 8XGV (35); 8T5G (36); 8T5M (37); 8T5R (38); 8UC9 (39); 8UH0 (40); 9D0X (41); 8QVU (42); 9RK8 (43); 9L6F (44); 9BG0 (45); 8G9P (46); 8OV6 (47); 9DUR (48); 8TZX (49); 8UVM (50); 8V3A (51); 8V39 (52); 8V4U (53); 8JHL (54); 8ZKN (55); 9CY3 (56); 9LK8 (57); 8R62 (58); 8QMH (59); 8R8P (60); 8ZNQ (61); 9CPD (62); 9IO0 (63); 7GRE (64); 7GRF (65); 7GRJ (66); 7GRN (67); 7GRS (68); 7GRT (69); 7GRU (70); 7GRZ (71); 7GS0 (72); 9BVE (73); and 4CI1 (74 and 75); 7GLV (76); 7GAW (77); 7GKS (78); 7GNT (79); 7GLG (80); 7GIU (81); 7GM1 (82); 7GNA (83); 7GIX (84); 7GHM (85); 7GG2 (86); 7GMR (87); 7GE3 (88); 7GMP (89); 7GE2 (90); 7GE4 (91); 7GGC (92); 7GFJ (93); 7GCK (94); and 7GBV (95). PDB accession codes for the six newly determined sEH complexes (1–6) will be provided upon deposition. Authors will release the atomic coordinates and experimental data upon article publication.

## AUTHOR INFORMATION

### Author contributions

K.C. and Z.Q. contributed equally. R.X. conceived the project and curated the dataset. K.C., Z.Q., H.L., A.L., and R.X. coded the script for data analysis. K.C. and M.M. performed the sEH assays and data analysis. R.X., K.C., and Z.Q. wrote the manuscript. M.R.G. All authors discussed the results and contributed to the final editing of the manuscript.

## Acknowledgements

This study is supported in part by NIH R01AI168165 to RX, by the University of Arizona College of Pharmacy faculty startup fund to R.X. and H.L., and by R. Ken and Donna Coit Endowed Chair fund in Drug Discovery to H.L.

## ABBREVIATIONS

AF3: AlphaFold 3;
Å: angstrom
ATP: adenosine triphosphate
Aurora A: Aurora kinase A
BL2: blocking loop 2
BRAF: B-Raf proto-oncogene, serine/threonine kinase
BRD4: bromodomain-containing protein 4
C9orf72: chromosome 9 open reading frame 72
CDK2: cyclin-dependent kinase 2
CRBN: cereblon
cryo-EM: cryogenic electron microscopy
CYPA: cyclophilin A
DCAF16: DDB1- and CUL4-associated factor 16
DDB1: DNA damage-binding protein 1
E3: ubiquitin-protein ligase E3
EGFR: Epidermal growth factor receptor
ENL: eleven-nineteen leukemia
ERK2: Extracellular signal-regulated kinase 2
ETKDGv3: experimental-torsion knowledge distance geometry version 3
FEP: free energy perturbation
GABA: γ-aminobutyric acid / gamma-aminobutyric acid
GAT3: GABA transporter 3 / sodium- and chloride-dependent GABA transporter 3
GDP: guanosine diphosphate
Hck: Hematopoietic cell kinase
ipTM: interface predicted Template Modeling score
JAK2: Janus kinase 2
KEAP1: Kelch-like ECH-associated protein 1
KRAS: KRAS proto-oncogene, GTPase
MCT8: monocarboxylate transporter 8
MD: molecular dynamics
MM-GBSA: molecular mechanics-generalized Born surface area
Mpro: main protease
NRAS: NRAS proto-oncogene, GTPase
NRF2: nuclear factor erythroid 2-related factor 2
OATP1B1: organic anion transporting polypeptide 1B1
p38α: p38 alpha mitogen-activated protein kinase
PAE: predicted aligned error
PLpro: Papain-like protease
PPAR: Peroxisome proliferator-activated receptor
PPI: protein–protein interaction
PROTAC: Proteolysis-targeting chimera
Rac1 / RAC1: Ras-related C3 botulinum toxin substrate 1
RhoGDI2: Rho GDP-dissociation inhibitor 2
RMSD: Root-Mean-Square Deviation
SAR: Structure–Activity Relationship
SARS-CoV-2: severe acute respiratory syndrome coronavirus 2
sEH: soluble epoxide hydrolase
SHP2: Src homology region 2 domain-containing phosphatase 2
SMN2: survival motor neuron 2
SOS1: son of sevenless homolog 1
SOS2: son of sevenless homolog 2
TPX2: targeting protein for Xklp2
TYK2 JH2: Tyrosine kinase 2 Janus homology 2 domain
VHL: von Hippel–Lindau protein
WIZ: widely interspaced zinc finger protein
WRN: Werner syndrome ATP-dependent helicase
WT: wild type.

