## Supplementary material for "A Medicinal Chemistry-Centered Evaluation of AlphaFold 3 and Boltz-2 Across Diverse Binding Modalities": 4. SI.docx

**Table of Contents**

1. Violin plot of Boltz-2 predictions….......................................................................pp 2-10

2. Table of PDB Entries, Names, Index Numbers, and Annotations ……..……….pp 11-14

3. Tables of AF3 and Boltz-2 Entries and Prediction Statistics ……….…………. pp 15-39

4. References…....................................................................................................pp 40-43

**Violin plot of Boltz-2 predictions**

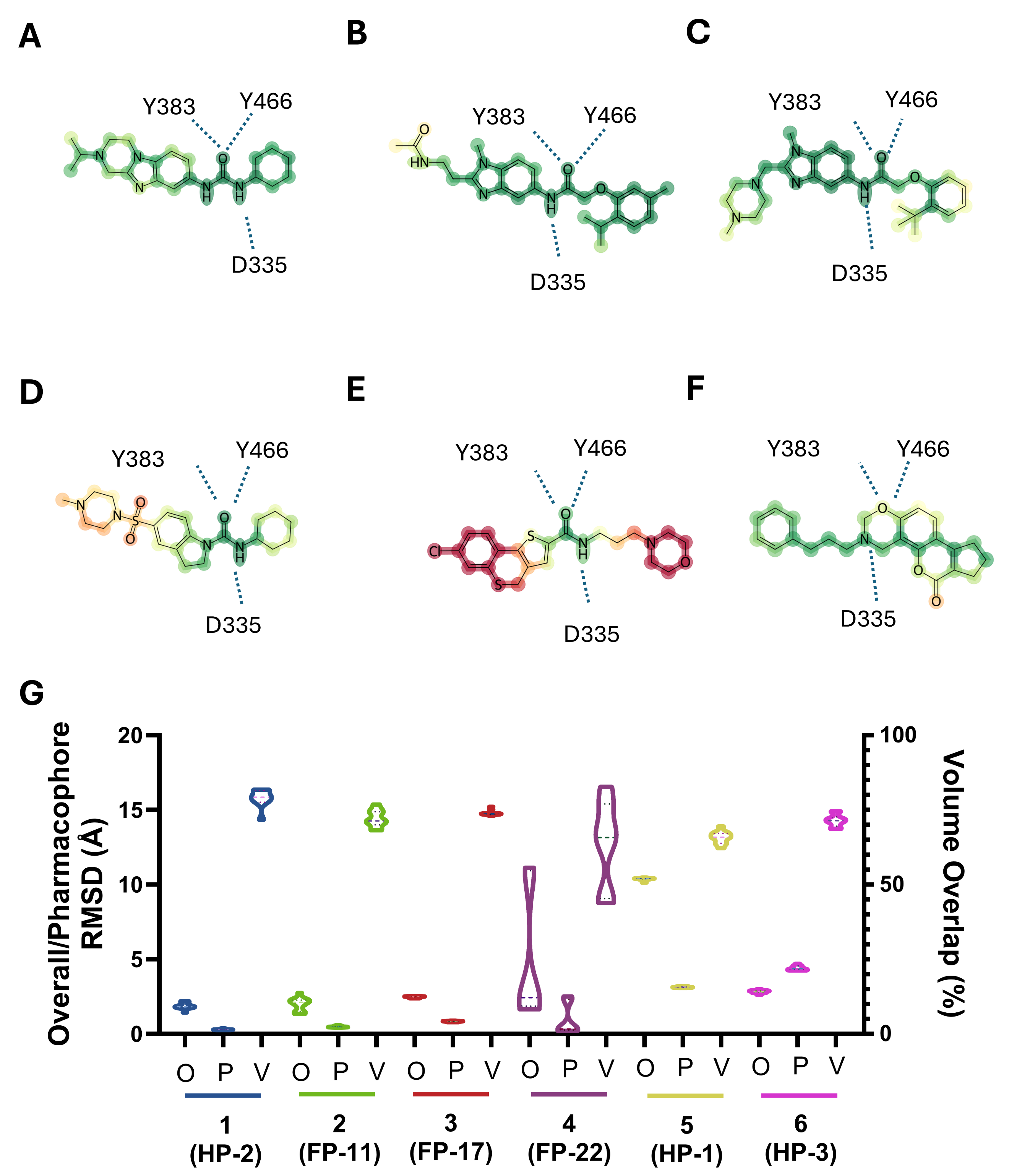

**Figure S1**. (**A-F**) Boltz-2 prediction analysis of six sEH inhibitor complexes (sEH-1–sEH-6). 2D ligand interaction maps colored by per-atom overall prediction RMSD, ranging from green (highly accurate atom placement) to red (poorly predicted atom placement). (**G**) Violin plot of Boltz-2 predictions. Quantitative evaluation of six Boltz-2 predictions using ligand overall RMSD (O), ligand pharmacophore RMSD (P), and ligand volume overlap (%) (V).

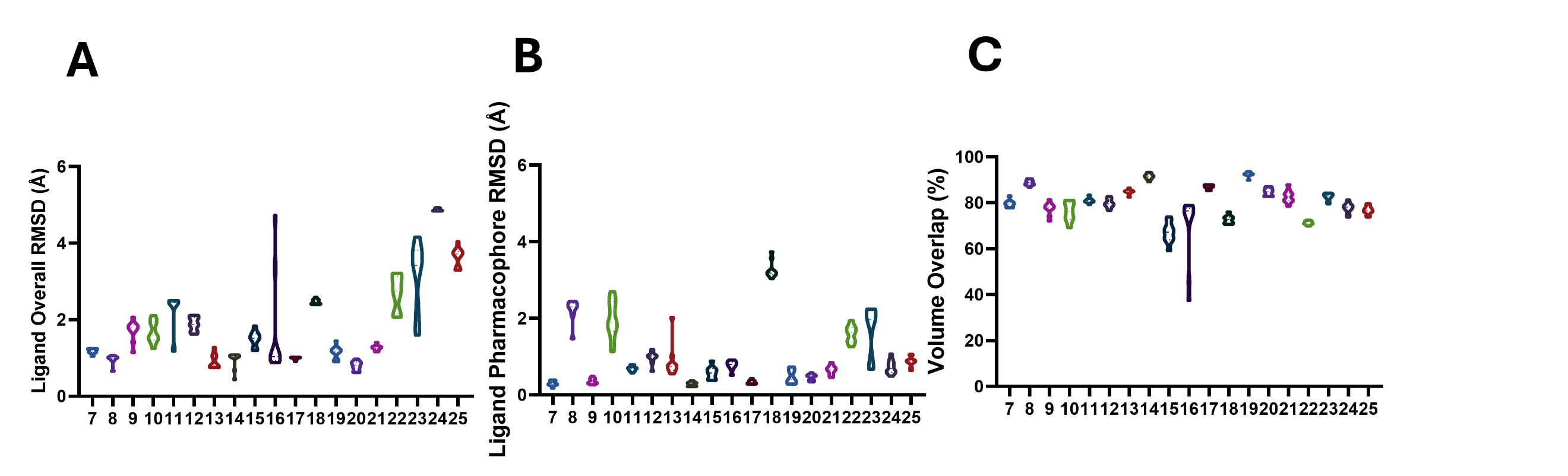

**Figure S2**. (**A-C**) Boltz-2 prediction analysis of canonical ligand-binding complexes. Dataset-wide evaluation of ligand overall RMSD, pharmacophore RMSD, volume overlap (%). For entries 7–25, the corresponding PDB structures are 8BM2, 8BPW, 8BX6, 8BX9, 8BXC, 8BXH, 8S9A, 8C7Y, 8FV3, 8FV4, 8U8J, 8ZQW, 9BYJ, 9CJ1, 9CJ4, 8HUK, 8HUQ, 8UOB, and 9CSY.

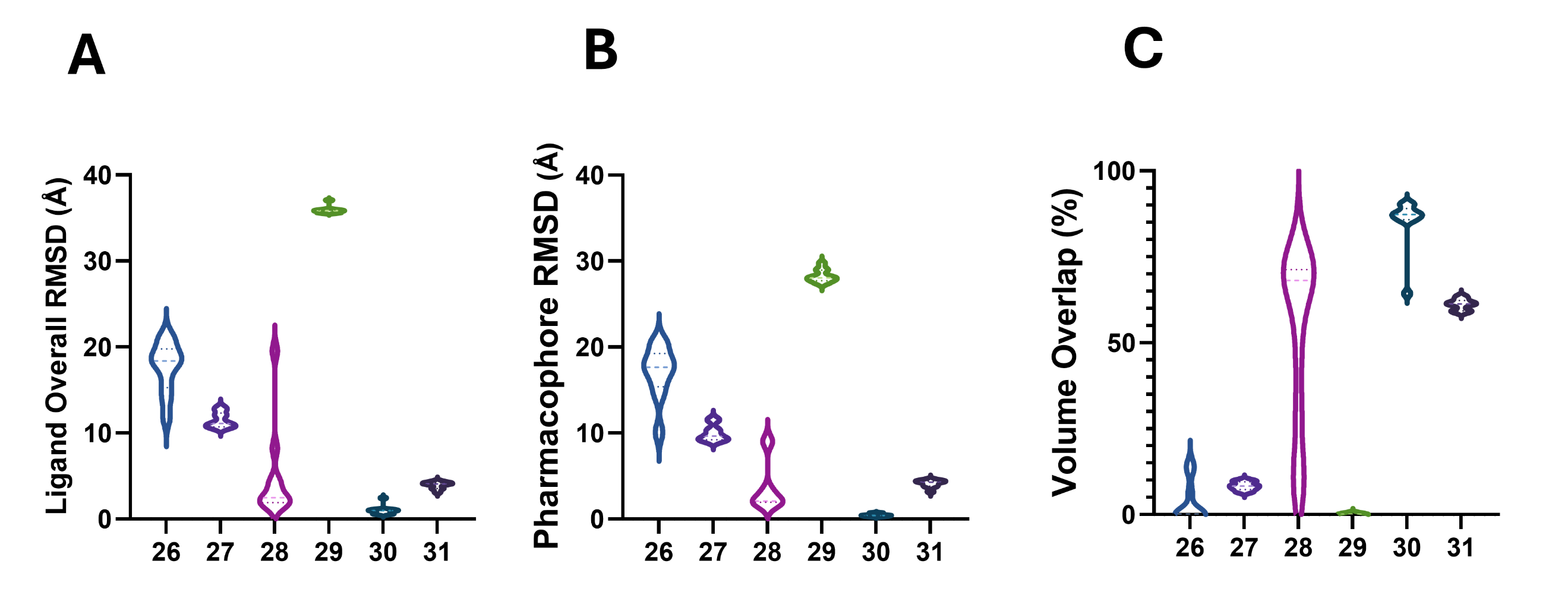

**Figure S3**. (**A-C**) Boltz-2 prediction analysis of allosteric inhibitors. Dataset-wide evaluation of ligand overall RMSD, pharmacophore RMSD, volume overlap (%). For entries 26-31, the corresponding PDB structures are 8PFO, 7GQU, 8A2D, 8JUE, 8AZV, 8S0O.

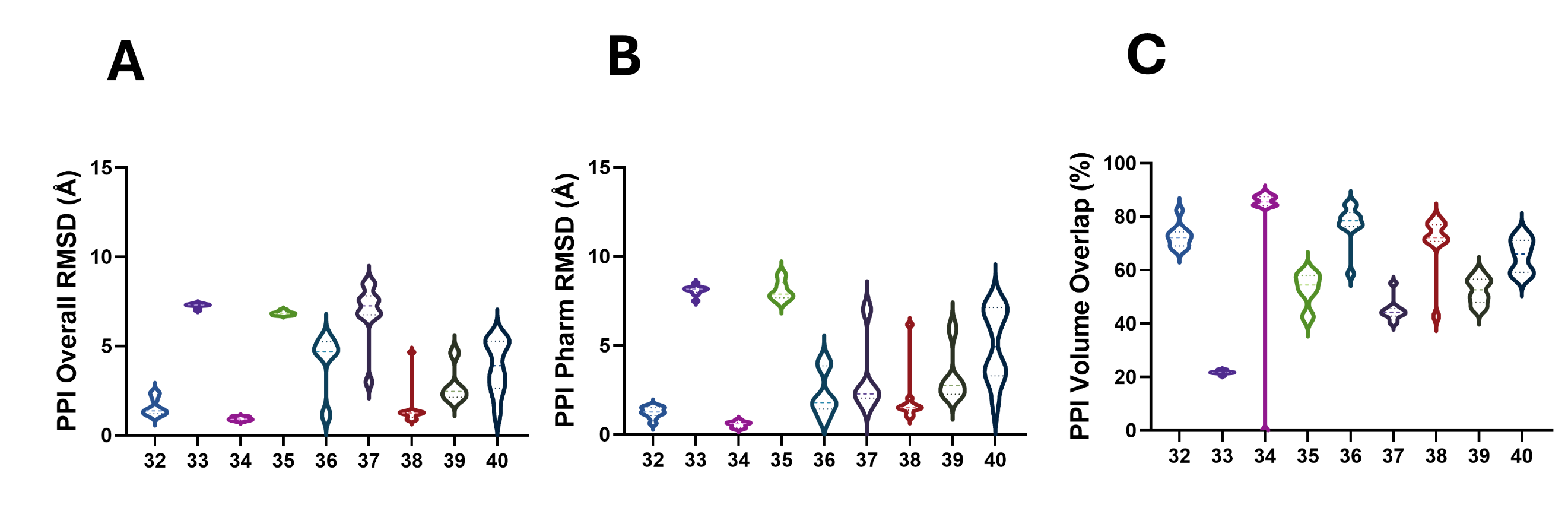

**Figure S4**. (**A-C**) Boltz-2 prediction analysis of PPI inhibitors. Dataset-wide evaluation of ligand overall RMSD, pharmacophore RMSD, volume overlap (%). For entries 32-40, the corresponding PDB structures are 8C14, 9X53, 9MJM, 8XGV, 8T5G, 8T5M, 8T5R, 8UC9 and 8UH0.

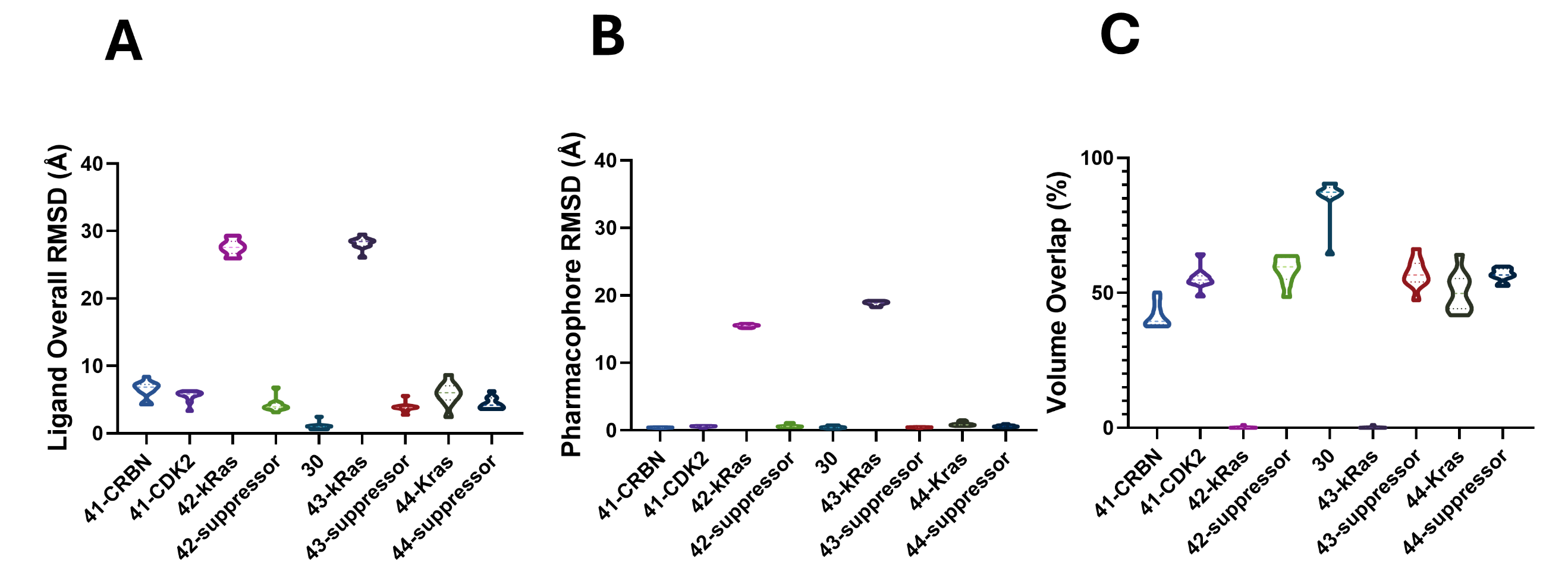

**Figure S5**. (**A-C**) Boltz-2 prediction analysis of heterobifunctional PROTAC complexes. Dataset-wide evaluation of ligand overall RMSD, pharmacophore RMSD, volume overlap (%). For PROTAC entries, the corresponding PDB structures are 9D0X (41), 8QVU (42), 8AZV (30), 9RK8 (43), 9L6F (44).

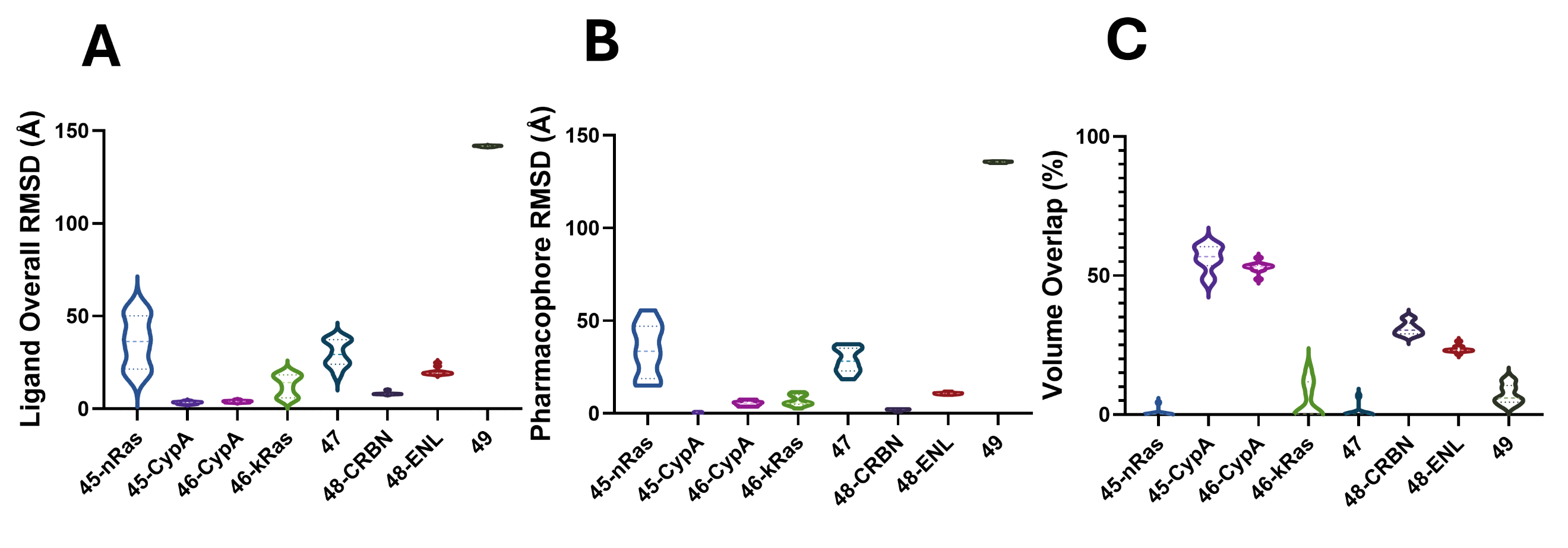

**Figure S6**. (**A-C**) Boltz-2 prediction analysis of molecular glue. Dataset-wide evaluation of ligand overall RMSD, pharmacophore RMSD, volume overlap (%). For molecular glue entries, the corresponding PDB structures are 9BG0 (45), 8G9P (46), 8OV6 (47), 9DUR (48), 8TZX (49).

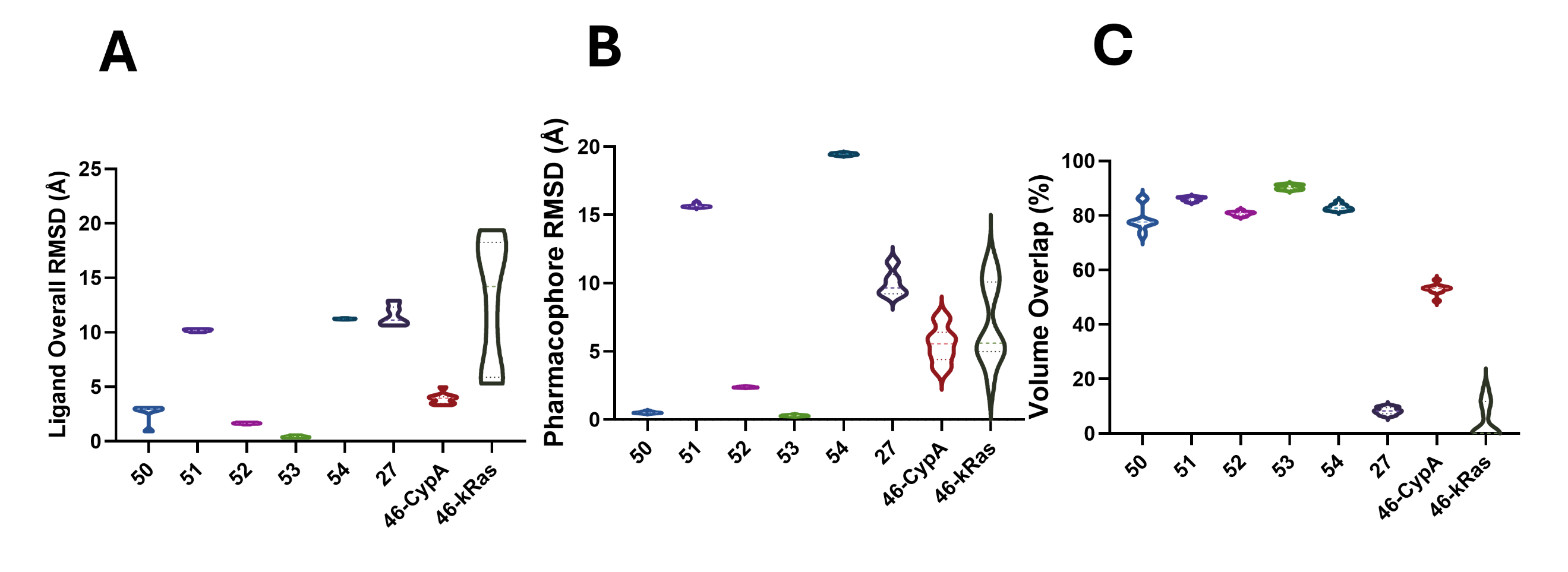

**Figure S7**. (**A-C**) Boltz-2 prediction analysis of covalent inhibitors. Dataset-wide evaluation of ligand overall RMSD, pharmacophore RMSD, volume overlap (%). For covalent inhibitors entries, the corresponding PDB structures are 8UVM (50), 8V3A (51), 8V39 (52), 8V4U (53), 8JHL (54), 7GQU (27), and 8G9P (46).

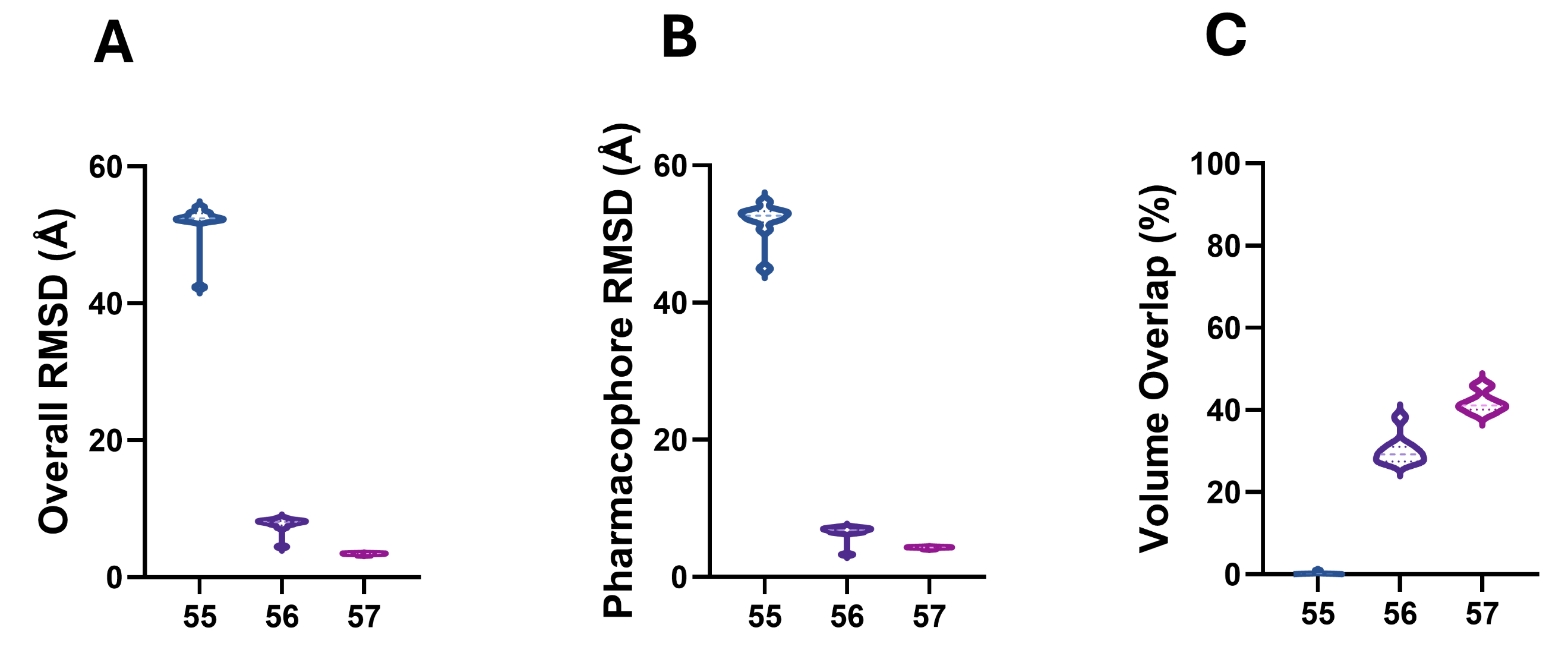

**Figure S8**. (**A-C**) Boltz-2 prediction analysis of membrane. Dataset-wide evaluation of ligand overall RMSD, pharmacophore RMSD, volume overlap (%). For entries 55-57, the corresponding PDB structures are 8ZKN, 9CY3, 9LK8.

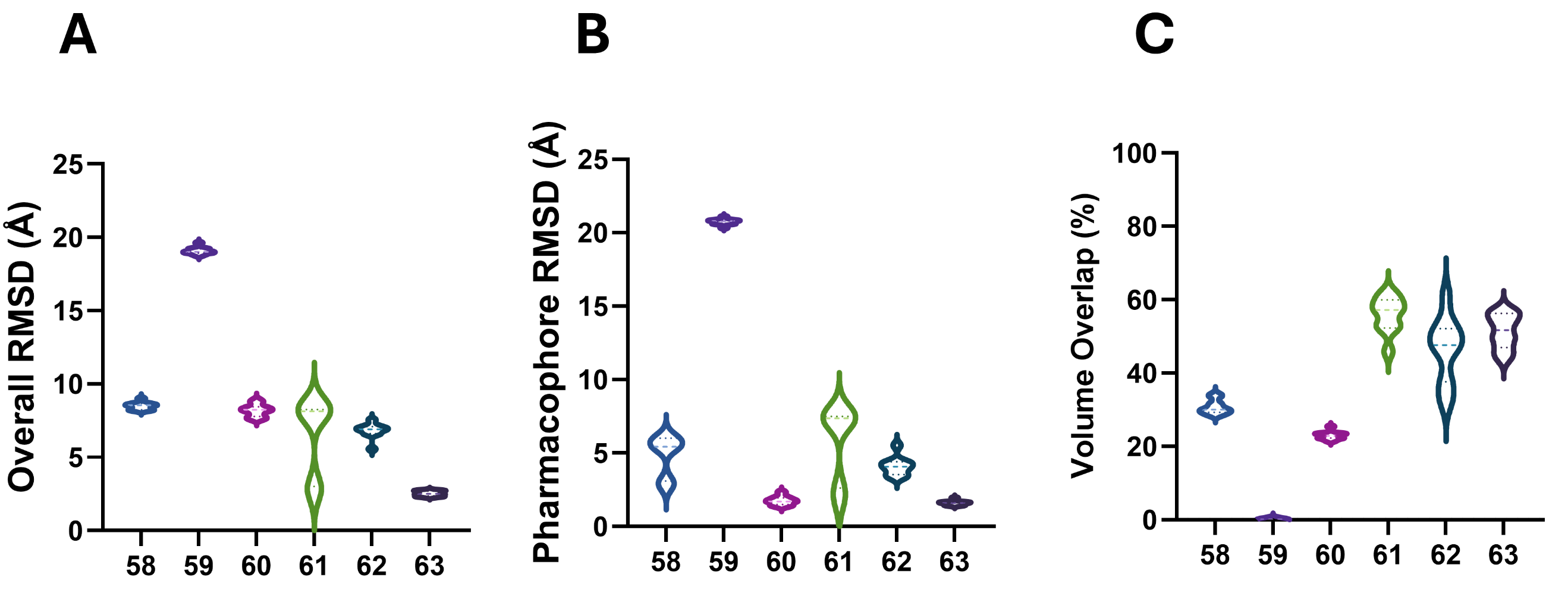

**Figure S9**. (**A-C**) Boltz-2 prediction analysis of RNA. Dataset-wide evaluation of ligand overall RMSD, pharmacophore RMSD, volume overlap (%). For entries 58-63, the corresponding PDB structures are 8R62, 8QMH, 8R8P, 8ZNQ, 9CPD and 9IO0.

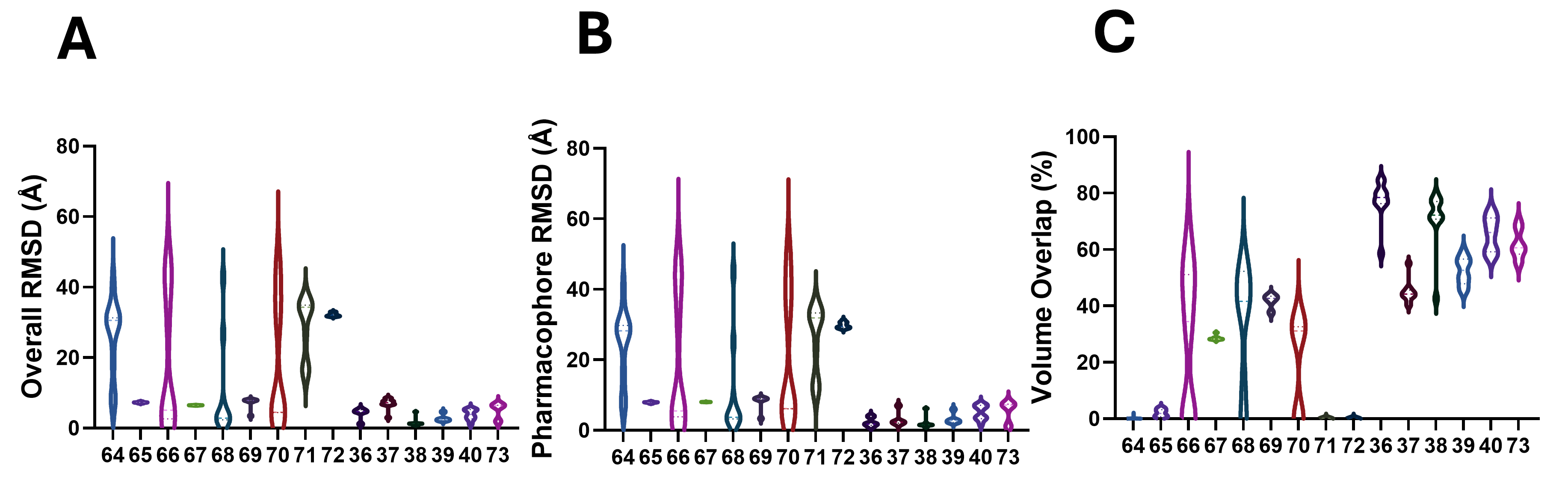

**Figure S10**. (**A-C**) Boltz-2 prediction analysis of fragments. Dataset-wide evaluation of ligand overall RMSD, pharmacophore RMSD, volume overlap (%). For entries 36-40, 64-73, the corresponding structures are 8T5G, 8T5M, 8T5R, 8UC9, 8UH0, 7GRE, 7GRF, 7GRJ, 7GRN, 7GRS, 7GRT, 7GRU, 7GRZ, 7GS0 and 9BVE.

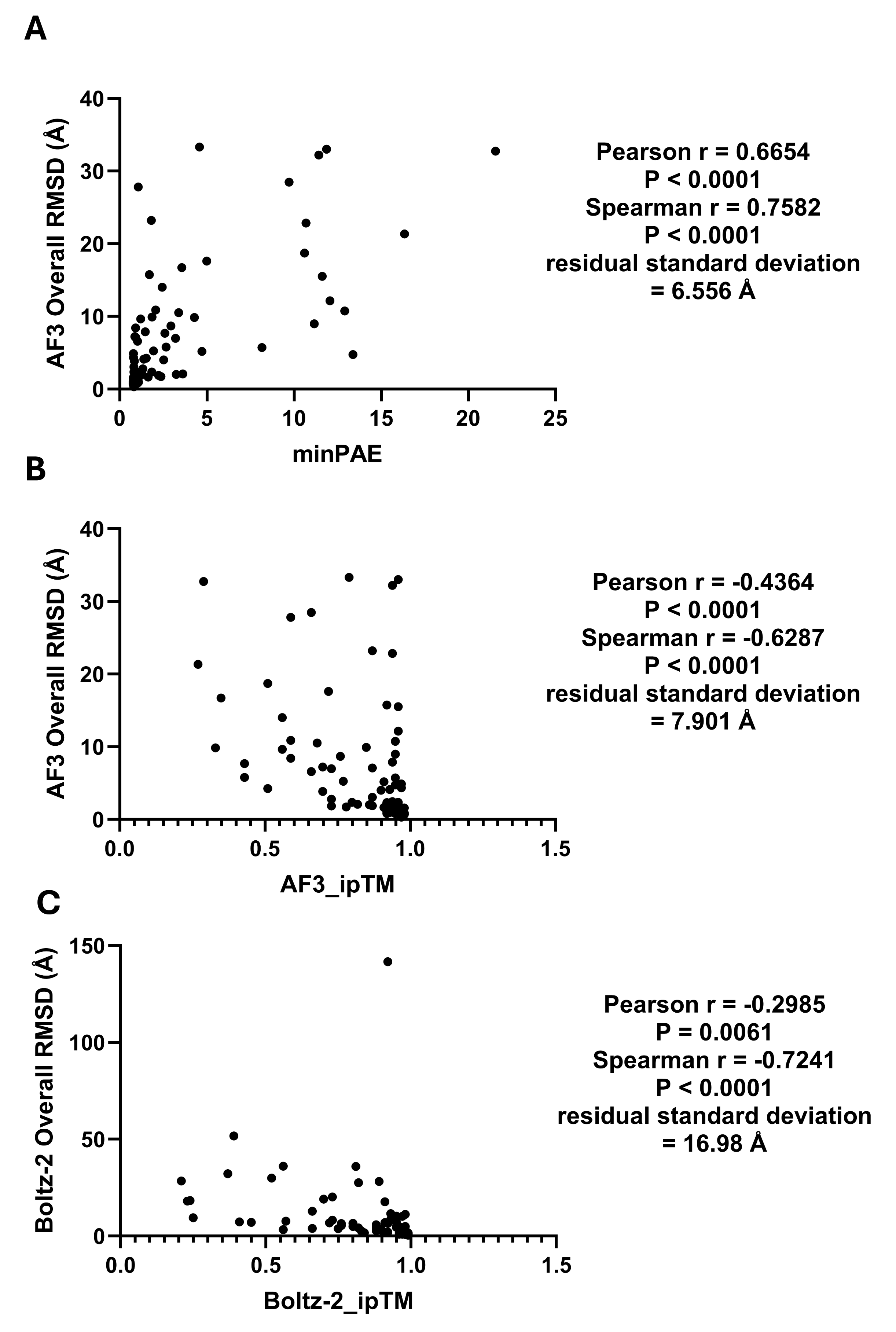

**Figure S11**. Relationship between predicted pose accuracy and model confidence for all complexes. (**A**) AF3 overall ligand RMSD versus minPAE. (**B**) AF3 overall ligand RMSD versus ipTM. (**C**) Boltz-2 overall ligand RMSD versus ipTM. Lower RMSD indicates greater agreement with the experimental ligand pose.

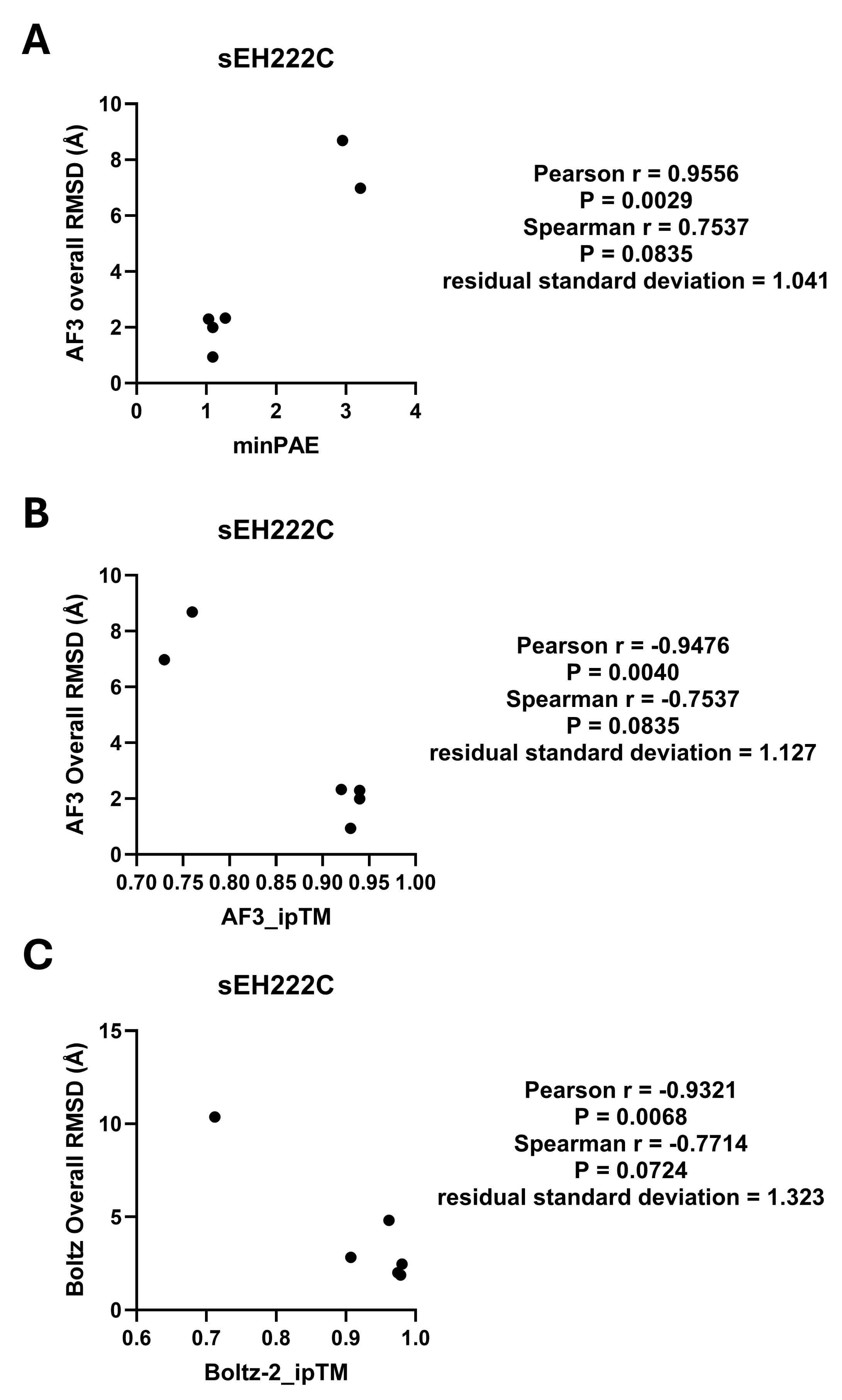

**Figure S12**. Relationship between predicted pose accuracy and model confidence for sEH222C. (**A**) AF3 overall ligand RMSD versus minPAE. (**B**) AF3 overall ligand RMSD versus ipTM. (**C**) Boltz-2 overall ligand RMSD versus ipTM. Lower RMSD indicates greater agreement with the experimental ligand pose.

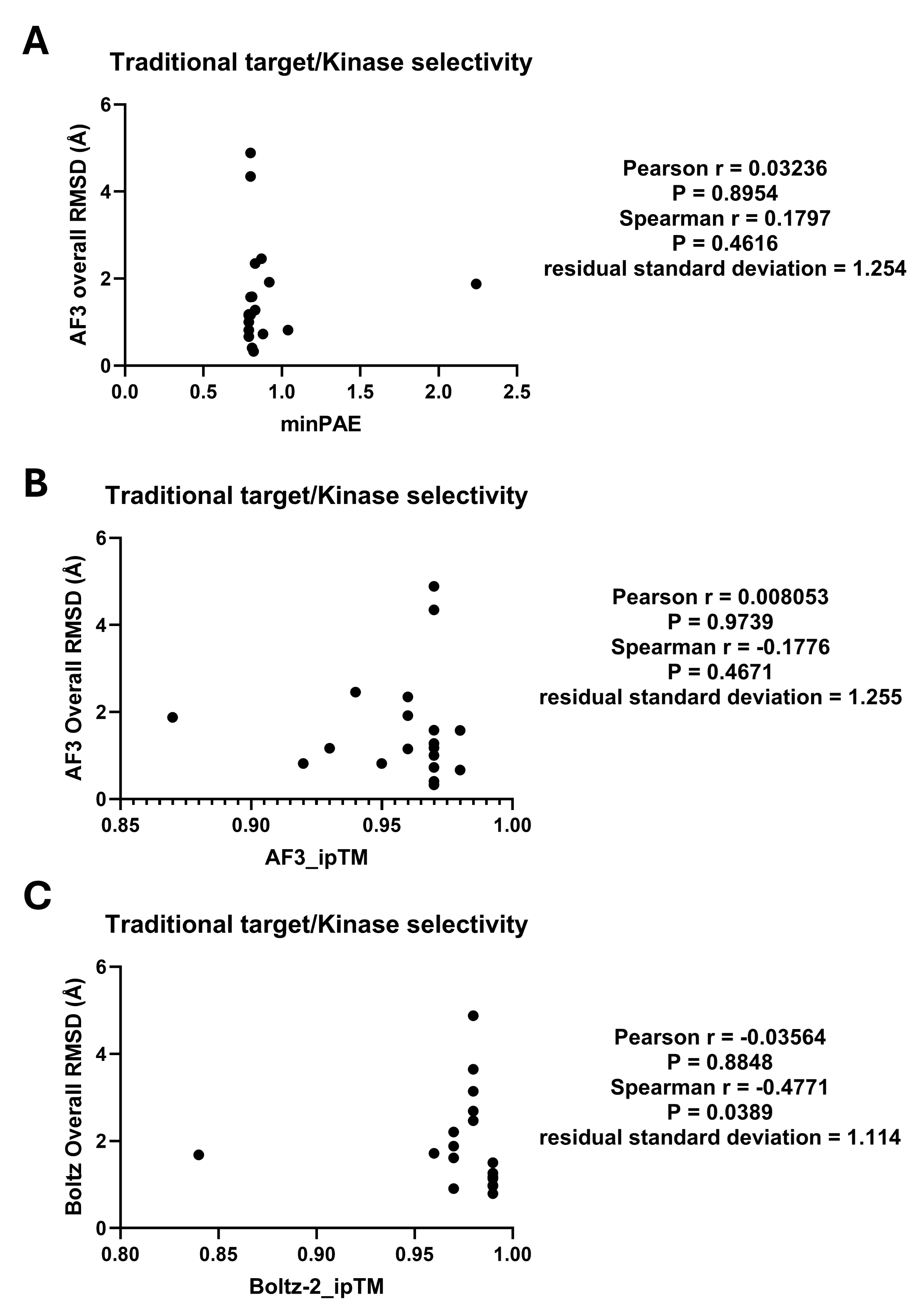

**Figure S13**. Relationship between predicted pose accuracy and model confidence for canonical ligand-binding complexes. (**A**) AF3 overall ligand RMSD versus minPAE. (**B**) AF3 overall ligand RMSD versus ipTM. (**C**) Boltz-2 overall ligand RMSD versus ipTM. Lower RMSD indicates greater agreement with the experimental ligand pose.

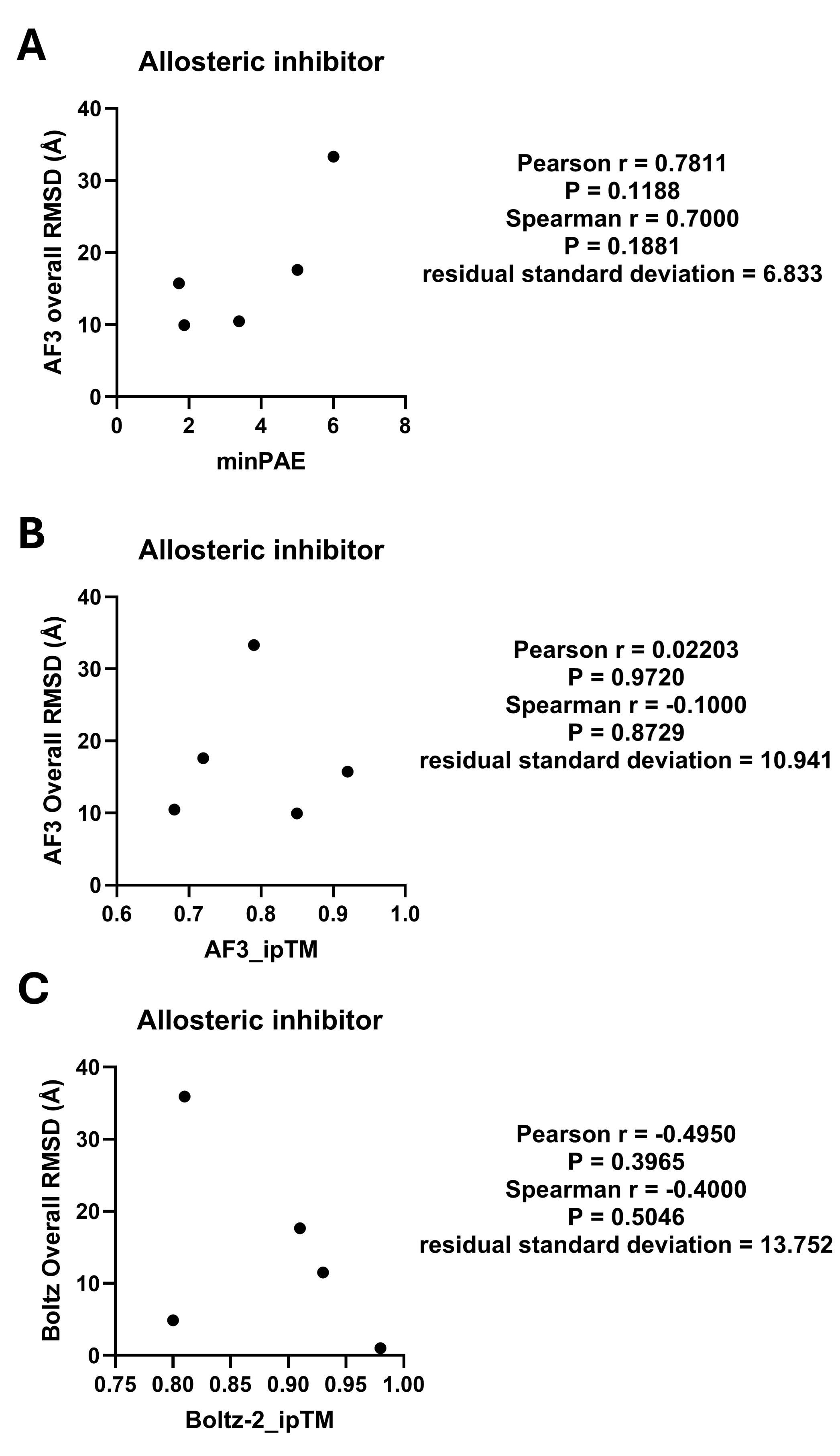

**Figure S14**. Relationship between predicted pose accuracy and model confidence for allosteric inhibitors. (**A**) AF3 overall ligand RMSD versus minPAE. (**B**) AF3 overall ligand RMSD versus ipTM. (**C**) Boltz-2 overall ligand RMSD versus ipTM. Lower RMSD indicates greater agreement with the experimental ligand pose.

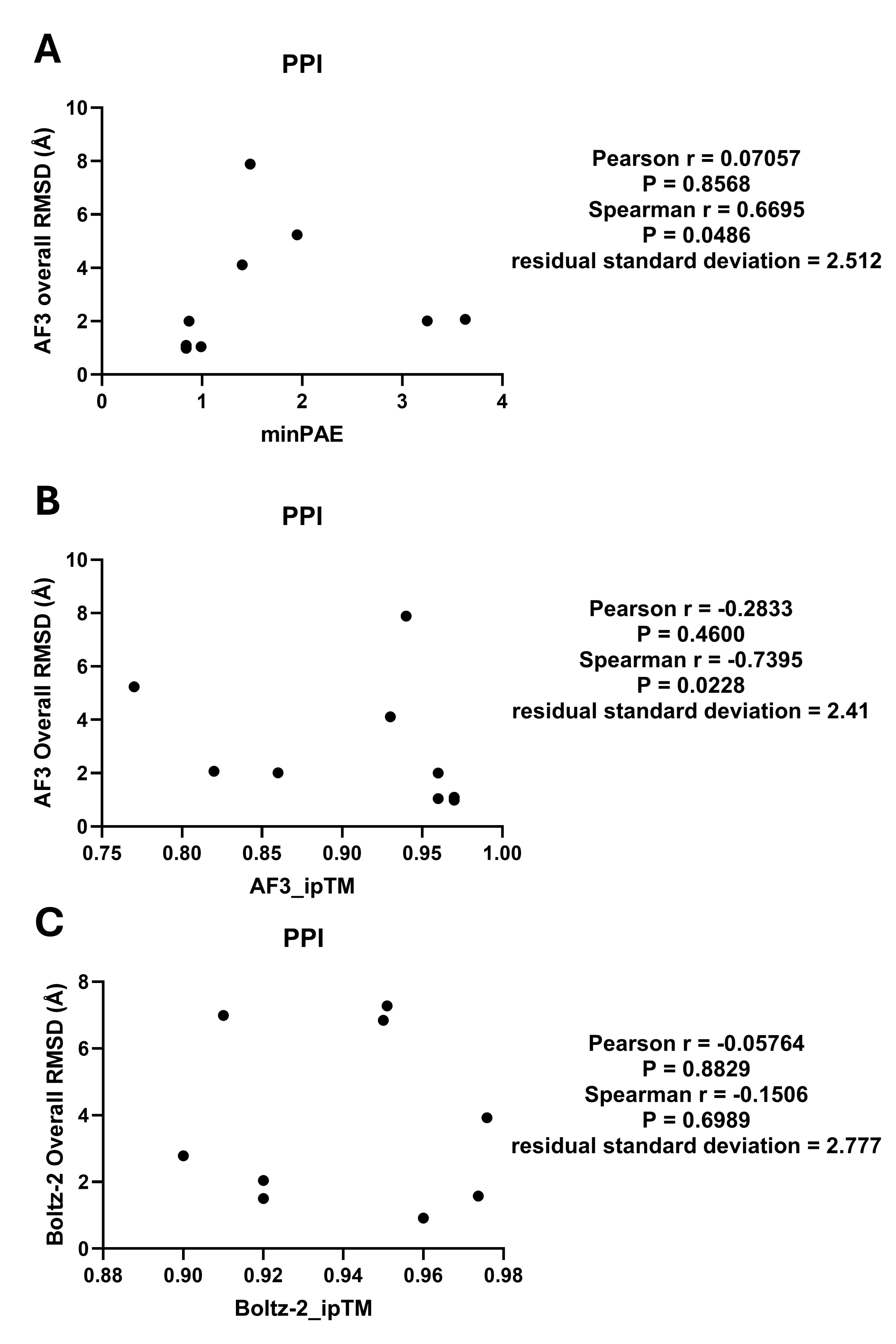

**Figure S15**. Relationship between predicted pose accuracy and model confidence for PPI. (**A**) AF3 overall ligand RMSD versus minPAE. (**B**) AF3 overall ligand RMSD versus ipTM. (**C**) Boltz-2 overall ligand RMSD versus ipTM. Lower RMSD indicates greater agreement with the experimental ligand pose.

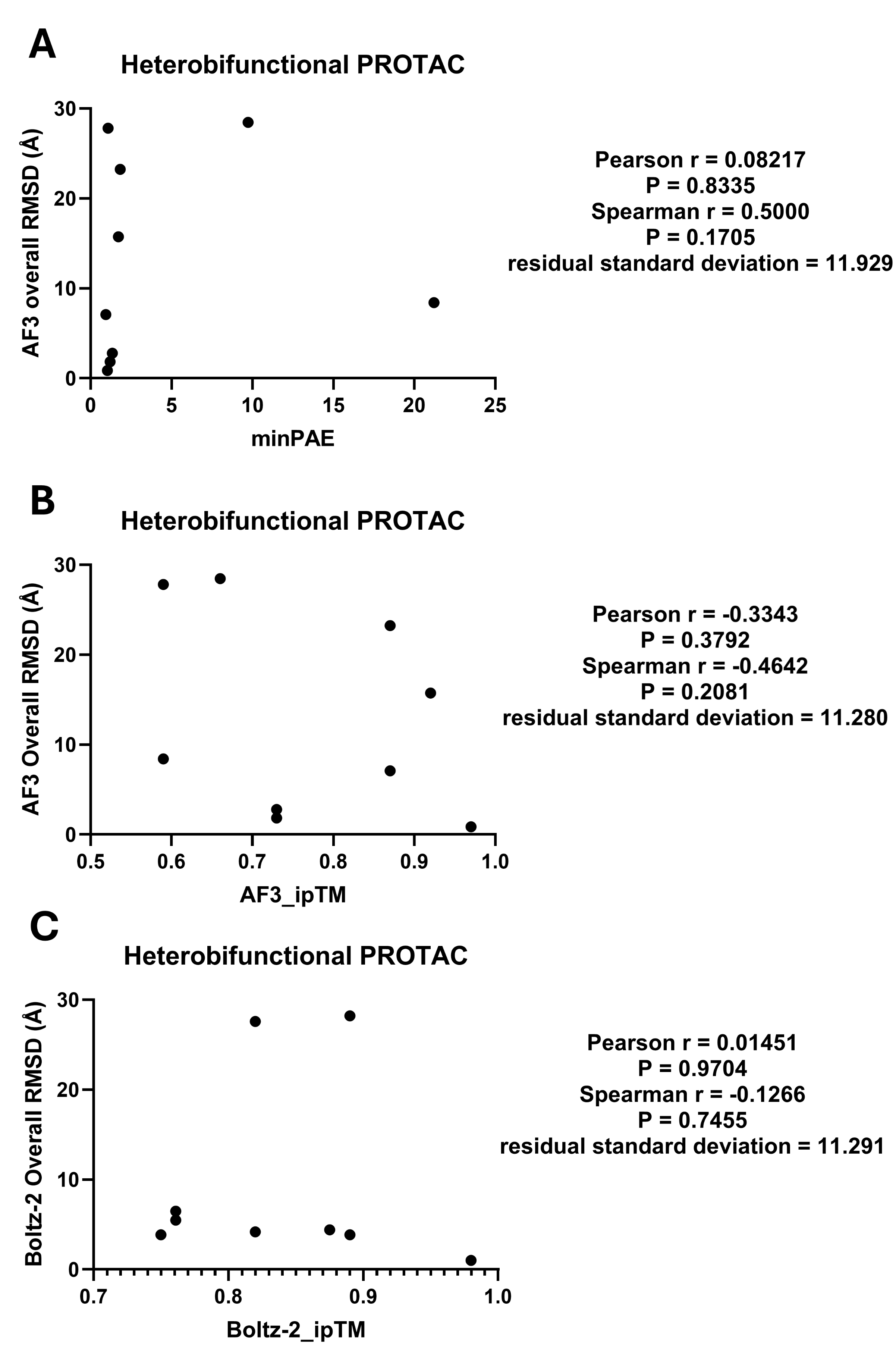

**Figure S16**. Relationship between predicted pose accuracy and model confidence for heterobifunctional PROTAC. (**A**) AF3 overall ligand RMSD versus minPAE. (**B**) AF3 overall ligand RMSD versus ipTM. (**C**) Boltz-2 overall ligand RMSD versus ipTM. Lower RMSD indicates greater agreement with the experimental ligand pose.

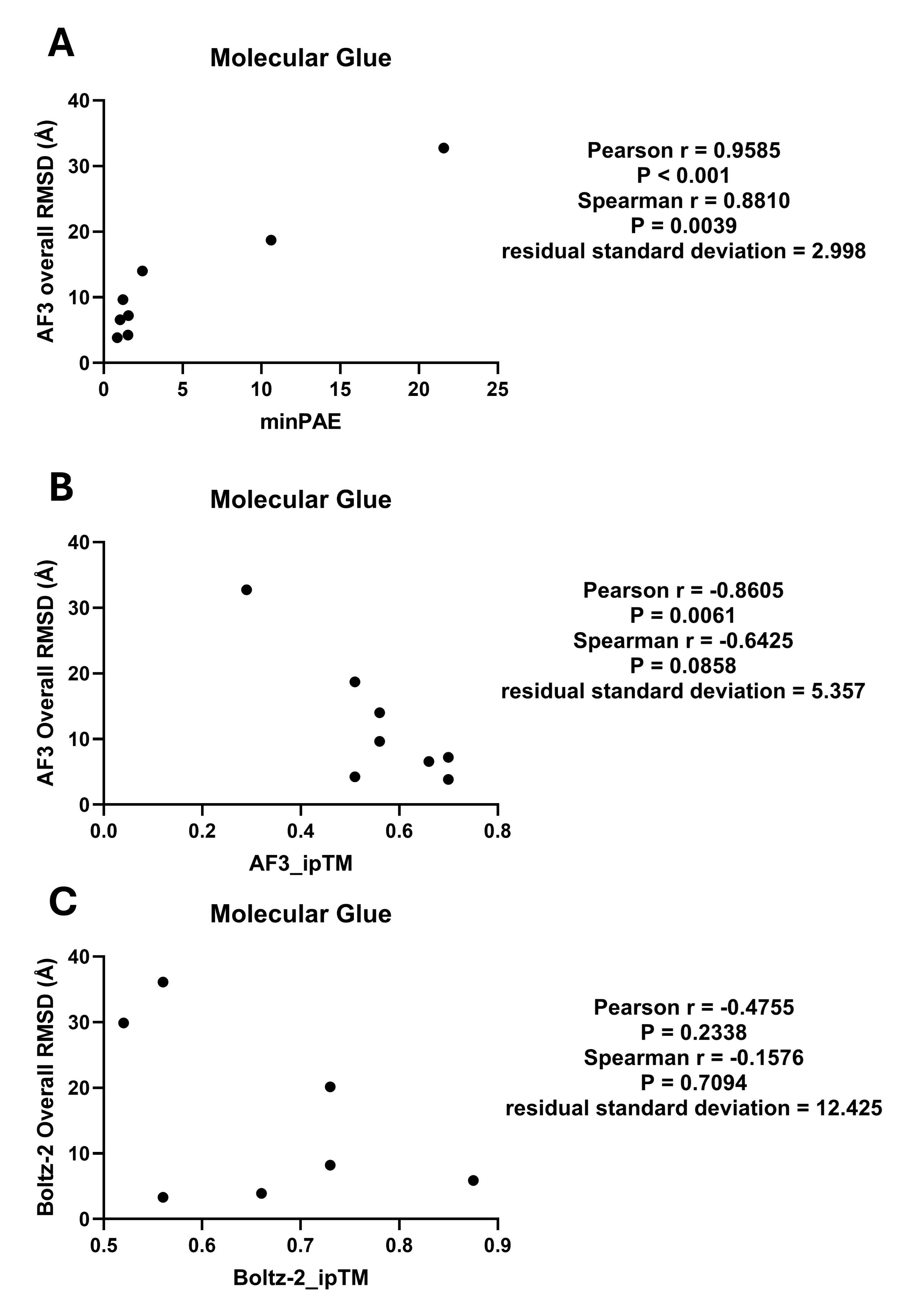

**Figure S17**. Relationship between predicted pose accuracy and model confidence for molecular glue. (**A**) AF3 overall ligand RMSD versus minPAE. (**B**) AF3 overall ligand RMSD versus ipTM. (**C**) Boltz-2 overall ligand RMSD versus ipTM. Lower RMSD indicates greater agreement with the experimental ligand pose.

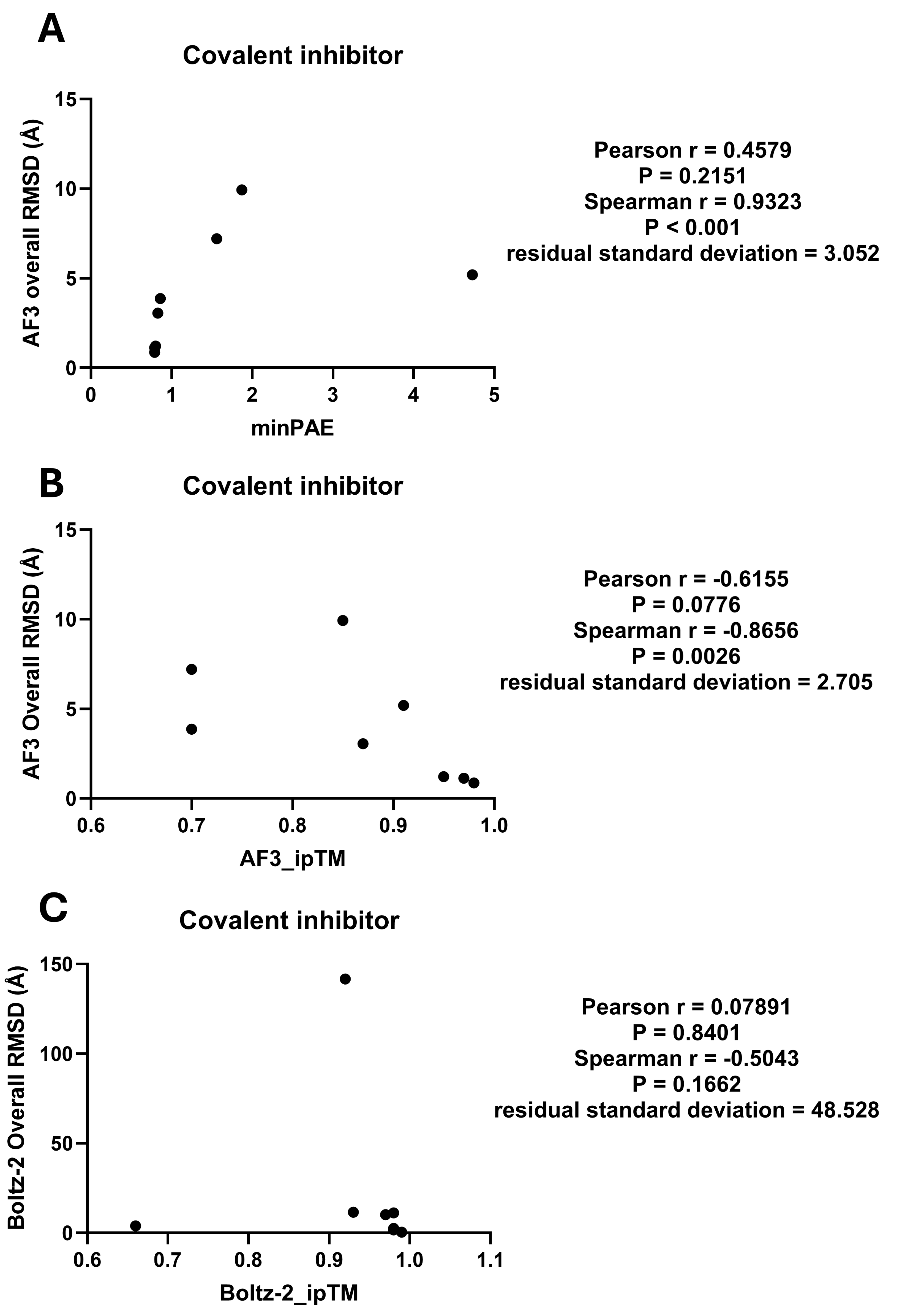

**Figure S18**. Relationship between predicted pose accuracy and model confidence for covalent inhibitor. (**A**) AF3 overall ligand RMSD versus minPAE. (**B**) AF3 overall ligand RMSD versus ipTM. (**C**) Boltz-2 overall ligand RMSD versus ipTM. Lower RMSD indicates greater agreement with the experimental ligand pose.

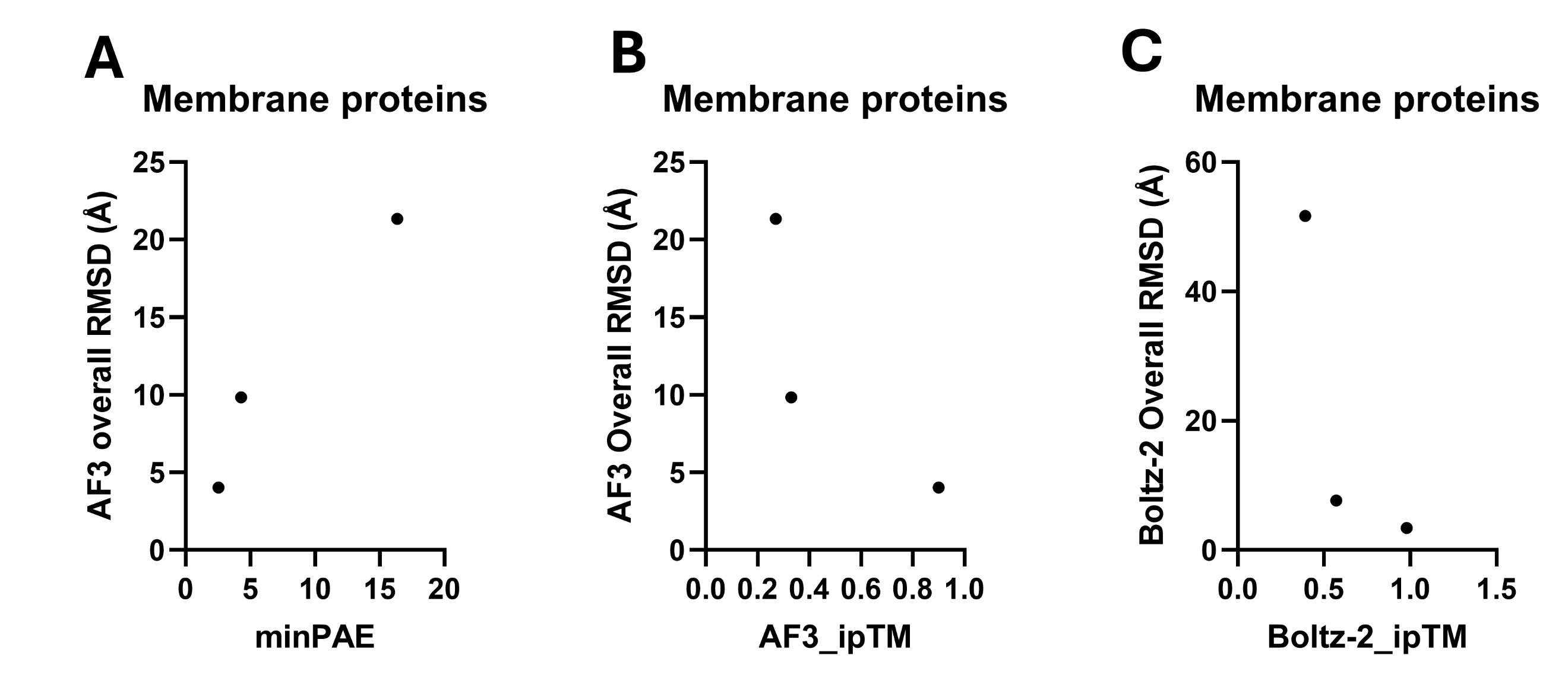

**Figure S19**. Relationship between predicted pose accuracy and model confidence for membrane proteins. (**A**) AF3 overall ligand RMSD versus minPAE. (**B**) AF3 overall ligand RMSD versus ipTM. (**C**) Boltz-2 overall ligand RMSD versus ipTM. Lower RMSD indicates greater agreement with the experimental ligand pose.

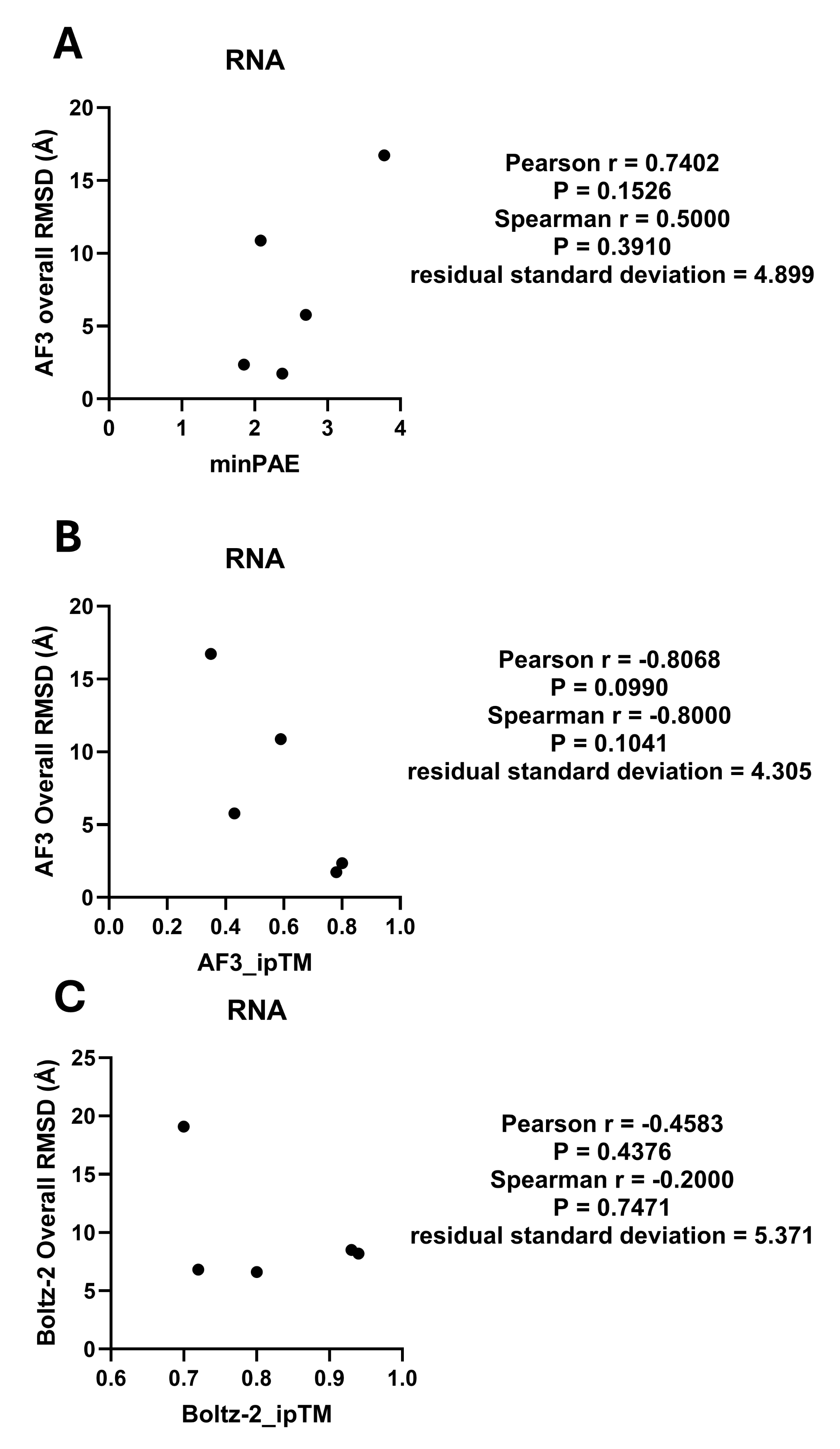

**Figure S20**. Relationship between predicted pose accuracy and model confidence for RNA. (**A**) AF3 overall ligand RMSD versus minPAE. (**B**) AF3 overall ligand RMSD versus ipTM. (**C**) Boltz-2 overall ligand RMSD versus ipTM. Lower RMSD indicates greater agreement with the experimental ligand pose.

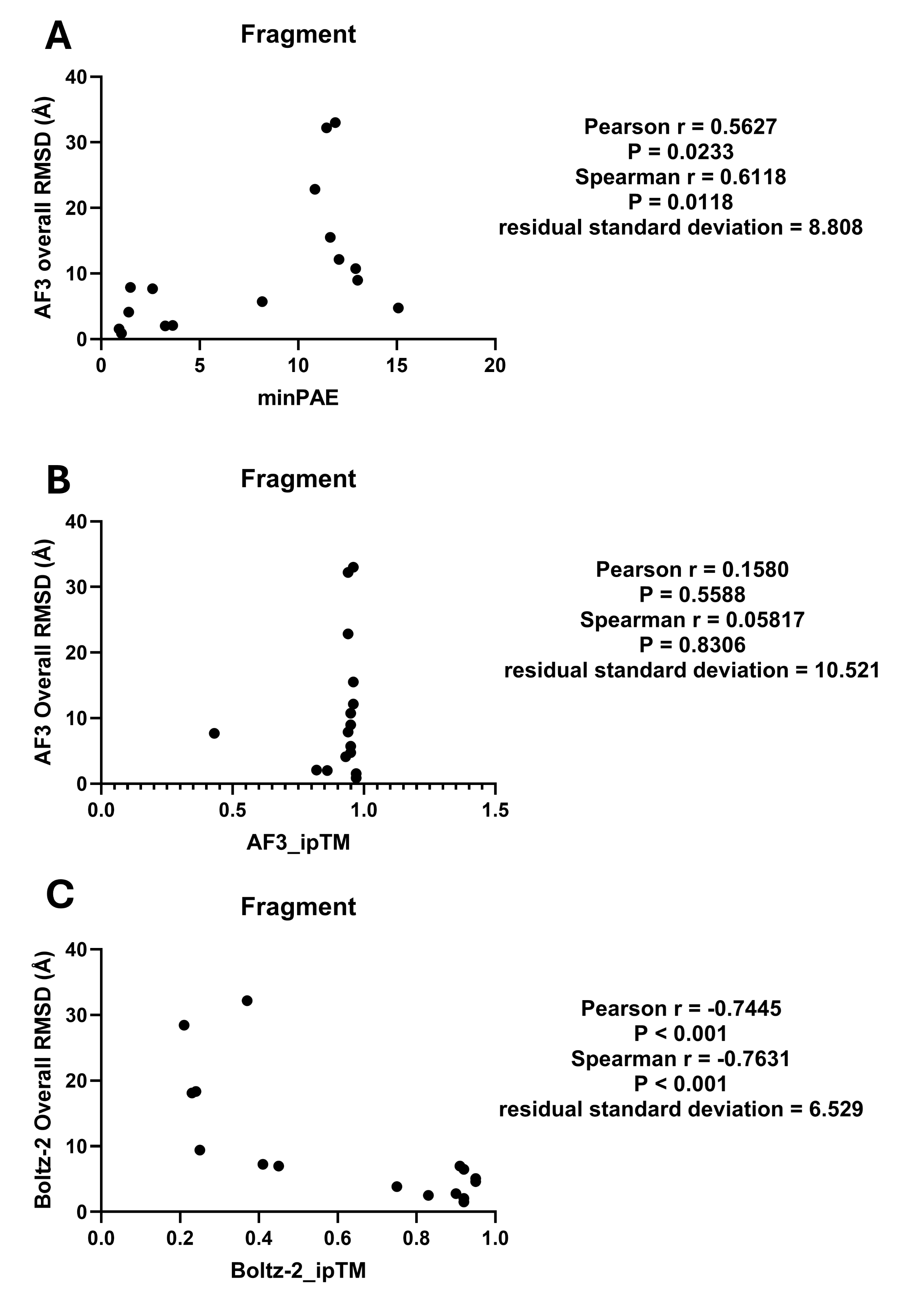

**Figure S21**. Relationship between predicted pose accuracy and model confidence for fragment. (**A**) AF3 overall ligand RMSD versus minPAE. (**B**) AF3 overall ligand RMSD versus ipTM. (**C**) Boltz-2 overall ligand RMSD versus ipTM. Lower RMSD indicates greater agreement with the experimental ligand pose.

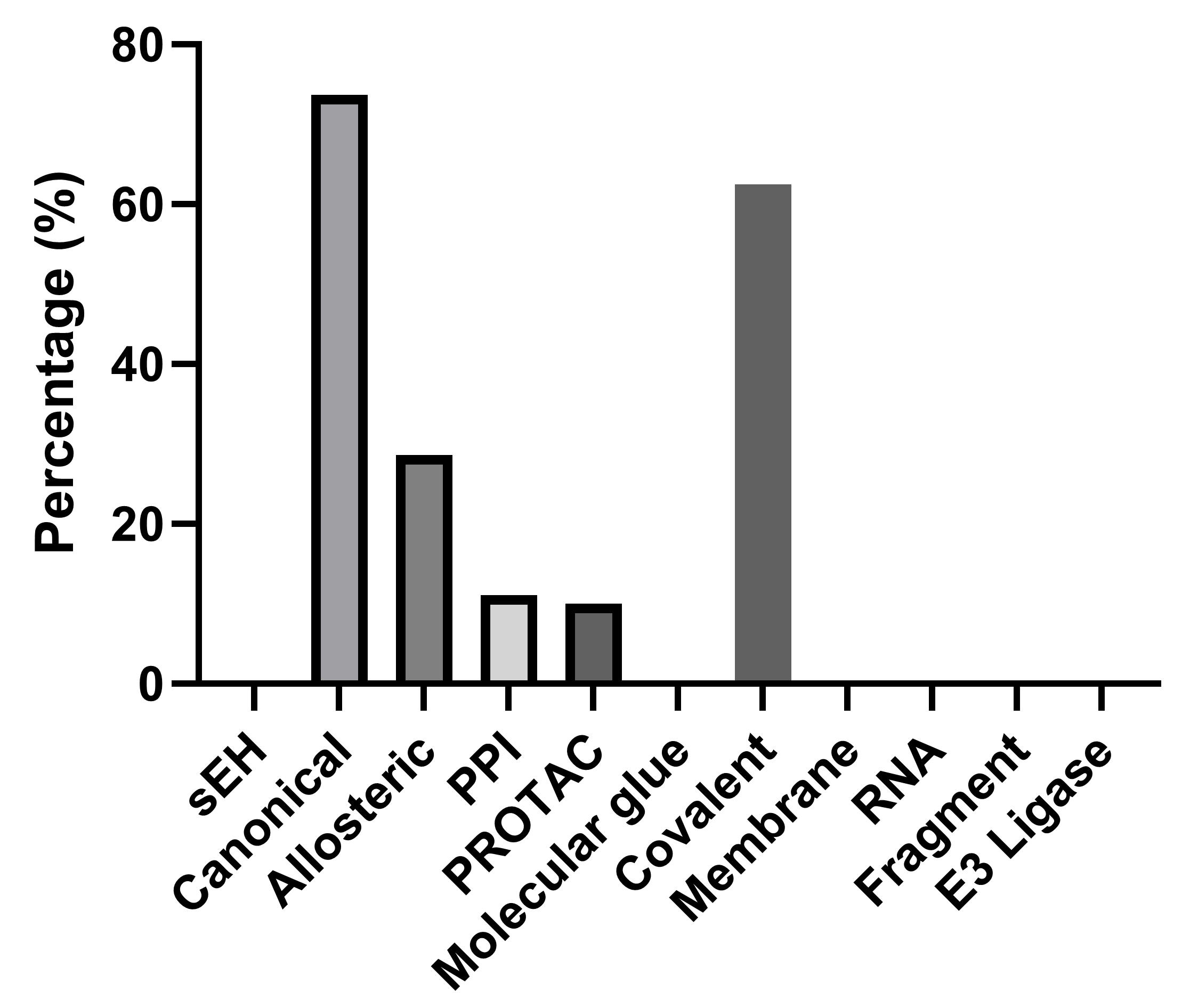

**Figure S22. Distribution of low-confidence-error AF3 predictions across ligand-binding modalities.** Percentage of examples within each ligand modality with AF3 minPAE < 0.85 Å. Canonical and covalent ligand-binding complexes showed the highest proportions of predictions within the minPAE < 0.85 Å range.

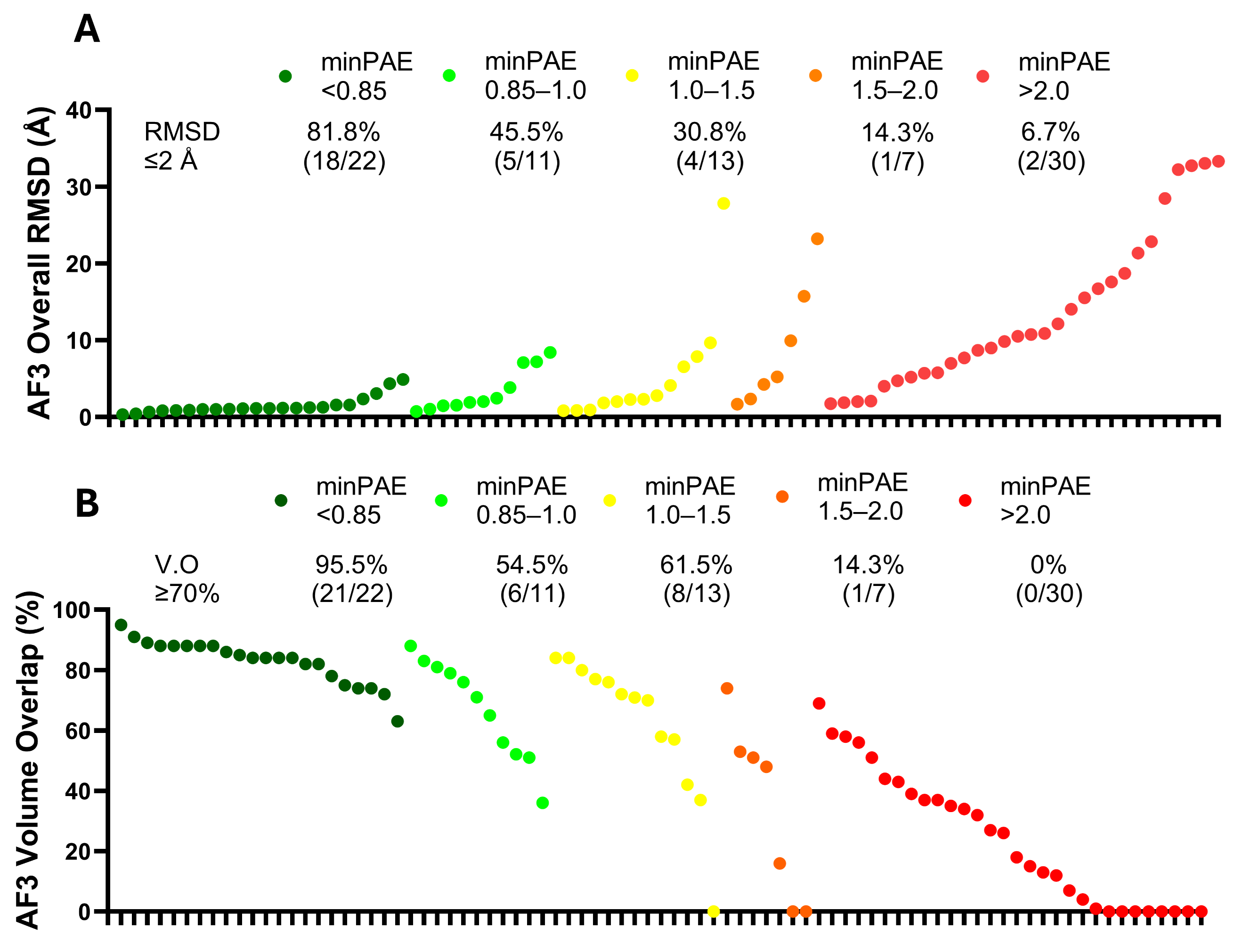

**Figure S23**. (**A**) AF3 minPAE versus ligand overall RMSD. (**B**) AF3 minPAE versus ligand volume overlap (%).

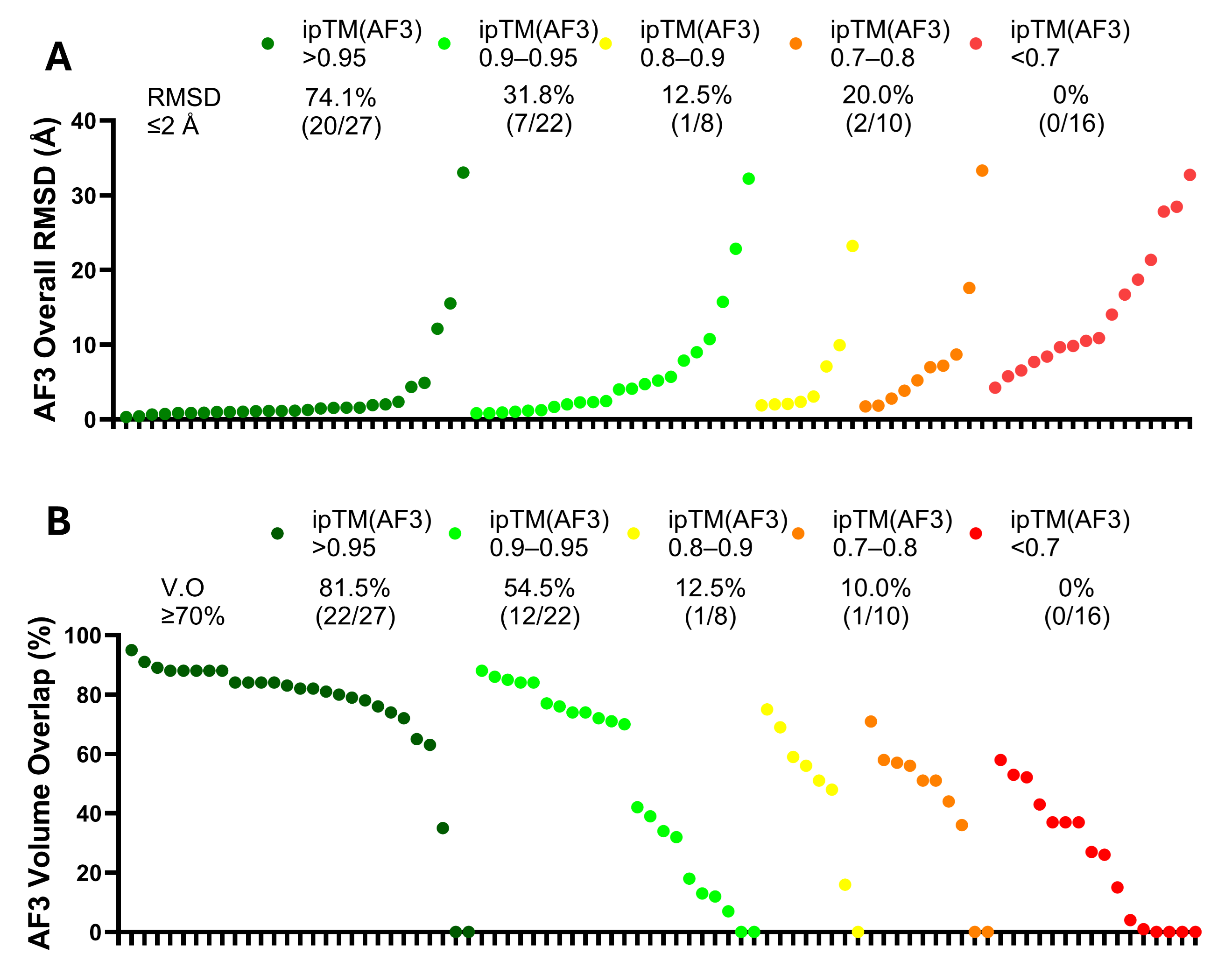
 **Figure S24**. (**A**) AF3 ipTM versus ligand overall RMSD. (**B**) AF3 ipTM versus ligand volume overlap (%).

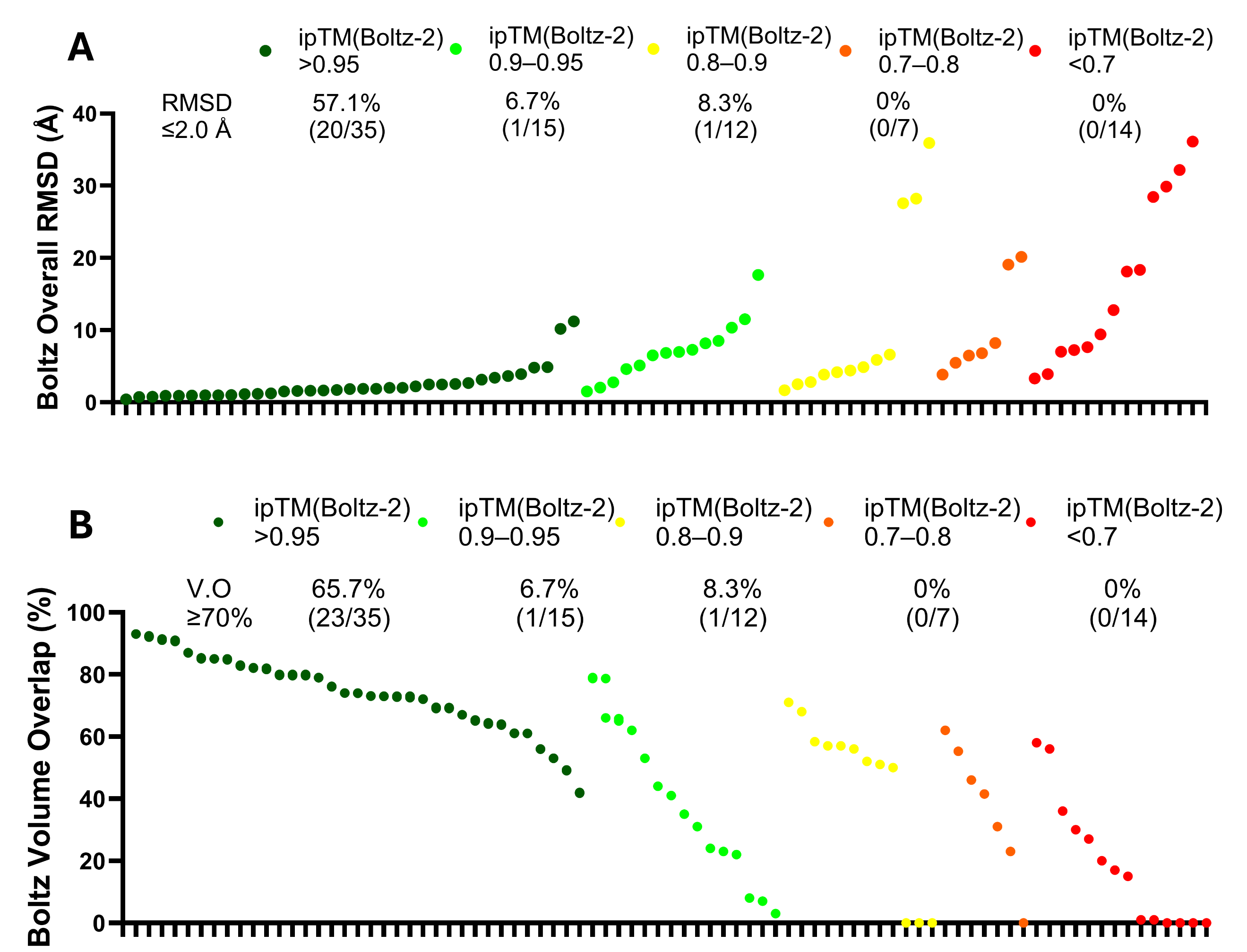

**Figure S25**. (**A**) Boltz-2 ipTM versus ligand overall RMSD. (**B**) Boltz-2 ipTM versus ligand volume overlap (%).

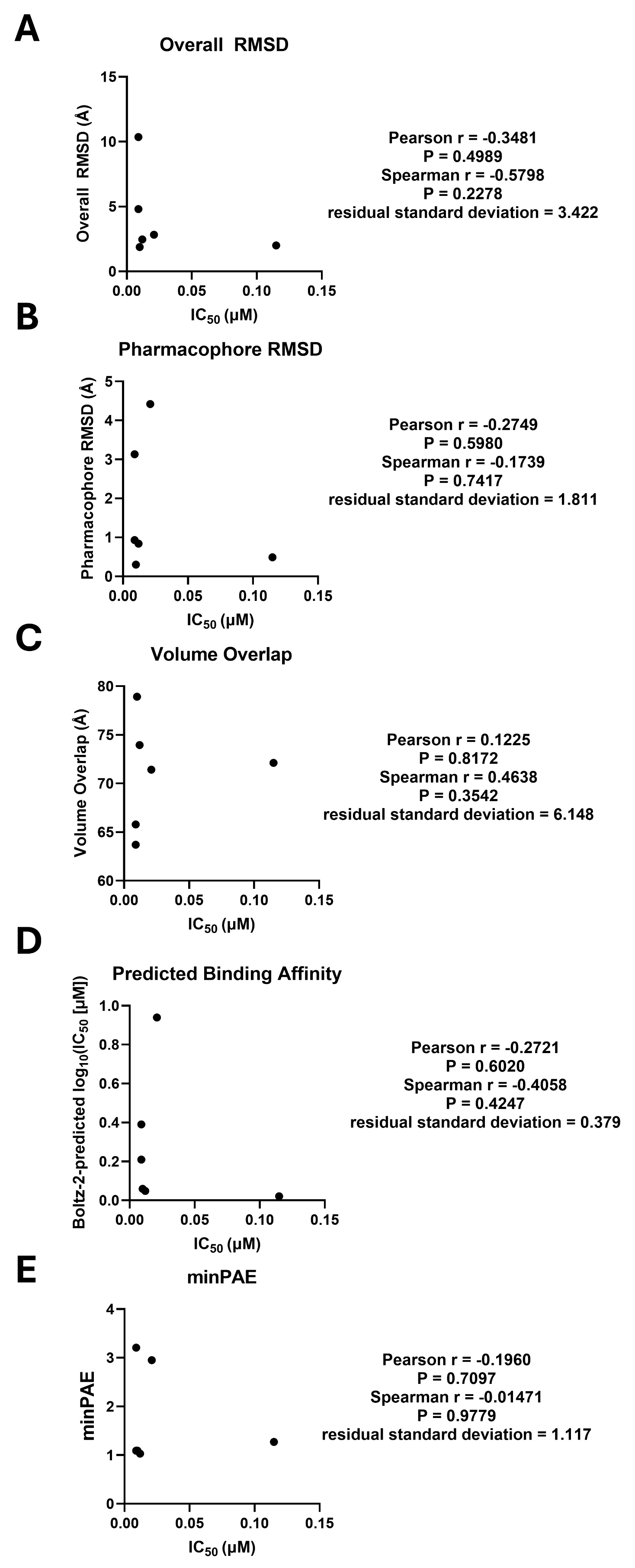

**Figure S26**. **Relationship between experimental IC_50_ and AF3/Boltz-2 prediction metrics for the sEH inhibitor series.** (**A**) Experimental IC_50_ versus AF3 overall ligand RMSD. (**B**) Experimental IC_50_ versus AF3 pharmacophore RMSD. (**C**) Experimental IC_50_ versus AF3 ligand volume overlap. (**D**) Experimental IC_50_ versus Boltz-2-predicted binding affinity. (**E**) Experimental IC_50_ versus AF3 minPAE.

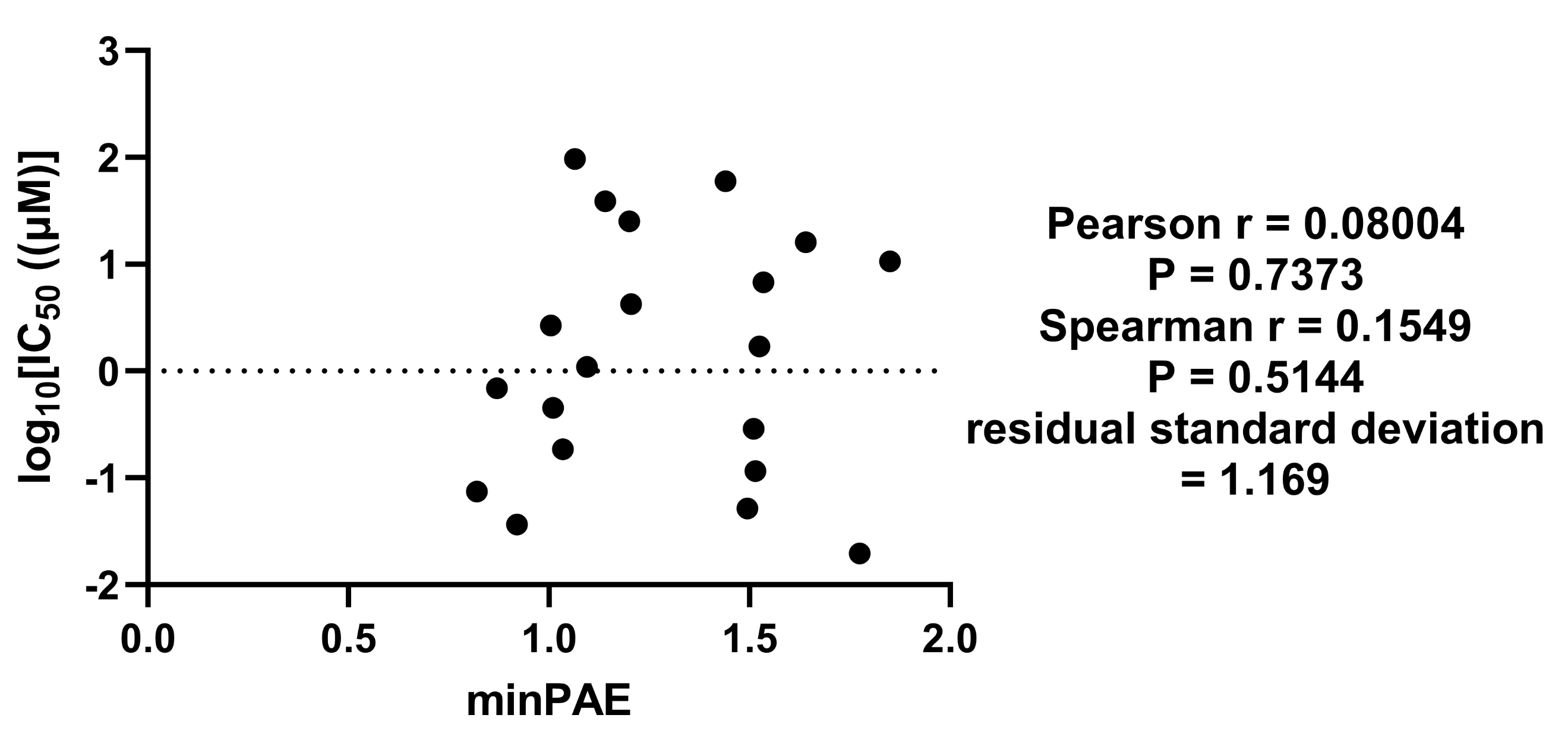

**Figure S27**. Relationship between minPAE and experimental IC_50_ for the Mpro series.

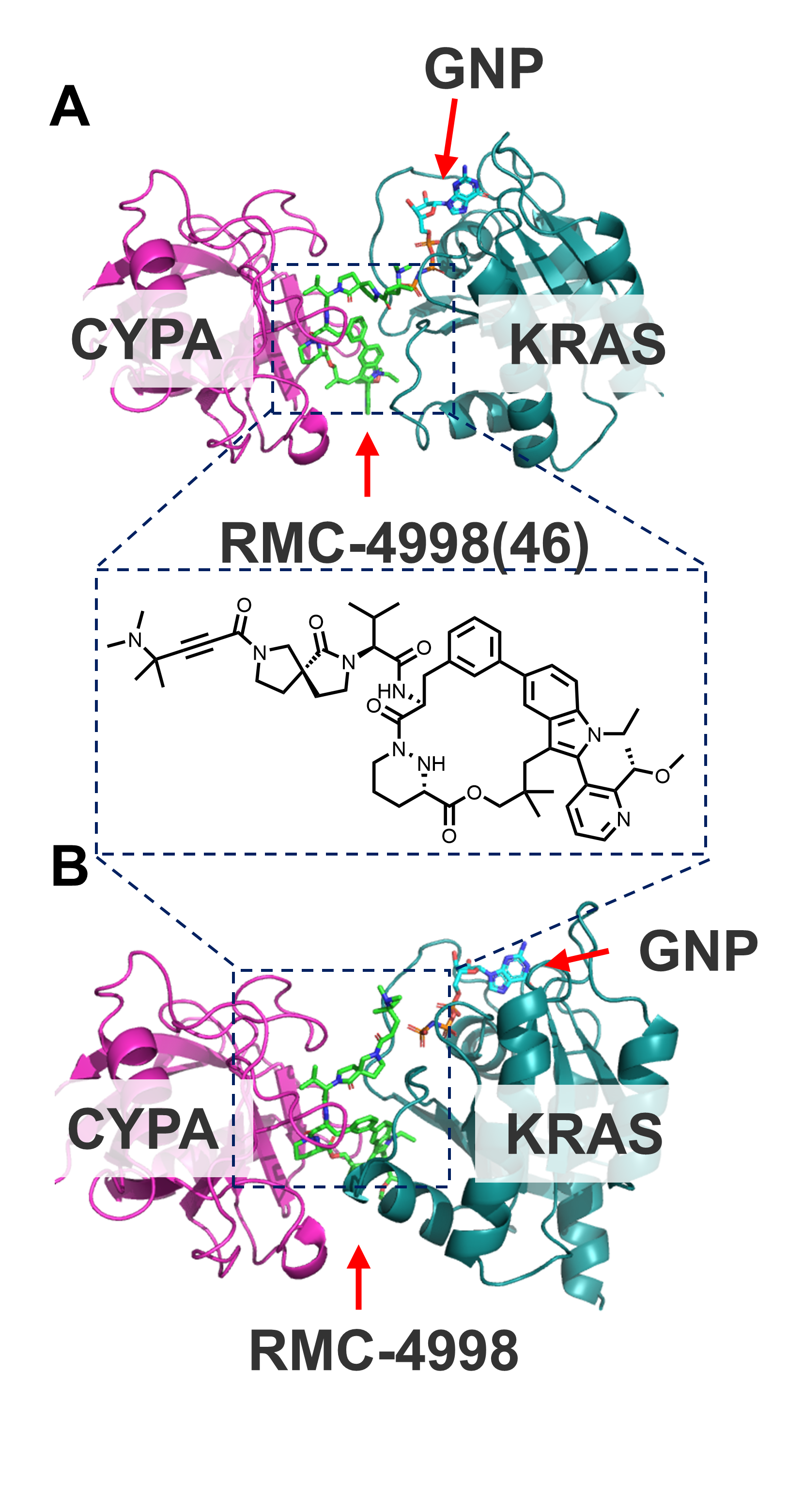

**Figure S28**. Representative example of an experimentally observed and AF3-predicted molecular-glue ternary complex. (**A**) Experimental structure of the RMC-4998–KRAS G12C–CypA complex (46: 8G9P). (**B**) Corresponding AF3 prediction for 46.

| **Table S1.** The numbers and their related PDB ID and the annotations of protein | | |
| --- | --- | --- |
| S. No. | Samples (PDB) | Annotation |
| 1 | sEH-P093D14 | Crystal structure of sEH in complex with HP-2 |
| 2 | sEH-P092B12 | Crystal structure of sEH in complex with FP-11 |
| 3 | sEH-P093E07 | Crystal structure of sEH in complex with FP-17 |
| 4 | sEH-P096D18 | Crystal structure of sEH in complex with FP-22 |
| 5 | sEH-P092O17 | Crystal structure of sEH in complex with HP-1 |
| 6 | sEH-P095M12 | Crystal structure of sEH in complex with HP-3 |
| 7 | 8BM2^1^ | Crystal structure of JAK2 JH1 in complex with Gandotinib |
| 8 | 8BPW^1^ | Crystal structure of JAK2 JH1 in complex with Lestaurtinib |
| 9 | 8BX6^1^ | Crystal structure of JAK2 JH1 in complex with Cerdulatinib |
| 10 | 8BX9^1^ | Crystal structure of JAK2 JH1 in complex with Ilginatinib |
| 11 | 8BXC^1^ | Crystal structure of JAK2 JH1 in complex with Itacitinib |
| 12 | 8BXH^1^ | Crystal structure of JAK2 JH1 in complex with Momelotinib |
| 13 | 8S9A^2^ | Crystal structure of the TYK2 pseudokinase domain in complex with TAK-279 |
| 14 | 8C7Y^3^ | Crystal structure of BRAF V600E in complex with a hybrid compound 6 |
| 15 | 8FV3^4^ | EGFR(T790M/V948R) in complex with compound 1 (LN4503) |
| 16 | 8FV4^4^ | EGFR(T790M/V948R) in complex with compound 2 (LN5993) |
| 17 | 8U8J^5^ | Co-crystal structure of phosphorylated ERK2 in complex with ERK1/2 inhibitor #16 |
| 18 | 8ZQW^6^ | The crystal structure of PDE4D with isoaurostatin derivatives 2-9 |
| 19 | 9BYJ^7^ | Crystal Structure of Hck in complex with the Src-family kinase inhibitor A-419259 |
| 20 | 9CJ1^8^ | Dual phosphorylated human p38 alpha bound to nilotinib |
| 21 | 9CJ4^8^ | Dual phosphorylated human p38 alpha bound to BIRB796 |
| 22 | 8HUK^9^ | X-ray structure of human PPAR alpha ligand binding domain-lanifibranor-SRC1 coactivator peptide co-crystals obtained by soaking |
| 23 | 8HUQ^9^ | X-ray structure of human PPAR alpha ligand binding domain-elafibranor-SRC1 coactivator peptide co-crystals obtained by soaking |
| 24 | 8UOB^10^ | SARS-CoV-2 Papain-like protease (PLpro) with Inhibitor Jun12682 |
| 25 | 9CSY^11^ | SARS-CoV-2 papain-like protease (PLpro) bound to PF-07957472 |
| 26 | 8PFO^12^ | Crystal structure of WRN helicase domain in complex with HRO761 |
| 27 | 7GQU^13^ | Crystal Structure of Werner helicase fragment 517-945 in covalent complex with N-[(E,1S)-1-cyclopropyl-3-methylsulfonylprop-2-enyl]-2-(1,1-difluoroethyl)-4-phenoxypyrimidine-5-carboxamide |
| 28 | 8A2D^14^ | EGFR kinase domain (L858R/V948R) in complex with 2-[4-(difluoromethyl)-6-[2-[4-[[4-(hydroxymethyl)-1-piperidyl]methyl]phenyl]ethynyl]-7-methyl-indazol-2-yl]-2-spiro[6,7-dihydropyrrolo[1,2-c]imidazole-5,1'-cyclopropane]-1-yl-N-thiazol-2-yl-acetamide |
| 29 | 8JUE^15^ | Crystal structure of glutaminase C in complex with compound 11 |
| 30 | 8AZV-GDP^16^ |  |
|  | 8AZV-OFU^16^ | KRAS in complex with BI-2865 |
| 31 | 8S0O^17^ | A fragment-based inhibitor of SHP2 |
| 32 | 8C14^18^ | Aurora A kinase in complex with TPX2-inhibitor 9 |
| 33 | 9X53^19^ | Crystal structure of RhoGDI2 in complex with Compound HR3119 |
| 34 | 9MJM^20^ | SOS1 IN COMPLEX WITH AN INHIBITOR |
| 35 | 8XGV^21^ | Optimization Efforts for Identification of Novel Highly Potent Keap1-Nrf2 Protein-Protein Interaction (PPI) Inhibitors |
| 36 | 8T5G^22^ | SOS2 co-crystal structure with fragment bound (compound 12) |
| 37 | 8T5M^22^ | SOS2 crystal structure with fragment bound (compound 14) |
| 38 | 8T5R^22^ | SOS2 crystal structure with fragment bound (compound 13) |
| 39 | 8UC9^22^ | SOS2 co-crystal structure with fragment bound (compound 9) |
| 40 | 8UH0^22^ | SOS2 co-crystal structure with fragment bound (compound 10) |
| 41 | 9D0X_CRBN^23^ | Cryo-EM structure of CDK2/CyclinE1 in complex with CRBN/DDB1 and Cpd 4 (local mask) |
|  | 9D0X_CDK2^23^ |  |
| 42 | 8QVU-kRas^24^ | Crystal Structure of ligand ACBI3 in complex with KRAS G12D C118S GDP and pVHL:ElonginC:ElonginB complex |
|  | 8QVU-suppressor^24^ |  |
| 43 | 9RK8-kRas^25^ | Crystal Structure of compound 3-mediated ternary complex of KRAS G12V C118S GDP with pVHL:ElonginC:ElonginB |
|  | 9RK8-suppressor^25^ |  |
| 44 | 9L6F-Kras^26^ | Crystal structure of KRas G12D (GDP) in complex with ASP3082 |
|  | 9L6F-suppressor^26^ |  |
| 45 | 9BG0-nRas^27^ | Tri-complex of Daraxonrasib (RMC-6236), NRAS WT, and CypA |
|  | 9BG0-CypA^27^ |  |
| 46 | 8G9P-CypA^28^ | Tricomplex of RMC-4998, KRAS G12C, and CypA |
|  | 8G9P-kRas^28^ |  |
| 47 | 8OV6^29^ | Ternary structure of intramolecular bivalent glue degrader IBG1 bound to BRD4 and DCAF16:DDB1deltaBPB |
| 48 | 9DUR_CRBN^30^ | Cryo-EM Structure of CRBN:dHTC1:ENL YEATS |
|  | 9DUR_ENL^30^ |  |
| 49 | 8TZX^31^ | Ternary complex structure of Cereblon-DDB1 bound to WIZ(ZF7) and the molecular glue dWIZ-1 |
| 50 | 8UVM^10^ | SARS-CoV-2 papain-like protease (PLpro) complex with covalent inhibitor Jun11313 |
| 51 | 8V3A^32^ | Crystal structure of KRAS-G12C (GDP-bound) in complex with BBO-8520 |
| 52 | 8V39^32^ | Crystal structure of active KRAS-G12C (GMPPNP-bound) in complex with BBO-8520 |
| 53 | 8V4U^33^ | Structure of SARS-CoV-2 main protease in complex with a covalent inhibitor |
| 54 | 8JHL^34^ | GDP-bound KRAS G12D in complex with YK-8S |
| 55 | 8ZKN^35^ | CryoEM structure of Thyroid Hormone Transporter MCT8 bound with silychristin |
| 56 | 9CY3^36^ | Outward-facing Atorvastatin-bound OATP1B1 with sybody Sb5 |
| 57 | 9LK8^37^ | Cryo-EM structure of GAT3 |
| 58 | 8R62^38^ | Solution structure of Risdiplam bound to the RNA duplex formed upon 5'-splice site recognition |
| 59 | 8QMH^39^ | Crystal structure of RNA G2C4 repeats in complex with small synthetic molecule ANP77 |
| 60 | 8R8P^38^ | Solution structure of SMN-CX bound to the RNA helix formed upon SMN2 exon7 5'-splice site recognition |
| 61 | 8ZNQ^40^ | Solution structure of the complex of naphthyridine-azaquinolone and an RNA with ACG/AUA motif |
| 62 | 9CPD^41^ | Structures of small molecules bound to RNA repeat expansions that cause Huntington's disease-like 2 and myotonic dystrophy type 1 |
| 63 | 9IO0^42^ | INTERACTION BETWEEN A FLUOROQUINOLONE DERIVATIVE KG022 AND RNAS: EFFECT OF BASE PAIRS 5' ADJACENT TO THE BULGE OUT ESIDUES |
| 64 | 7GRE^43^ | Crystal structure of SARS-CoV-2 main protease in complex with cpd-1 |
| 65 | 7GRF^43^ | Crystal structure of SARS-CoV-2 main protease in complex with cpd-2 |
| 66 | 7GRJ^43^ | Crystal structure of SARS-CoV-2 main protease in complex with cpd-6 |
| 67 | 7GRN^43^ | Crystal structure of SARS-CoV-2 main protease in complex with cpd-10 |
| 68 | 7GRS^43^ | Crystal structure of SARS-CoV-2 main protease in complex with cpd-15 |
| 69 | 7GRT^43^ | Crystal structure of SARS-CoV-2 main protease in complex with cpd-16 |
| 70 | 7GRU^43^ | Crystal structure of SARS-CoV-2 main protease in complex with cpd-17 |
| 71 | 7GRZ^43^ | Crystal structure of SARS-CoV-2 main protease in complex with cpd-22 |
| 72 | 7GS0^43^ | Crystal structure of SARS-CoV-2 main protease in complex with cpd-23 |
| 73 | 9BVE^44^ | Identification of multiple ligand hotspots on SOS2, compound 9 |
| 74 | 4CI1_TEST1^45^ | Structure of the DDB1-CRBN E3 ubiquitin ligase bound to thalidomide |
| 75 | 4CI1_TEST2^45^ |  |
| 76 | 7GLV^46^ | Crystal Structure of SARS-CoV-2 main protease in complex with MAT-POS-a54ce14d-2 (Mpro-P2031) |
| 77 | 7GAW^46^ | Crystal Structure of SARS-CoV-2 main protease in complex with MAT-POS-e194df51-1 (SARS2_MproA-x0862) |
| 78 | 7GKS^46^ | Crystal Structure of SARS-CoV-2 main protease in complex with MAT-POS-4223bc15-11 (Mpro-P1062) |
| 79 | 7GNT^46^ | Crystal Structure of SARS-CoV-2 main protease in complex with MAT-POS-50a80394-2 (Mpro-P3054) |
| 80 | 7GLG^46^ | Crystal Structure of SARS-CoV-2 main protease in complex with MAT-POS-5cd9ea36-18 (Mpro-P1980) |
| 81 | 7GIU^46^ | Crystal Structure of SARS-CoV-2 main protease in complex with RAL-THA-05e671eb-10 (Mpro-P0130) |
| 82 | 7GM1^46^ | Crystal Structure of SARS-CoV-2 main protease in complex with MAT-POS-2e8b2191-10 (Mpro-P2072) |
| 83 | 7GNA^46^ | Crystal Structure of SARS-CoV-2 main protease in complex with ALP-POS-4483ae88-4 (Mpro-P2415) |
| 84 | 7GIX^46^ | Crystal Structure of SARS-CoV-2 main protease in complex with VLA-UNK-cf7facf1-1 (Mpro-P0143) |
| 85 | 7GHM^46^ | Crystal Structure of SARS-CoV-2 main protease in complex with ERI-UCB-ce40166b-17 (Mpro-P0008) |
| 86 | 7GG2^46^ | Crystal Structure of SARS-CoV-2 main protease in complex with MAT-POS-8a69d52e-7 (Mpro-x12073) |
| 87 | 7GMR^46^ | Crystal Structure of SARS-CoV-2 main protease in complex with PET-UNK-7fb4f80a-2 (Mpro-P2214) |
| 88 | 7GE3^46^ | Crystal Structure of SARS-CoV-2 main protease in complex with TRY-UNI-714a760b-3 (Mpro-x11317) |
| 89 | 7GMP^46^ | Crystal Structure of SARS-CoV-2 main protease in complex with MAT-POS-90fd5f68-7 (Mpro-P2207) |
| 90 | 7GE2^46^ | Crystal Structure of SARS-CoV-2 main protease in complex with EDJ-MED-6af13d92-1 (Mpro-x11313) |
| 91 | 7GE4^46^ | Crystal Structure of SARS-CoV-2 main protease in complex with TRY-UNI-714a760b-19 (Mpro-x11318) |
| 92 | 7GGC^46^ | Crystal Structure of SARS-CoV-2 main protease in complex with MAT-POS-044491d2-3 (Mpro-x12321) |
| 93 | 7GFJ^46^ | Crystal Structure of SARS-CoV-2 main protease in complex with ALP-POS-c0c213c9-1 (Mpro-x11757) |
| 94 | 7GCK^46^ | Crystal Structure of SARS-CoV-2 main protease in complex with BAR-COM-0f94fc3d-48 (Mpro-x10638) |
| 95 | 7GBV^46^ | Crystal Structure of SARS-CoV-2 main protease in complex with JAN-GHE-83b26c96-22 (Mpro-x10422) |

| **Table S2.** The Overall RMSD, Pharm RMSD, Volume Overlap of sEH ligand predictions (AF3 and Boltz-2) & the confidence metrics of AF3 and Boltz-2 | | | | | | | | | | |
| --- | --- | --- | --- | --- | --- | --- | --- | --- | --- | --- |
| S. No. | Structure | AF3 (2021. Sep. 30) | | | Boltz-2 (2023. Jun.1) | | | AF3 | | Boltz-2 |
|  |  | Overall RMSD | Pharm RMSD | Volume Overlap | Overall RMSD | Pharm RMSD | Volume Overlap | ipTM | minPAE(A) |  |
|  |  |  |  |  |  |  |  |  |  | ipTM |
|  |  | Mean ± SD | Mean ± SD | Mean ± SD | Mean ± SD | Mean ± SD | Mean ± SD |  |  |  |
| 1 | 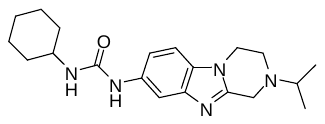 | 2.00±0.16 | 0.54±0.02 | 0.72±0.03 | 1.87±0.22 | 0.29±0.03 | 0.79±0.03 | 0.94 | 1.09 | 0.98 |
| 2 | 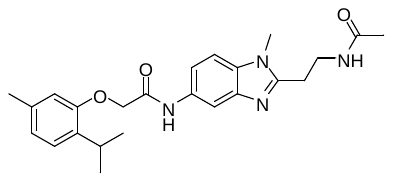 | 2.33±0.47 | 0.58±0.05 | 0.70±0.02 | 2.00±0.45 | 0.49±0.05 | 0.72±0.03 | 0.92 | 1.27 | 0.97 |
| 3 | 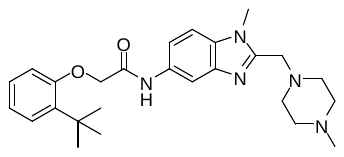 | 2.30±0.07 | 1.04±0.15 | 0.76±0.01 | 2.47±0.05 | 0.84±0.03 | 0.74±0.01 | 0.94 | 1.03 | 0.98 |
| 4 | 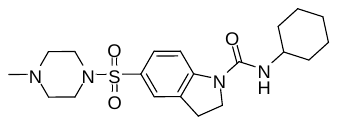 | 0.94±0.29 | 0.25±0.04 | 0.84±0.02 | 4.81±4.28 | 0.92±1.01 | 0.64±0.14 | 0.93 | 1.09 | 0.96 |
| 5 | 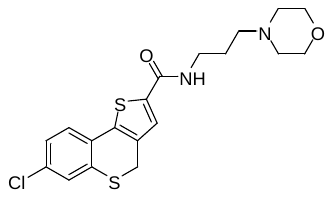 | 6.98±4.23 | 4.96±3.58 | 0.51±0.12 | 10.36±0.01 | 3.13±0.03 | 0.66±0.02 | 0.73 | 3.21 | 0.95 |
| 6 | 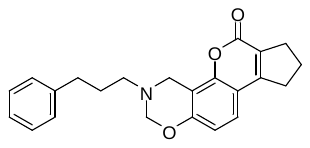 | 8.69±4.61 | 8.22±4.78 | 0.44±0.22 | 2.83±0.10 | 4.41±0.14 | 0.71±0.02 | 0.76 | 2.95 | 0.88 |

| **Table S3.** The Overall RMSD, Pharm RMSD, Volume Overlap of canonical ligand-binding complex predictions (AF3 and Boltz-2) & the confidence metrics of AF3 and Boltz-2 | | | | | | | | | | |
| --- | --- | --- | --- | --- | --- | --- | --- | --- | --- | --- |
| S. No. | Structure | AF3 (2021. Sep. 30) | | | Boltz-2 (2023. Jun.1) | | | AF3 | | Boltz-2 |
|  |  | Overall RMSD | Pharm RMSD | Volume Overlap | Overall RMSD | Pharm RMSD | Volume Overlap | ipTM | minPAE(A) | ipTM |
|  |  | Mean ± SD | Mean ± SD | Mean ± SD | Mean ± SD | Mean ± SD | Mean ± SD |  |  |  |
| 7 | 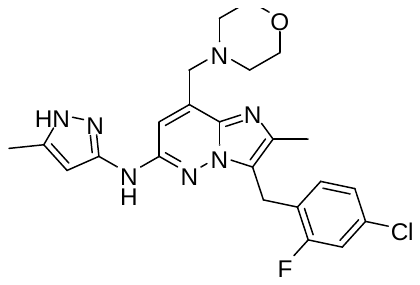 | 1.00±0.15 | 0.42±0.08 | 0.82±0.01 | 1.18±0.06 | 0.30±0.06 | 0.80±0.02 | 0.97 | 0.79 | 0.99 |
| 8 | 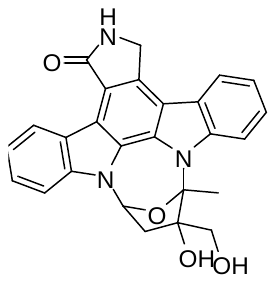 | 0.41±0.20 | 0.85±0.58 | 0.88±0.02 | 0.97±0.12 | 2.22±0.29 | 0.82±0.01 | 0.97 | 0.81 | 0.99 |
| 9 | 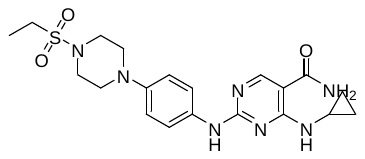 | 1.58±0.27 | 0.62±0.31 | 0.74±0.02 | 1.72±0.26 | 0.36±0.07 | 0.69±0.03 | 0.97 | 0.81 | 0.96 |
| 10 | 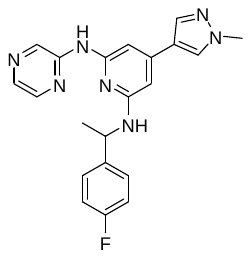 | 0.82±0.36 | 0.41±0.05 | 0.86±0.01 | 1.68±0.28 | 1.96±0.49 | 0.68±0.04 | 0.95 | 0.79 | 0.84 |
| 11 | 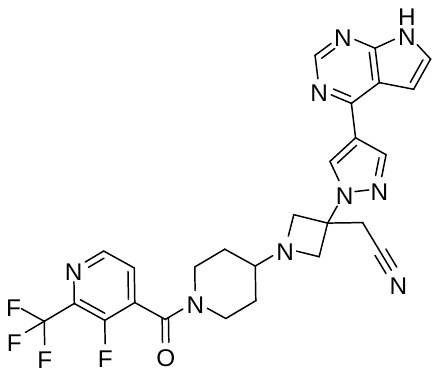 | 1.58±0.25 | 0.84±0.08 | 0.72±0.01 | 2.21±0.52 | 0.68±0.07 | 0.73±0.01 | 0.98 | 0.80 | 0.97 |
| 12 | 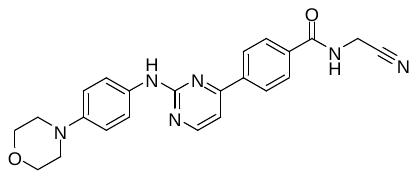 | 1.17±0.34 | 0.32±0.06 | 0.74±0.02 | 1.88±0.18 | 0.97±0.16 | 0.64±0.02 | 0.93 | 0.80 | 0.97 |
| 13 | 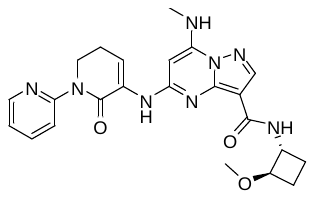 | 1.15±0.22 | 1.15±0.50 | 0.84±0.01 | 0.91±0.19 | 0.86±0.43 | 0.82±0.01 | 0.96 | 0.79 | 0.97 |
| 14 | 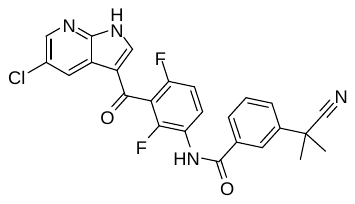 | 0.33±0.22 | 0.22±0.03 | 0.95±0.01 | 0.95±0.22 | 0.28±0.06 | 0.91±0.01 | 0.97 | 0.82 | 0.99 |
| 15 | 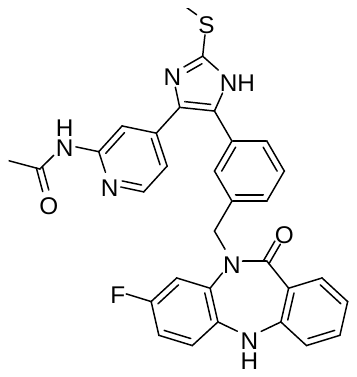 | 0.82±0.22 | 0.45±0.13 | 0.84±0.04 | 1.50±0.20 | 0.58±0.16 | 0.67±0.04 | 0.92 | 1.04 | 0.99 |
| 16 | 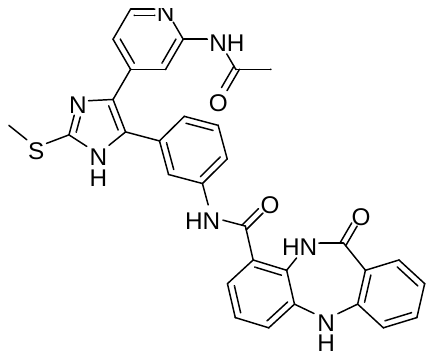 | 0.73±0.09 | 0.59±0.10 | 0.83±0.02 | 1.61±1.32 | 0.78±0.12 | 0.69±0.15 | 0.97 | 0.88 | 0.97 |
| 17 | 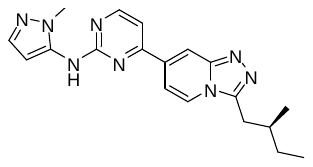 | 0.67±0.35 | 0.40±0.06 | 0.88±0.02 | 0.98±0.04 | 0.34±0.05 | 0.87±0.01 | 0.98 | 0.79 | 0.99 |
| 18 | 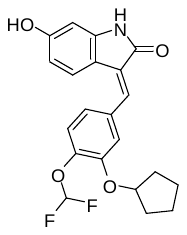 | 2.46±0.20 | 3.48±0.26 | 0.71±0.01 | 2.47±0.06 | 3.26±0.22 | 0.73±0.02 | 0.94 | 0.87 | 0.98 |
| 19 | 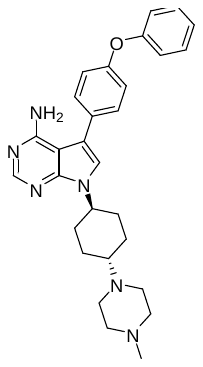 | 1.18±0.23 | 0.36±0.07 | 0.91±0.01 | 1.14±0.17 | 0.47±0.17 | 0.92±0.01 | 0.97 | 0.79 | 0.99 |
| 20 | 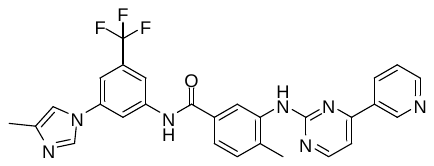 | 2.35±0.83 | 1.02±0.12 | 0.63±0.04 | 0.79±0.12 | 0.46±0.08 | 0.85±0.02 | 0.96 | 0.83 | 0.99 |
| 21 | 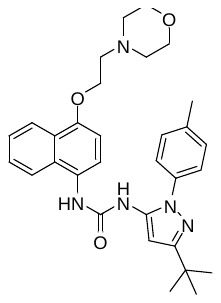 | 1.28±0.16 | 0.52±0.07 | 0.84±0.01 | 1.26±0.07 | 0.65±0.12 | 0.83±0.03 | 0.97 | 0.83 | 0.99 |
| 22 | 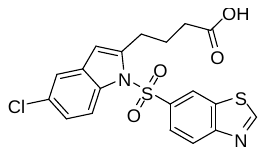 | 1.92±0.06 | 1.41±0.07 | 0.65±0.01 | 2.69±0.45 | 1.59±0.21 | 0.61±0.02 | 0.96 | 0.92 | 0.98 |
| 23 | 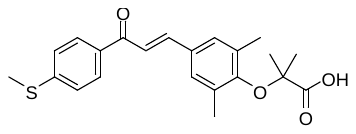 | 1.88±0.08 | 0.72±0.05 | 0.69±0.02 | 3.14±0.91 | 1.69±0.64 | 0.65±0.03 | 0.87 | 2.24 | 0.98 |
| 24 | 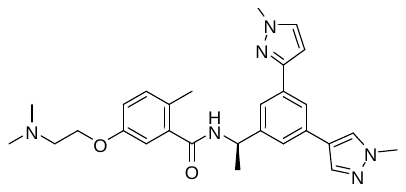 | 4.89±0.08 | 0.82±0.08 | 0.78±0.02 | 4.88±0.03 | 0.73±0.18 | 0.74±0.03 | 0.97 | 0.80 | 0.98 |
| 25 | 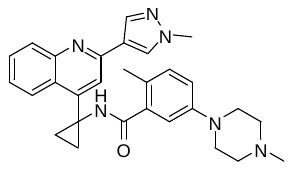 | 4.35±0.08 | 0.34±0.04 | 0.82±0.01 | 3.65±0.25 | 0.87±0.13 | 0.73±0.02 | 0.97 | 0.80 | 0.98 |

| **Table S4.** The Overall RMSD, Pharm RMSD, Volume Overlap of allosteric inhibitor predictions (AF3 and Boltz-2) & the confidence metrics of AF3 and Boltz-2 | | | | | | | | | | |
| --- | --- | --- | --- | --- | --- | --- | --- | --- | --- | --- |
| S. No. | Structure | AF3 (2021. Sep. 30) | | | Boltz-2 (2023. Jun.1) | | | AF3 | | Boltz-2 |
|  |  | Overall RMSD | Pharm RMSD | Volume Overlap | Overall RMSD | Pharm RMSD | Volume Overlap | ipTM | minPAE (A) | ipTM |
|  |  | Mean ± SD | Mean ± SD | Mean ± SD | Mean ± SD | Mean ± SD | Mean ± SD |  |  |  |
| 26 | 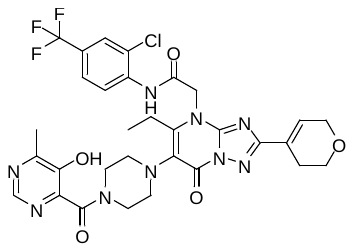 | 17.60±1.77 | 16.85±3.48 | 0.00±0.01 | 17.64±2.94 | 17.05±3.10 | 0.03±0.04 | 0.72 | 5.01 | 0.91 |
| 27 | 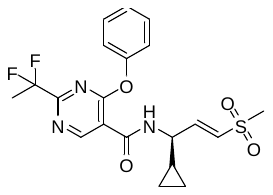 | 9.93±3.50 | 9.05±3.36 | 0.16±0.15 | 11.50±0.88 | 9.99±0.93 | 0.08±0.01 | 0.85 | 1.87 | 0.93 |
| 28 | 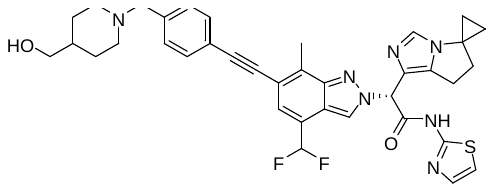 | 10.50±7.55 | 3.55±2.09 | 0.43±0.24 | 4.88±5.52 | 3.64±2.89 | 0.52±0.26 | 0.68 | 3.39 | 0.80 |
| 29 | 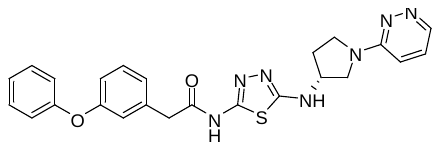 | 33.31±1.36 | 30.20±3.05 | 0.00±0.00 | 35.92±0.43 | 28.28±0.73 | 0.00±0.00 | 0.79 | 6.01, 4.59 | 0.81 |
| 30 | 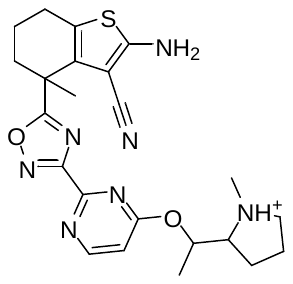 | 1.04±0.63 | 0.83±1.18 | 0.88±0.01 | 0.75±0.01 | 1.57±0.02 | 0.93±0.01 | 0.92 | 0.77 | 0.98 |
|  | 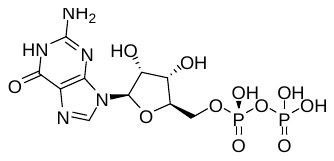 | 15.74±0.19 | 14.23±0.21 | 0.00±0.00 | 1.01±0.54 | 0.45±0.16 | 0.85±0.08 | 0.92 | 1.72 | 0.98 |
| 31 | 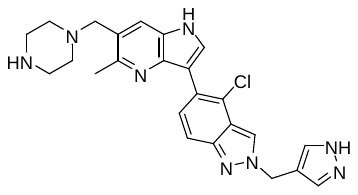 | 0.99±0.19 | 0.68±0.31 | 0.89±0.01 | 3.92±0.40 | 4.11±0.43 | 0.61±0.01 | 0.97 | 0.84 | 0.97 |

| **Table S5.** The Overall RMSD, Pharm RMSD, Volume Overlap of PPI predictions (AF3 and Boltz-2) & the confidence metrics of AF3 and Boltz-2 | | | | | | | | | | |
| --- | --- | --- | --- | --- | --- | --- | --- | --- | --- | --- |
| S. No. | Structure | AF3 (2021. Sep. 30) | | | Boltz-2 (2023. Jun.1) | | | AF3 | | Boltz-2 |
|  |  | Overall RMSD | Pharm RMSD | Volume Overlap | Overall RMSD | Pharm RMSD | Volume Overlap | ipTM | minPAE(A) | ipTM |
|  |  | Mean ± SD | Mean ± SD | Mean ± SD | Mean ± SD | Mean ± SD | Mean ± SD |  |  |  |
| 32 | 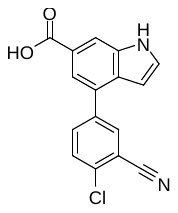 | 1.04±0.17 | 1.43±0.29 | 0.79±0.04 | 1.58±0.48 | 1.25±0.31 | 0.73±0.04 | 0.96 | 0.99 | 0.97 |
| 33 | 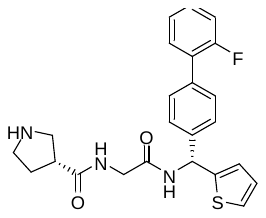 | 5.24±2.26 | 6.01±2.14 | 0.51±0.06 | 7.28±0.12 | 8.09±0.26 | 0.22±0.01 | 0.77 | 1.95 | 0.95 |
| 34 | 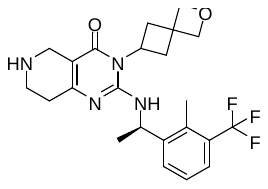 | 1.09±0.08 | 0.93±0.12 | 0.84±0.02 | 0.92±0.10 | 0.56±0.16 | 0.56±0.16 | 0.97 | 0.84 | 0.96 |
| 35 | 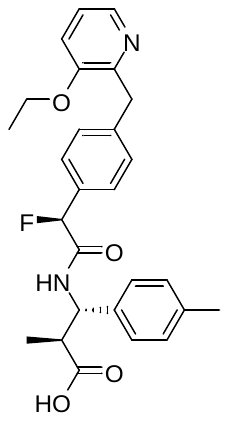 | 2.00±0.11 | 0.69±0.56 | 0.81±0.00 | 6.84±0.11 | 8.08±0.55 | 0.53±0.06 | 0.96 | 0.87 | 0.95 |
| 36 | 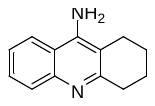 | 4.11±1.41 | 4.67±0.08 | 0.77±0.03 | 2.04±1.51 | 1.43±0.93 | 0.79±0.04 | 0.93 | 1.40 | 0.92 |
| 37 | 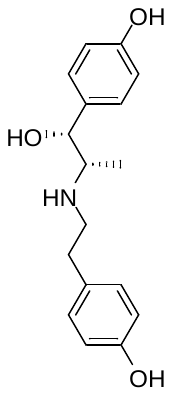 | 7.89±0.18 | 9.81±0.26 | 0.42±0.02 | 6.99±1.56 | 3.14±2.07 | 0.41±0.04 | 0.94 | 1.48 | 0.91 |
| 38 | 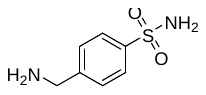 | 2.01±0.20 | 2.42±0.30 | 0.59±0.04 | 1.50±1.12 | 1.96±1.51 | 0.65±0.11 | 0.86 | 3.25 | 0.92 |
| 39 | 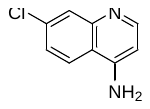 | 2.07±0.84 | 2.15±0.95 | 0.56±0.07 | 2.79±1.00 | 3.23±1.46 | 0.44±0.06 | 0.82 | 3.63 | 0.90 |
| 40 | 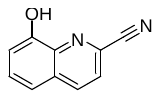 | 0.87±0.13 | 0.91±0.13 | 0.80±0.03 | 3.85±1.48 | 5.03±2.05 | 0.62±0.07 | 0.97 | 1.03 | 0.75 |

| **Table S6.** The Overall RMSD, Pharm RMSD, Volume Overlap of PROTAC predictions (AF3 and Boltz-2) & the confidence metrics of AF3 and Boltz-2 | | | | | | | | | | |
| --- | --- | --- | --- | --- | --- | --- | --- | --- | --- | --- |
| S. No. | Structure | AF3 (2021. Sep. 30) | | | Boltz-2 (2023. Jun.1) | | | AF3 | | Boltz-2 |
|  |  | Overall RMSD | Pharm RMSD | Volume Overlap | Overall RMSD | Pharm RMSD | Volume Overlap | ipTM | minPAE (A) | ipTM |
|  |  | Mean ± SD | Mean ± SD | Mean ± SD | Mean ± SD | Mean ± SD | Mean ± SD |  |  |  |
| 41 | 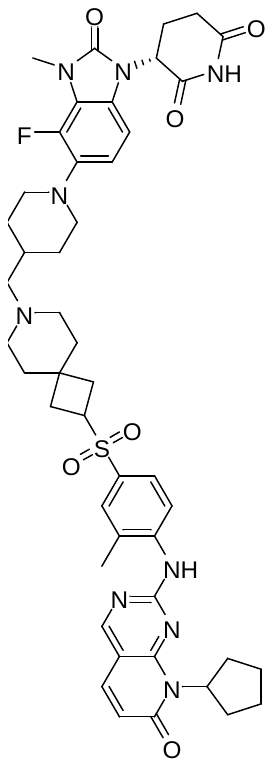 | 2.7795  ±0.4173 | 0.5478  ±0.1677 | 0.57  ±0.0592 | 6.4892  ±1.3054 | 0.3898  ±0.0374 | 0.4154  ±0.0442 | 0.73 | 1.34 | 0.76 |
|  |  | 1.8338  ±0.1533 | 0.9619  ±0.5516 | 0.7089  ±0.0369 | 5.4867  ±0.8856 | 0.5703  ±0.0842 | 0.5529  ±0.039 | 0.73 | 1.20 | 0.76 |
| 42 | 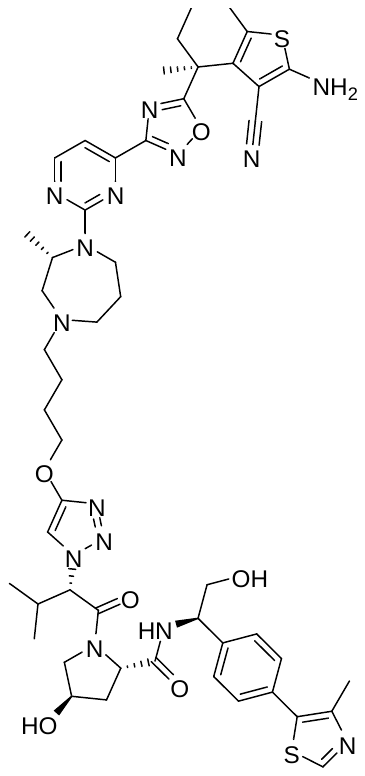 | 27.8084  ±5.9296 | 21.0007  ±8.7453 | 0±0 | 27.5953  ±1.1316 | 15.4861  ±0.2004 | 0±0 | 0.59 | 1.07, 3.19, 2.19, 10.66 | 0.82 |
|  |  | 8.4122  ±2.0493 | 0.7357  ±0.1315 | 0.5208  ±0.0383 | 4.177  ±0.9969 | 0.6733  ±0.2331 | 0.5837  ±0.0489 | 0.59 | 21.21, 20.16, 21.39, 0.92 | 0.82 |
| 30 | 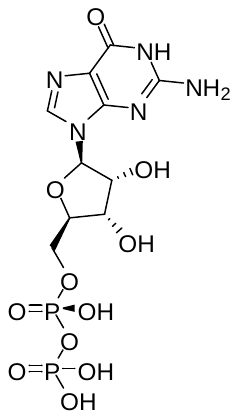 | 1.04±0.63 | 0.83±1.18 | 0.88±0.01 | 0.75±0.01 | 1.57±0.02 | 0.93±0.01 | 0.92 | 0.77 | 0.98 |
|  | 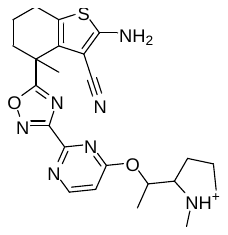 | 15.74±0.19 | 14.23±0.21 | 0.00±0.00 | 1.01±0.54 | 0.45±0.16 | 0.85±0.08 | 0.92 | 1.72 | 0.98 |
| 43 | 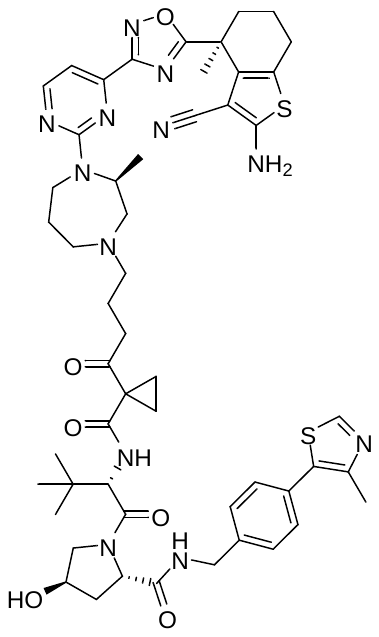 | 23.23±2.65 | 20.13±1.24 | 0.00±0.00 | 28.21±0.92 | 18.80±0.30 | 0.00±0.00 | 0.87 | 1.82 | 0.89 |
|  |  | 7.09±1.09 | 0.39±0.07 | 0.51±0.03 | 3.86±0.71 | 0.42±0.05 | 0.57±0.05 | 0.87 | 0.94 | 0.89 |
| 44 | 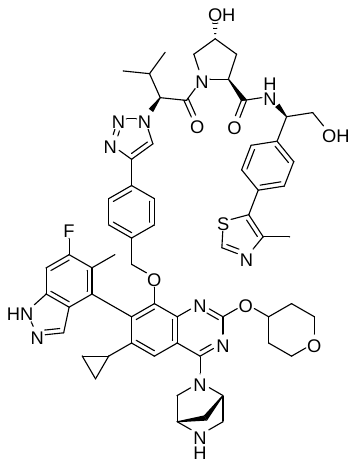 | 28.47±5.11 | 21.54±5.06 | 0.00±0.00 | 5.87±1.70 | 0.96±0.28 | 0.50±0.07 | 0.66 | 9.73 | 0.88 |
|  |  | 6.57±0.36 | 8.84±0.31 | 0.58±0.02 | 4.42±0.88 | 0.62±0.13 | 0.57±0.02 | 0.66 | 1.04 | 0.88 |

| **Table S7.** The Overall RMSD, Pharm RMSD, Volume Overlap of molecular glue predictions (AF3 and Boltz-2) & the confidence metrics of AF3 and Boltz-2 | | | | | | | | | | |
| --- | --- | --- | --- | --- | --- | --- | --- | --- | --- | --- |
| S. No. | Structure | AF3 (2021. Sep. 30) | | | Boltz-2 (2023. Jun.1) | | | AF3 | | Boltz-2 |
|  |  | Overall RMSD | Pharm RMSD | Volume Overlap | Overall RMSD | Pharm RMSD | Volume Overlap | ipTM | minPAE (A) | ipTM |
|  |  | Mean ± SD | Mean ± SD | Mean ± SD | Mean ± SD | Mean ± SD | Mean ± SD |  |  |  |
| 45 | 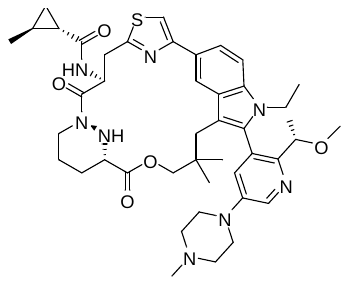 | 18.71±3.44 | 16.00±5.26 | 0.04±0.06 | 36.10±13.71 | 34.22±14.36 | 0.00±0.01 | 0.51 | 10.62 | 0.56 |
|  |  | 4.24±1.86 | 0.56±0.15 | 0.53±0.06 | 3.30±0.63 | 0.58±0.08 | 0.56±0.05 | 0.51 | 1.53 | 0.56 |
| 46 | 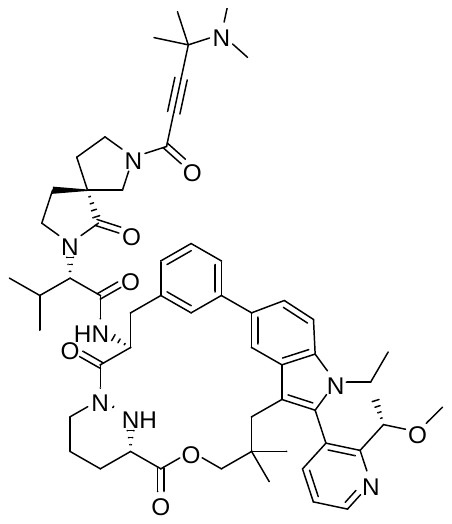 | 3.86±0.30 | 5.76±0.70 | 0.56±0.01 | 3.90±0.49 | 5.50±1.31 | 0.58±0.03 | 0.70 | 0.86, 3.82 | 0.66 |
|  |  | 7.21±1.94 | 8.65±2.61 | 0.36±0.07 | 12.79±5.78 | 7.05±3.02 | 0.20±0.13 | 0.70 | 1.56, 0.88 | 0.66 |
| 47 | 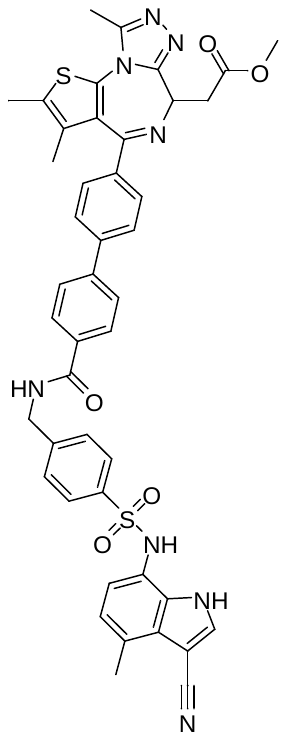 | 32.74±6.50 | 34.40±6.80 | 0.00±0.00 | 29.89±7.19 | 28.62±6.71 | 0.01±0.02 | 0.29 | 21.57 | 0.52 |
| 48 | 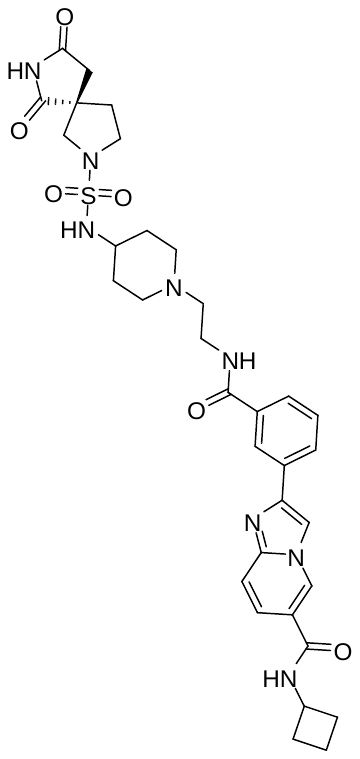 | 9.66±0.37 | 1.76±0.37 | 0.37±0.01 | 8.21±0.75 | 2.07±0.40 | 0.31±0.02 | 0.56 | 1.21 | 0.73 |
|  |  | 14.03±0.57 | 3.66±0.12 | 0.37±0.02 | 20.14±2.08 | 10.81±0.48 | 0.23±0.01 | 0.56 | 2.45 | 0.73 |
| 49 | 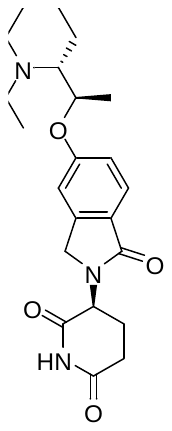 | 5.19±0.73 | 5.35±0.72 | 0.13±0.07 | 141.68±0.28 | 135.62±0.32 | 0.07±0.03 | 0.91 | 4.73 | 0.92 |

| **Table S8.** The Overall RMSD, Pharm RMSD, Volume Overlap of covalent inhibitor predictions (AF3 and Boltz-2) & the confidence metrics of AF3 and Boltz-2 | | | | | | | | | | |
| --- | --- | --- | --- | --- | --- | --- | --- | --- | --- | --- |
| S. No. | Structure | AF3 (2021. Sep. 30) | | | Boltz-2 (2023. Jun.1) | | | AF3 | | Boltz-2 |
|  |  | Overall RMSD | Pharm RMSD | Volume Overlap | Overall RMSD | Pharm RMSD | Volume Overlap | ipTM | minPAE(A) | ipTM |
|  |  | Mean ± SD | Mean ± SD | Mean ± SD | Mean ± SD | Mean ± SD | Mean ± SD |  |  |  |
| 50 | 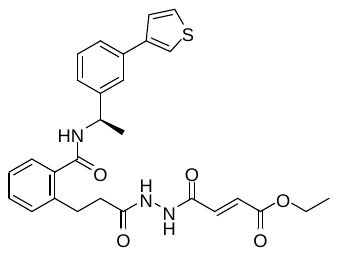 | 3.06±0.05 | 0.84±0.05 | 0.75±0.02 | 2.56±0.86 | 0.53±0.06 | 0.76±0.04 | 0.87 | 0.83 | 0.98 |
| 51 | 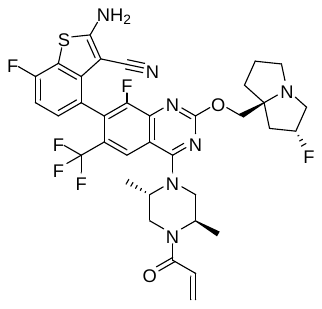 | 1.13±0.17 | 0.50±0.07 | 0.88±0.01 | 10.17±0.08 | 15.68±0.12 | 0.53±0.01 | 0.97 | 0.79 | 0.97 |
| 52 | 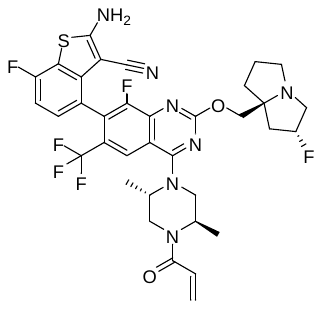 | 1.22±0.14 | 0.49±0.08 | 0.85±0.01 | 1.66±0.05 | 2.38±0.03 | 0.80±0.01 | 0.95 | 0.80 | 0.98 |
| 53 | 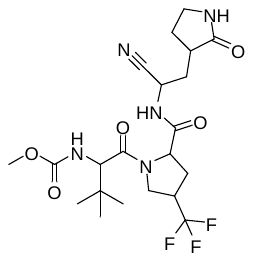 | 0.87±0.19 | 0.35±0.05 | 0.84±0.01 | 0.43±0.07 | 0.30±0.04 | 0.80±0.01 | 0.98 | 0.79 | 0.99 |
| 54 | 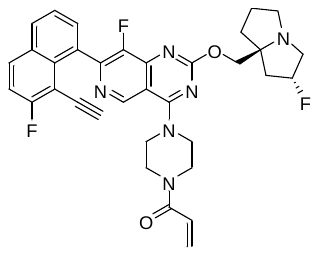 | 0.90±0.08 | 0.11±0.05 | 0.88±0.00 | 11.23±0.04 | 19.45±0.07 | 0.49±0.01 | 0.98 | 0.79 | 0.98 |
| 27 | 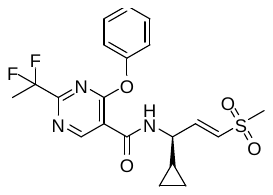 | 9.93±3.50 | 9.05±3.36 | 0.16±0.15 | 11.50±0.88 | 9.99±0.93 | 0.08±0.01 | 0.85 | 1.87 | 0.93 |
| 46 | 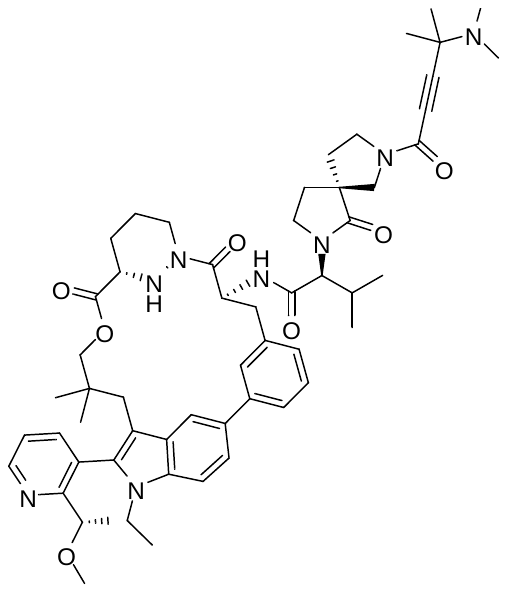 | 3.86±0.30 | 5.76±0.70 | 0.56±0.01 | 3.90±0.49 | 5.50±1.31 | 0.58±0.03 | 0.70 | 0.86, 3.82 | 0.66 |
|  |  | 7.21±1.94 | 8.65±2.61 | 0.36±0.07 | 12.79±5.78 | 7.05±3.02 | 0.20±0.13 | 0.70 | 1.56, 0.88 | 0.66 |

| **Table S9.** The Overall RMSD, Pharm RMSD, Volume Overlap of membrane proteins predictions (AF3 and Boltz-2) & the confidence metrics of AF3 and Boltz-2 | | | | | | | | | | |
| --- | --- | --- | --- | --- | --- | --- | --- | --- | --- | --- |
| S. No. | Structure | AF3 (2021. Sep. 30) | | | Boltz-2 (2023. Jun.1) | | | AF3 | | Boltz-2 |
|  |  | Overall RMSD | Pharm RMSD | Volume Overlap | Overall RMSD | Pharm RMSD | Volume Overlap | ipTM | minPAE(A) | ipTM |
|  |  | Mean ± SD | Mean ± SD | Mean ± SD | Mean ± SD | Mean ± SD | Mean ± SD |  |  |  |
| 55 | 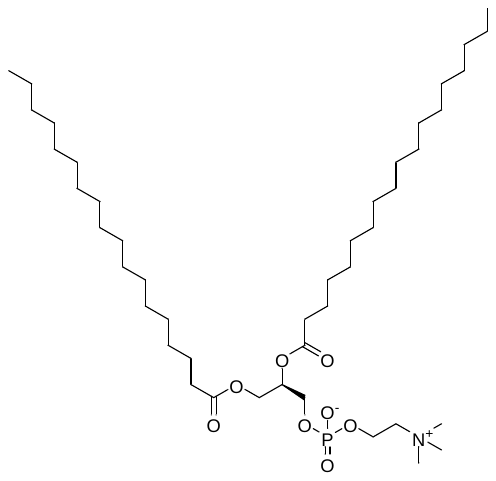 | 21.34±12.0 | 20.07±12.48 | 0.01±0.03 | 51.68±3.33 | 51.95±2.67 | 0.00±0.00 | 0.27 | 16.36, 17.88 | 0.39 |
| 56 | 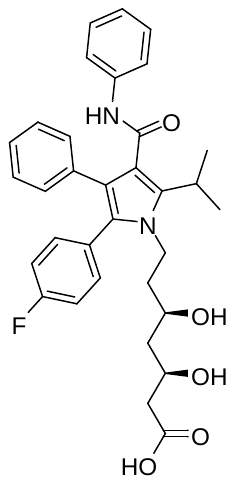 | 9.84±2.94 | 8.72±3.05 | 0.15±0.13 | 7.66±1.19 | 6.50±1.17 | 0.30±0.03 | 0.33 | 4.30, 4.30 | 0.57 |
| 57 | 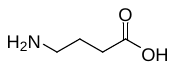 | 4.02±0.71 | 4.85±0.75 | 0.34±0.12 | 3.41±0.12 | 4.28±0.11 | 0.42±0.02 | 0.90 | 2.54 | 0.98 |

| **Table S10.** The Overall RMSD, Pharm RMSD, Volume Overlap of RNA predictions (AF3 and Boltz-2) & the confidence metrics of AF3 and Boltz-2 | | | | | | | | | | |
| --- | --- | --- | --- | --- | --- | --- | --- | --- | --- | --- |
| S. No. | Structure | AF3(2021. Sep. 30) | | | Boltz-2 (2023. Jun. 1) | | | AF3 | | Boltz-2 |
|  |  | Overall RMSD | Pharm RMSD | Volume Overlap | Overall RMSD | Pharm RMSD | Volume Overlap | ipTM | minPAE(A) | ipTM |
|  |  | Mean ± SD | Mean ± SD | Mean ± SD | Mean ± SD | Mean ± SD | Mean ± SD |  |  |  |
| 58 | 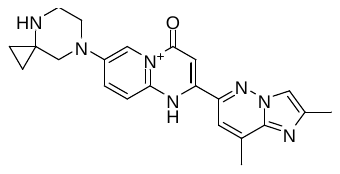 | 1.73±0.74 | 1.61±1.72 | 0.58±0.06 | 8.51±0.26 | 4.86±1.36 | 0.31±0.02 | 0.78 | 2.38 | 0.93 |
| 59 | 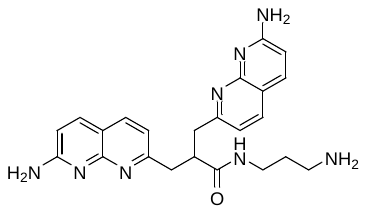 | 16.73±1.44 | 16.21±1.55 | 0.00±0.00 | 19.09±0.24 | 20.74±0.20 | 0.00±0.00 | 0.35 | 3.78, 3.58, 9.02, 8.80 | 0.70 |
| 60 | 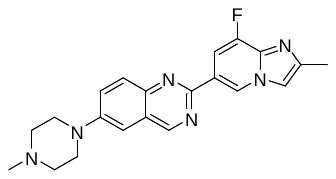 | 2.35±0.11 | 1.35±0.39 | 0.48±0.03 | 8.2±0.42 | 1.75±0.29 | 0.23±0.01 | 0.80 | 1.85, 1.98 | 0.94 |
| 61 | 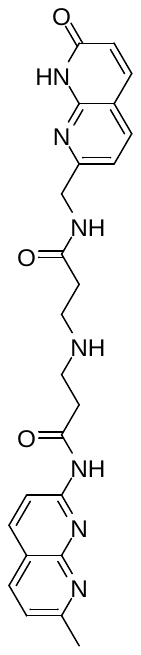 | 10.88±0.36 | 9.47±0.55 | 0.27±0.04 | 6.61±2.60 | 5.84±2.54 | 0.56±0.05 | 0.59 | 2.08 | 0.80 |
| 62 | 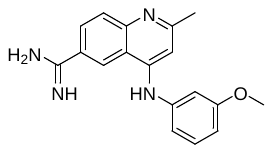 | 5.78±1.88 | 4.57±1.86 | 0.37±0.05 | 6.82±0.52 | 4.09±0.63 | 0.46±0.09 | 0.43 | 2.70, 2.68 | 0.72 |
| 63 | 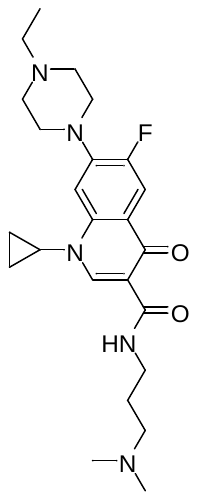 | 7.69±1.98 | 3.45±0.60 | 0.26±0.10 | 2.53±0.17 | 1.60±0.16 | 0.51±0.05 | 0.43 | 2.61 | 0.83 |

| **Table S11.** The Overall RMSD, Pharm RMSD, Volume Overlap of fragment predictions (AF3 and Boltz-2) & the confidence metrics of AF3 and Boltz-2 | | | | | | | | | | |
| --- | --- | --- | --- | --- | --- | --- | --- | --- | --- | --- |
| S. No. | Structure | AF3 (2021. Sep. 30) | | | Boltz-2 (2023. Jun. 1) | | | AF3 | | Boltz-2 |
|  |  | Overall RMSD | Pharm RMSD | Volume Overlap | Overall RMSD | Pharm RMSD | Volume Overlap | ipTM | minPAE (A) | ipTM |
|  |  | Mean ± SD | Mean ± SD | Mean ± SD | Mean ± SD | Mean ± SD | Mean ± SD |  |  |  |
| 64 | 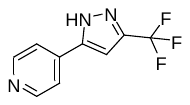 | 10.75±7.85 | 11.12±8.04 | 0.07±0.16 | 4.62±0.25 | 3.81±0.29 | 0.35±0.02 | 0.95 | 12.92, 14.03 | 0.95 |
| 65 | 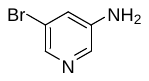 | 15.53±10.58 | 15.77±10.46 | 0.00±0.00 | 7.26±0.18 | 7.91±0.18 | 0.01±0.01 | 0.96 | 11.63, 13.35 | 0.41 |
| 66 | 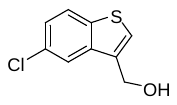 | 12.14±12.21 | 12.60±11.97 | 0.35±0.30 | 18.34±19.64 | 19.00±19.67 | 0.27±0.25 | 0.96 | 12.07, 12.25 | 0.24 |
| 67 | 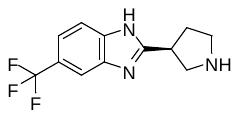 | 5.70±1.41 | 6.86±2.03 | 0.32±0.07 | 6.50±0.04 | 8.04±0.05 | 0.24±0.01 | 0.95 | 8.17, 9,57 | 0.92 |
| 68 | 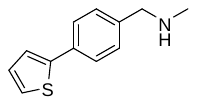 | 4.74±6.41 | 4.49±5.92 | 0.18±0.07 | 9.43±13.92 | 10.21±14.00 | 0.15±0.09 | 0.95 | 15.08, 13.40 | 0.25 |
| 69 | 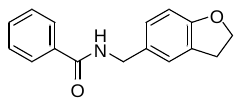 | 22.85±17.52 | 22.46±18.01 | 0.12±0.15 | 7.00±1.86 | 7.84±2.38 | 0.36±0.03 | 0.94 | 10.85, 10.70 | 0.45 |
| 70 | 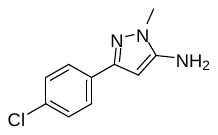 | 8.99±12.88 | 9.43±12.87 | 0.39±0.24 | 18.12±18.09 | 20.24±18.74 | 0.17±0.15 | 0.95 | 13.02, 11.17 | 0.23 |
| 71 | 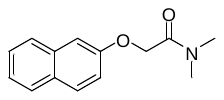 | 33.03±4.10 | 30.81±3.42 | 0.00±0.00 | 28.45±8.59 | 25.68±9.73 | 0.00±0.00 | 0.96 | 11.88, 12.52 | 0.21 |
| 72 | 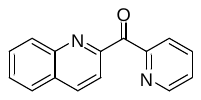 | 32.23±3.42 | 31.18±3.63 | 0.00±0.00 | 32.17±0.48 | 29.78±0.85 | 0.00±0.00 | 0.94 | 11.44, 12.30 | 0.37 |
| 36 | 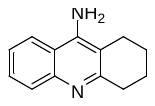 | 4.11±1.41 | 4.67±0.08 | 0.77±0.03 | 2.04±1.51 | 1.43±0.93 | 0.79±0.04 | 0.93 | 1.40 | 0.92 |
| 37 | 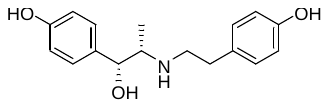 | 7.89±0.18 | 9.81±0.26 | 0.42±0.02 | 6.99±1.56 | 3.14±2.07 | 0.41±0.04 | 0.94 | 1.48 | 0.91 |
| 38 | 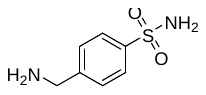 | 2.01±0.20 | 2.42±0.30 | 0.59±0.04 | 1.50±1.12 | 1.96±1.51 | 0.65±0.11 | 0.86 | 3.25 | 0.92 |
| 39 | 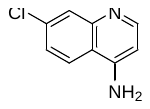 | 2.07±0.84 | 2.15±0.95 | 0.56±0.07 | 2.79±1.00 | 3.23±1.46 | 0.44±0.06 | 0.82 | 3.63 | 0.90 |
| 40 | 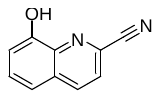 | 0.87±0.13 | 0.91±0.13 | 0.80±0.03 | 3.85±1.48 | 5.03±2.05 | 0.62±0.07 | 0.97 | 1.03 | 0.75 |
| 73 | 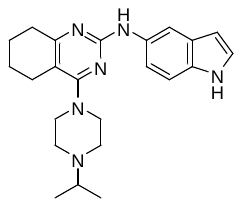 | 1.55±0.25 | 1.03±0.24 | 0.76±0.03 | 5.1±2.20 | 5.47±3.00 | 0.62±0.05 | 0.97 | 0.90 | 0.95 |

| **Table S12.** The Overall RMSD, Pharm RMSD, Volume Overlap of E3 ligase predictions (AF3 and Boltz-2) & the confidence metrics of AF3 and Boltz-2 | | | | | | | | | | |
| --- | --- | --- | --- | --- | --- | --- | --- | --- | --- | --- |
| S. No. | Structure | AF3 (2021. Sep. 30) | | | Boltz-2 (2023. Jun. 1) | | | AF3 | | Boltz-2 |
|  |  | Overall RMSD | Pharm RMSD | Volume Overlap | Overall RMSD | Pharm RMSD | Volume Overlap | ipTM | minPAE（A) | ipTM |
|  |  | Mean ± SD | Mean ± SD | Mean ± SD | Mean ± SD | Mean ± SD | Mean ± SD |  |  |  |
| 74 | 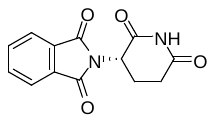 | 1.49±0.73 | 0.28±0.05 | 0.88±0.02 | 1.86±0.55 | 0.21±0.09 | 0.91±0.03 | 0.96 | 0.91 | 0.99 |
| 75 | 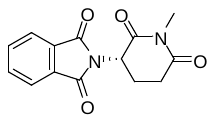 | 1.66±0.68 | 1.10±0.49 | 0.74±0.09 | 2.01±0.72 | 1.68±1.11 | 0.85±0.05 | 0.91 | 1.64 | 0.98 |

| **Table S13**. Experimental activity and Boltz-2-predicted binding affinity for 20 SARS-CoV-2 Mpro inhibitors. | | | | | |
| --- | --- | --- | --- | --- | --- |
| S. No. | PDB ID | Experimental IC_50_ (μM) | Experimental log_10_[IC_50_ (μM)] | Boltz-2 predicted log_10_[IC_50_ (μM)] | minPAE (A) |
| 76 | 7GLV | 0.0197 | -1.705534 | -2.001705 | 1.775 |
| 77 | 7GAW | 0.0368 | -1.434152 | -2.558633 | 0.92 |
| 78 | 7GKS | 0.0516 | -1.287350 | -2.294081 | 1.495 |
| 79 | 7GNT | 0.0744 | -1.128427 | -2.636732 | 0.82 |
| 80 | 7GLG | 0.116 | -0.935542 | -2.418862 | 1.515 |
| 81 | 7GIU | 0.186 | -0.730487 | -1.025081 | 1.035 |
| 82 | 7GM1 | 0.288 | -0.540608 | -2.510219 | 1.51 |
| 83 | 7GNA | 0.452 | -0.344862 | -1.272362 | 1.01 |
| 84 | 7GIX | 0.692 | -0.159894 | -1.450610 | 0.87 |
| 85 | 7GHM | 1.1 | 0.041393 | -2.695688 | 1.095 |
| 86 | 7GG2 | 1.71 | 0.232996 | -1.434742 | 1.525 |
| 87 | 7GMR | 2.67 | 0.426511 | -1.380679 | 1.005 |
| 88 | 7GE3 | 4.22 | 0.625312 | -0.308186 | 1.205 |
| 89 | 7GMP | 6.78 | 0.831230 | -1.415606 | 1.535 |
| 90 | 7GE2 | 10.6 | 1.025306 | -1.059038 | 1.85 |
| 91 | 7GE4 | 16.1 | 1.206826 | -0.258463 | 1.64 |
| 92 | 7GGC | 25.3 | 1.403121 | 0.099318 | 1.2 |
| 93 | 7GFJ | 38.8 | 1.588832 | -0.866213 | 1.14 |
| 94 | 7GCK | 59.9 | 1.777427 | -1.609528 | 1.44 |
| 95 | 7GBV | 96.9 | 1.986324 | 0.156321 | 1.065 |
